# Threshold-awareness in adaptive cancer therapy

**DOI:** 10.1101/2022.06.17.496649

**Authors:** MingYi Wang, Jacob G. Scott, Alexander Vladimirsky

## Abstract

Although adaptive cancer therapy shows promise in integrating evolutionary dynamics into treatment scheduling, the stochastic nature of cancer evolution has seldom been taken into account. Various sources of random perturbations can impact the evolution of heterogeneous tumors, making performance metrics of any treatment policy random as well. In this paper, we propose an efficient method for selecting optimal adaptive treatment policies under randomly evolving tumor dynamics. The goal is to improve the cumulative “cost” of treatment, a combination of the total amount of drugs used and the total treatment time. As this cost also becomes random in any stochastic setting, we maximize the probability of reaching the treatment goals (tumor stabilization or eradication) without exceeding a pre-specified threshold (or a “budget”). We use a novel Stochastic Optimal Control formulation and Dynamic Programming to find such “threshold-aware” optimal treatment policies. Our approach enables an efficient algorithm to compute these policies for a range of threshold values simultaneously. Compared to treatment plans shown to be optimal in a deterministic setting, the new “threshold-aware” policies significantly improve the chances of the therapy succeeding under the budget, which is correlated with a lower general drug usage. We illustrate this method using two specific examples, but our approach is far more general and provides a new tool for optimizing adaptive therapies based on a broad range of stochastic cancer models.

**Author Summary:** Tumor heterogeneities provide an opportunity to improve therapies by leveraging complex (often competitive) interactions of different types of cancer cells. These interactions are usually stochastic due to both individual cell differences and random events affecting the patient as a whole. The new generation of cancer models strive to account for this inherent stochasticity, and *adaptive* treatment plans need to reflect it as well. In optimizing such treatment, the most common approach is to maximize the probability of eventually stabilizing or eradicating the tumor. In this paper, we consider a more nuanced version of success, maximizing the probability of reaching these therapy goals before the cumulative burden from the disease and treatment exceed a chosen threshold. Importantly, our method allows computing such optimal treatment plans efficiently and for a range of thresholds at once. If used on a high-fidelity personalized model, our general approach could potentially be used by clinicians to choose the most suitable threshold after a detailed discussion of a specific patient’s goals (e.g., to include the trade-offs between toxicity and quality of life).

## 1 Introduction

Optimizing the schedule and composition of drug therapies for cancer patients is an important and active research area, with mathematical tools often employed to improve the outcomes and reduce the negative side effects. Tumor heterogeneity is increasingly viewed as a key aspect that can be leveraged to improve therapies through the use of optimal control theory [55]. Most researchers using this perspective focus on deterministic models of tumor evolution, with typical optimization objectives of maximizing the survival time [46], minimizing the tumor size [6], or minimizing the time until the tumor size is stabilized [7]. In models that address the stochasticity in tumor evolution, a typical optimization goal is to find treatment policies that maximize the likelihood of patient’s eventual cure (e.g., [10, 13, 37]) or minimize the likelihood of negative events (e.g., metastasis) after specified time [19]. However, this ignores the need for more nuanced treatment policies that maximize the likelihood of different levels of success – e.g., the probability of reaching remission or tumor stabilization without exceeding the specified amount of drugs and/or the specified treatment duration. The primary goal of this paper is to introduce a rigorous and computationally efficient approach that addresses such challenging objectives in stochastic cancer models.

The advent of personalized medicine in cancer has changed the way we think about therapy for patients whose tumors have actionable mutations. This has been a game changer for some patients, drastically increasing life spans, reducing toxicity and improving quality of life. Frustratingly, however, this population of patients is still small; it was estimated in 2020 that only *≈* 5% of patients benefit from these targeted therapies [30]. Further, despite the many advantages of personalized therapies, they rarely, if ever, lead to a complete cure since tumors develop resistance through the process of Darwinian evolution [24]. In response to this realization, a new approach called “evolutionary therapy” seeks to use the evolutionary dynamics of diseases to alter therapeutic schedules and drug choices. Through a combination of mathematical and experimental modeling, investigators have worked to understand a range of theoretical questions of practical importance. E.g., how does the emergence of resistance to one drug affect the sensitivity to another? Do heterogeneous (phenotypically or genotypically mixed) populations within tumors respond to drugs differently depending on their current state? The insights gained in these investigations have already led to progress in rational drug ordering/cycling for bacterial infections [34, 45, 50, 51] as well as for a number of cancers [14, 65]. In the study of therapeutic scheduling, adaptive therapy, which uses mathematical tools from Evolutionary Game Theory (EGT), has shown promise not only in theory [23], but also in a phase 2 trial for men with metastatic prostate cancer [63]. Experimentally, there have been confirmations of EGT principles *in vivo*[17] as well as more quantitatively focused assay development *in vitro* [38], and observations of game interactions using these methods [18]. There are also many other models capturing the competition within heterogeneous tumors without using game-theoretic derivations; e.g., [6, 9, 26, 42]. The majority of theoretical work in this space has focused on optimization of different drug regimens for *deterministic* models of cancer evolution [11, 12, 25, 60, 61]. In contrast, our goal here is to provide efficient computational tools for nuanced therapeutic scheduling in cancer models that directly account for stochastic perturbations.

Cancers (and other populations of living things) are comprised of individual cells (or organisms) with their own behaviours and evolutionary histories. Stochastic phenomena are ubiquitous in their interactions and life histories. These include individual genetic differences, fate transitions [27], varying reactions to drugs [41], differences in signalling, and small-scale variations in the tumor environment. Many of these are instances of *demographic stochasticity* [43], which often can be “averaged-out” when dealing with a sufficiently large population. Indeed, this notion is crucial for any description of tumor heterogeneity through splitting the cells into sub-populations. Such splitting is natural if the mutation-selection balance is tuned so that only closely related genotypes, encoding the same phenotype, will stably exist. These groups are also referred to as quasispecies [44, 62] and exist as distributions around a central genotype, with all cells in the group behaving in a similar manner despite random birth/death events [16, 43] and small within-the-group genetic heterogeneities [35]. In contrast, our focus here is on *environmental stochasticity*, which cannot be ignored even in large populations since it describes random events that simultaneously affect the entire groups. Such perturbations are typically external [16, 43]; e.g., for cancer they might result from therapy-unrelated drugs or from frequent small changes in the host’s physiology. Of course, any such event will also cause varying responses of individual cells within each subpopulation; so, our use of the term “environmental stochasticity” should be interpreted as direct modeling of subpopulation-averaged responses to such system-wide perturbations.

Modeling such perturbations in continuous-time usually results in *Stochastic Differential Equations* (SDEs) [2, 4], whose behavior can be optimized using Stochastic Optimal Control Theory [20]. The latter provides a mathematical framework for handling sequential decision making (e.g., how much drug to administer at each point in time) under random perturbations (e.g., stochastic changes in respective fitness of competing subpopulations of cancer cells). Any fixed treatment strategy will result in a random tumor-evolutionary trajectory and a random cumulative “cost” (e.g., cumulative amount of drugs used, or time to recovery, or a combination of these two metrics). The key idea of *Dynamic Programming* (DP) is to pose equations for the cumulative cost of the optimal strategy and to recover that strategy *in feedback form*: i.e., decisions about the dose and duration of therapy are frequently re-evaluated based on the current state of the tumor instead of selecting a fixed time-dependent treatment schedule in advance. This idea is applicable across a wide range of cancer models and therapy types, including those intended to stabilize the tumor and those aiming to eradicate it. We follow this approach here, but with an important caveat: instead of selecting an *on-average optimal* strategy (e.g., the one which minimizes the expected cost of treatment) as would be usual in stochastic DP, we select a strategy maximizing the probability of some desirable outcome (e.g., reaching the goals of the therapy *without exceeding a specific cost threshold*). The resulting risk-aware (or, more precisely, “threshold-aware”) policies are designed to be adaptive, adjusting the treatment plan along the way based on the responsiveness of tumor to drugs already used (and the cost already incurred) so far. In contrast to standard methods of constrained stochastic optimal control, our approach makes it easy to compute such threshold-aware policies for a range of thresholds simultaneously.

As is often the case, there remains a significant gap between simplified mathematical models and clinical applications. Much work remains in refining and calibrating EGT models, and also in measuring different aspects of biological stochasticity. But once high-fidelity personalized models become available, our general approach could potentially be used by clinicians to choose the most suitable threshold after a detailed discussion of a specific patient’s goals (to include the trade-offs between toxicity and quality of life, for example).

## 2 Methods and Models

To emphasize the broad applicability of our “risk-aware” adaptive therapy optimization approach, we first describe it for a fairly generic cancer model. Two specific examples are then studied in detail in §2.2 and §2.3.

### 2.1 Traditional and risk-aware control in drug therapy optimization

We note that most of the literature on dynamic programming in cancer models starts with positing a specific known/fixed treatment horizon *T*, with the success or failure of therapy assessed after that time (or earlier, in case of the modeled patient’s death). This makes it easier to use the standard equations and algorithms of “finite-horizon” optimal control theory. But such a pre-determined *T* is not well-aligned with the notion of adaptive therapies. Instead, we adopt the *indefinite-horizon* framework, in which the process terminates as soon as the tumor’s state satisfies some predefined conditions, with the terminal time *T* thus dependent on the chosen treatment policy. We use this framework in all of the control approaches described below, even though many of them have direct finite-horizon analogs as well.

We begin by describing several “traditional” optimal control formulations, followed by the threshold-aware version, which addresses some of their shortcomings in cancer applications. Starting with the deterministic setting summarized in Box 1, we use ***x***(*t*) ∈ ℝ^*n*^ to encode the time-dependent state of a tumor (e.g., this could be the size or the relative abundance of *n* different sub-types of cancer cells). Tumor dynamics are modeled by a function ***f***, which takes as inputs both the current state ***x***(*t*) and the current control, the “therapy intensity” ***d***(*t*). In models with a single drug, this is just a scalar *d*(*t*) ∈ [0, *d*_max_] indicating the current rate of that drug’s delivery, where *d*_max_ encodes the Maximum Tolerated Dose (MTD). But the same framework can be also used for multiple drugs, with a separate upper bound specified for each element of ***d***(*t*). Given an initial tumor state ***x***(0) = ***ξ***, a successful therapy aims to drive the tumor state to a set Δ_succ_ while ensuring that the set Δ_fail_ is avoided^I^. If the therapy accomplishes this, its overall “cost” 𝒥 is assessed by integrating some running cost *K* along the “trajectory” from ***ξ*** to Δ = Δ_succ_ ∪ Δ_fail_ and adding the “terminal cost” *g* depending on its final state. E.g., *g* might be defined as +∞ on Δ_fail_ to make such outcomes unacceptable. The running cost *K* depends on the changing state of the tumor and the current drug usage levels and can be used to model the direct impact on the patient of the tumor size and composition as well as the side effects of the therapy. The key idea of dynamic programming [3] is to define a *value function u*(***ξ***) encoding the minimal overall cost for each specific initial tumor state and to show that this *u* must satisfy a stationary Hamilton-Jacobi-Bellman (HJB) equation (2.4). Once that partial differential equation (PDE) is solved numerically, the globally optimal rate of treatment can be obtained in *feedback form* for all cancer states (i.e., ***d*** = ***d***_⋆_(***ξ***)), which makes it suitable for the adaptive therapy framework. If *K* is chosen so that the overall cost of a successful therapy 𝒥 reflects a weighted sum of the total therapy duration and the cumulative use of each drug, the weights can be adjusted to reflect the relative importance of these optimization criteria. In this case, if ***f*** is also a linear function of ***d***, it is easy to show that the optimal treatment policy ***d***_⋆_(***ξ***) is generally *bang-bang*; i.e., for each drug, it prescribes either no usage or the maximal (MTD) usage in every cancer state ***ξ***.

#### Box 1

Problem setup of a typical deterministic optimal cancer-control problem

**Evolutionary dynamics with control on therapy intensity:**

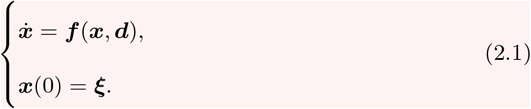

**Process *terminates* as soon as either**

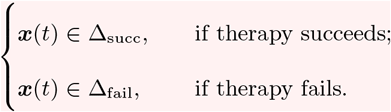

**Definitions and Parameters:**

- ***x*** ∈ ℝ^*n*^, *n*-dimensional cancer state;
- ***d*** : ℝ_+_ → 𝒟 (𝒟 compact), time-dependent intensity of the therapy (control);
- Δ_succ_ ⊂ ℝ^*n*^, success region;
- Δ_fail_ ⊂ ℝ^*n*^, failure region;
- Δ = Δ_succ_ ∪ Δ_fail_, terminal set.

**Total treatment time:**

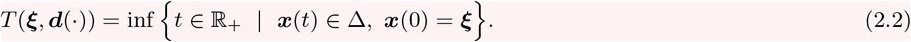

**Treatment cost function:**

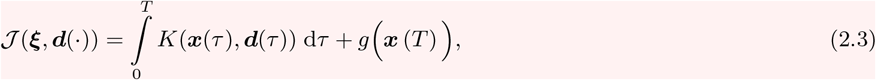

where *T* := *T* (***ξ, d***(·)) is the terminal time, *K*(***x, d***) is the running cost, and the terminal cost is

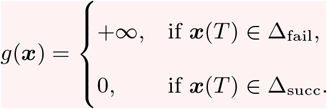

**Value function:** 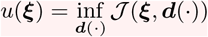 is found by numerically solving a first-order HJB PDE

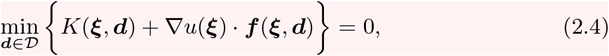

with the boundary condition *u* = *g* on Δ.

See SM §3S.1 for the derivation and §4S.2 for the numerics.

In a generic continuous-time stochastic cancer model (summarized in Box 2), the tumor state ***X***(*t*) becomes a random variable, with the dynamics specified by a Stochastic Differential Equation (SDE) (2.5), which replaces the deterministic Ordinary Differential Equation (ODE) (2.1). The definitions of the *total treatment time T* (***ξ, d***(·)) and the *overall treatment cost 𝒥* (***ξ, d***(·)) remain the same, but both of them become random variables. The standard (*risk-neutral*) approach of stochastic optimal control is to find a feedback-form treatment policy ***d***_⋆_(***ξ***) that minimizes the expected treatment cost 𝔼 [*𝒥*]. As explained in Box 3, the resulting value function satisfies another stationary HJB PDE (2.7). The choice of suitable boundary conditions is more subtle here: setting *g* = +∞ on Δ_fail_ is no longer an option since the probability of entering Δ_fail_ before Δ_succ_ is usually positive under every treatment policy, which would result in 𝒥 = +∞ for every occasional failure and the overall 𝔼 [*𝒥*]= +∞. This makes it necessary to either choose a specific finite “cost” of failure (which can be problematic both for practical and ethical reasons) or switch to an entirely different optimization objective. For example, one can try to simply maximize the probability of fulfilling the therapy goals (i.e., eventually reaching Δ_succ_ while avoiding Δ_fail_) by solving the equation (2.9). But this latter formulation ignores many important practical considerations: e.g., it can easily result in an unreasonably long treatment time or in significant side effects from a prolonged MTD-level drug administration.

#### Box 2

Problem setup of the stochastic optimal control problem

**Stochastic evolution dynamics with control on therapy intensity (a *drift-diffusion* process):**

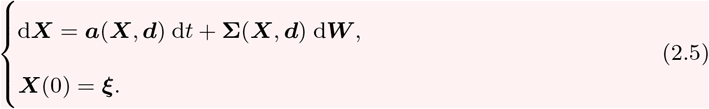

**Process *terminates* as soon as either**

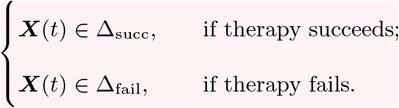

**Definitions and Parameters:**

- ***X*** ∈ ℝ^*n*^, *n*-dimensional cancer state;
- ***W***, standard *m*-dimensional Brownian motion;
- ***a***(***X, d***) ∈ ℝ^*n*^, the drift function;
- **Σ**(***X, d***) ∈ ℝ^*n×m*^, the diffusion function.

**Note:** Definitions of the *total treatment time T* := *T* (***ξ, d***(·)) and the *treatment cost function J* (***ξ, d***(·)) stay the same as in Box 1. But they are now *random variables* as we will replace ***x***(*t*) by ***X***(*t*).

#### Box 3

Standard stochastic dynamic programming approaches

**A risk-neutral (expectation-minimizing) approach [21] :**

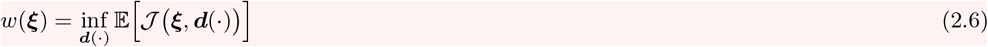

can be found by solving a second-order Hamilton-Jacobi-Bellman (HJB) equation:

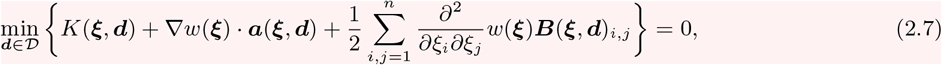

where ***B*** = **ΣΣ**^⊤^ and *w* = *g* on Δ = Δ_succ_ ∪ Δ_fail_.

**Note:** If one uses *g*(***X***(*T*)) = +∞ when therapy fails (i.e., when ***X***(*T*) ∈ Δ_fail_), the diffusion will generally result in *w* = +∞ for most if not all initial tumor configurations outside of Δ_succ_.

**An alternative is to maximize the probability of eventual goal attainment:**

**Value function:**

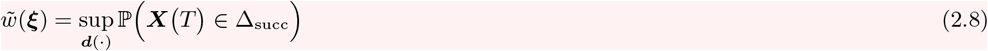

can be found by solving a second-order Hamilton-Jacobi-Bellman (HJB) equation:

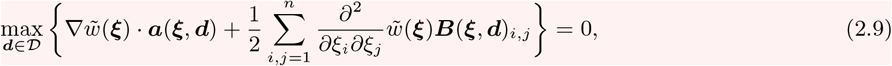

with the boundary condition 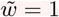on Δ_succ_ and 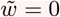 on Δ_fail_.

In contrast, the approach we are pursuing here allows for a more nuanced definition of success (e.g., taking into account the total drug usage, the treatment duration, and/or the cumulative burden from the tumor). Choosing a running cost *K* to reflect the above factors, we define the overall cost as 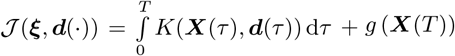, which might be infinite if ***X***(*T*) ∈ Δ_fail_. We then maximize the probability of reaching the policy goals, but constraining the overall cost by some pre-specified threshold 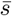. I.e., we need to find an adaptive therapy that maximizes 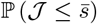.

Our goal is to compute such *threshold-aware* policies efficiently for all starting tumor configurations ***ξ*** and a broad range of threshold levels simultaneously. It is easy to see that here good treatment policies will have to also take into account the cost accumulated so far. This makes it natural to treat our chosen threshold 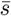 as an *initial cost budget*, tracking the remaining budget *s*(*t*) by solving equation (2.13) in Box 4. The value function can be found by solving the parabolic PDE^II^ (2.11) numerically, and the optimal feedback policy 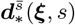 is recovered in the process^III^ for all 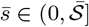.

#### Box 4

Our threshold-aware approach

**Value function:**

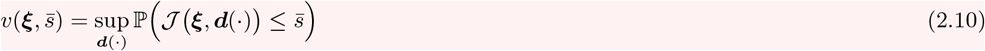

can be found by solving a different second-order HJB equation:

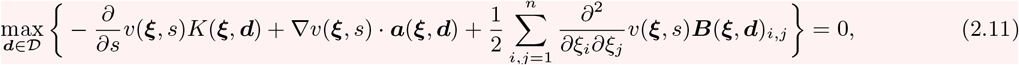

where 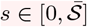. See the detailed derivation in §3S.2 of Supplementary Materials.

The boundary conditions of HJB equation:

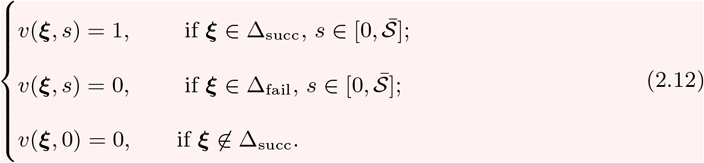

The (random) ODE describing the reduction of budget:

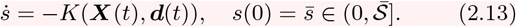

While this threshold-aware framework has important advantages illustrated below, it also brings to the forefront several subtleties avoided in the more traditional stochastic optimal control approaches. First, an adaptive treatment policy optimal for one specific threshold is usually not optimal for another. (The starting budget in (2.13) is important for deciding when to administer drugs.) This would make it necessary for a practitioner to have a detailed discussion with their patient to choose a suitable threshold value before the treatment is started. Second, stochastic perturbations make the outcome random and the budget might run out (*s*(*t*) = 0) under any treatment policy. But this scenario is only a failure in the sense that the overall cost *𝒥* will now definitely exceed the threshold value 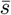. If the patient is still alive (***X***(*τ*) *∉* Δ_fail_, *∀τ* ∈ [0, *t*]) and interested in continuing treatment, one has to make a decision on the new strategy. This can be done either by posing a new threshold for future treatment costs or by switching to an entirely different policy – e.g., either by employing some traditional stochastic optimal control approach (based on equations (2.7) or (2.9)) or by using a deterministic-optimal policy based on equation (2.4). The latter version is used in all stochastic simulations in the following sections.

Informally, threshold-aware policies reflect a tension between two objectives which are often (but not always) in conflict. Maximizing the probability of treatment attaining its primary goals (e.g., tumor stabilization or eradication) is balanced against reducing the cost (a combination of tumor and treatment burdens) suffered along the way. The former is optimized but only over the scenarios where the latter stays below the prescribed threshold. We close this subsection by highlighting connections of our approach to general multiobjective optimal control and optimal control with integral constraints.

In deterministic optimal control theory, the idea of treating some version of cumulative cost as an additional state variable is well-known. But the resulting ODE systems are typically treated within the framework of *Pontryagin’s Maximum Principle* (PMP) [54], which has an important advantage (its suitability for high-dimensional problems) but also a number of serious drawbacks: the fact that policies are not recovered in feedback form, the fact that these policies are generally not guaranteed to be *globally* optimal, occasional difficulties in ensuring the convergence of numerical methods needed to find such policies, and challenges in handling non-trivial state constraints. In cancer literature, this PMP-based approach has been used to impose “isoperimetric constraints” on the amount of administered chemotherapy [66] or imunotherapy [29]. In addition to the issues listed above, we note that the suitability of equality (isoperimetric) constraints is not obvious in many cancer applications. Indeed, the fact that a less aggressive treatment may in some cases improve the outcomes is one of the main reasons for the interest in adaptive therapies. Thus, insisting that all available drugs must be used is hard to justify, and inequality constraints (e.g., imposing an upper bound on the cumulative drug use) seem much more reasonable.

The first dynamic programming (HJB-based) formulation for handling such constraints in general deterministic control problems was developed in [40]. It circumvents all these PMP-associated difficulties with an added benefit of finding globally optimal policies for a range of inequality constraint levels simultaneously. The threshold-aware method presented here extends many of the same ideas to a stochastic setting.

### 2.2 Example 1: an EGT-based competition model

To develop our first example, we adopt the base model of cancer evolution proposed by Kaznatcheev et al. in [39], which describes a competition of 3 types of cancer cells. Glycolytic cells (GLY) are anaerobic and produce lactic acid. The other two types are aerobic and benefit from better vasculature, development of which is promoted by production of the VEGF signaling protein. Thus, the VEGF (over)-producing cells (VOP) devote some of their resources to vasculature development, while the remaining aerobic cells are essentially free-riders or *defectors* (DEF) in game-theoretic terminology.

If (*z*_G_(*t*), *z*_D_(*t*), *z*_V_(*t*)) encode the time-dependent sizes of these three cancer types, their dynamics are given by 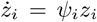, where *i* ∈ {*G, D, V*} and (*ψ*_G_, *ψ*_D_, *ψ*_V_) are the respective type fitnesses. The actual expressions for these *ψ*_*i*_ are derived from the inter-population competition in the usual EGT framework; see Supplementary Materials (SM) §1S.1. This competition of cells in the tumor is modeled as a “public goods” / “club goods” game: VEGF is a “club good” since it benefits only VOP and DEF cells, while the acid generated by GLY is a “public good” for all cancer cells since it is damaging for the surrounding non-cancerous tissue. The base model in [39] assumes that each cell interacts with *n* others nearby. How much it benefits from these interactions depends on its own type and the proportions of different cell types among those nearby cells. Assuming that all participants are drawn uniformly at random from a large well-mixed population, one can derive all fitnesses *ψ*_*i*_ as expected payoffs in this game of (*n* + 1) players. Those expected payoffs will naturally depend on the current subpopulation fractions (or relative abundances) 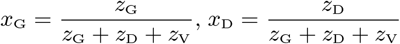, and 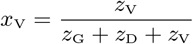. A *Replicator* Ordinary Differential Equation (ODE) [33, 57] is a standard EGT model for predicting the changes in these subpopulation-fractions as a function of time.

In both the original deterministic case and its stochastic extension, it is easier to view the replicator equation as a 2-dimensional system (e.g., by noting that *x*_D_ = 1 *− x*_G_ *− x*_V_). Following [39], we use a slightly different reduction, rewriting everything in terms of the proportion of glycolytic cells in the tumor *p*(*t*) = *x*_G_(*t*) and the proportion of VOP among aerobic cells 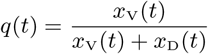. A drug therapy (in this example, affecting the fitness of GLY cells only) is similarly easy to encode by modifying the Replicator ODE; see equation (2.14) in Box 5 and the Supplementary Materials in [39] for the derivation. The goal of the drug therapy here is to drive the GLY fraction *p*(*t*) = *x*_G_(*t*) down^IV^ below a specified “stabilization barrier” *γ*_r_. For a range of parameter values, this model yields periodic behavior of cancer subpopulations: without drugs, *x*_G_(*t*), *x*_D_(*t*), and *x*_V_(*t*) alternate in being dominant in the tumor, with the amplitude of oscillations determined by the initial conditions [39]. This highlights the importance of proper timing in therapies: starting from the same initial tumor composition (*q*_0_, *p*_0_), the same MTD therapy of a fixed duration could lead to either a stabilization (*p*(*t*) falling below *γ*_r_) or a death (*p*(*t*) rising above the specified “failure barrier” *γ*_f_) depending on how long we wait until this therapy starts; see Figure 2 in [39].

This strongly suggests the advantage of *adaptive therapies*, which prescribe the amount of drugs based on continuous or occasional monitoring of (*q*(*t*), *p*(*t*)) or some proxy (non-invasively measured) variables. A natural question is how to optimize such policies to reduce the total amount of drugs used and the total duration of treatment until *p*(*t*) *< γ*_r_. Gluzman et al. have addressed this in [25] using the framework of deterministic optimal control theory [20]. A time-dependent intensity of the therapy *d*(*t*) (ranging from 0 to the MTD level *d*_max_) was chosen to minimize the overall cost of treatment 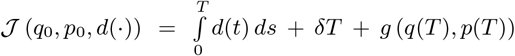, where *T* is the time till stabilization (or failure, if (*q*(*T*), *p*(*T*)) ∈ Δ_fail_ and *g* = +∞) while the value of *δ >* 0 reflects the relative importance of two optimization goals (total drugs vs total time). In the framework of deterministic dynamic programming [3] summarized in Box 1, this corresponds to minimizing the integral of the running cost *K* = *d*(*t*)+*δ*. In [25], the deterministic-optimal policy is obtained in *feedback form* (i.e., *d* = *d*_⋆_(*q, p*)) by numerically solving the Hamilton-Jacobi-Bellman (HJB) PDE (2.4). Figure 1(a) summarizes this policy (showing in yellow the MTD region where *d*_⋆_(*q, p*) = *d*_max_) and illustrates the corresponding trajectory for one specific initial (*q*_0_, *p*_0_).

**Figure 1:**
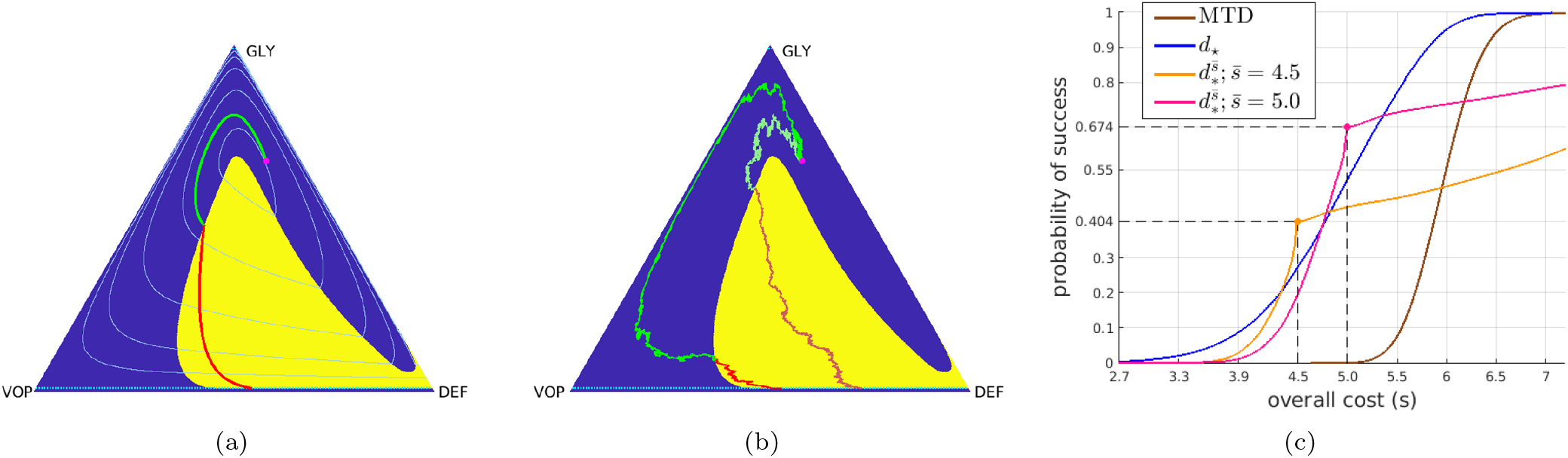
Deterministic-optimal policy in the EGT-model. The (GLY-VOP-DEF) triangle represents all possible relative abundances of respective subpopulations. Since the policy is bang-bang, we show it by using the yellow background where drugs should be used at the MTD rate and the blue background where no drugs should be used at all. Starting from an initial state (*q*_0_, *p*_0_) = (0.26, 0.665) (magenta dot), the subfigures show (a) the optimal trajectory found from the truly deterministically driven system (2.14) with cost 5.13; (b) two representative sample paths generated under the deterministicoptimal policy but subject to stochastic fitness perturbations (the brighter one has a cost of 3.33 while the other has a cost of 6.23); (c) CDFs of the cumulative cost 𝒥 approximated using 10^5^ random simulations. In both (a) & (b), *the green parts of trajectories* correspond to not prescribing drugs and *the red parts of trajectories* correspond to prescribing drugs at the MTD rate. In (a), the level sets of the value function in the deterministic case are shown in *light blue*. In (c), the *blue* curve is the CDF generated with the deterministic-optimal policy *d*_⋆_. Its observed median and mean conditioning on success are 4.95 and 4.91 respectively. The *brown* curve is the CDF generated with the MTD-based therapy, which in this example also maximizes the chances of “budget-unconstrained” tumor stabilization. Its observed median and mean conditioning on success are 5.95 and 5.96 respectively. *Orange* and *pink* curves show the CDFs for two different threshold-aware policies (with 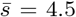 and 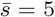 respectively). The large dot on each of them represents the maximized probability of not exceeding the corresponding threshold. The term “threshold-specific advantage” refers to the fact that, at 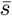, the CDF of 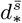 is above the CDFs of all other policies.

**Figure 2:**
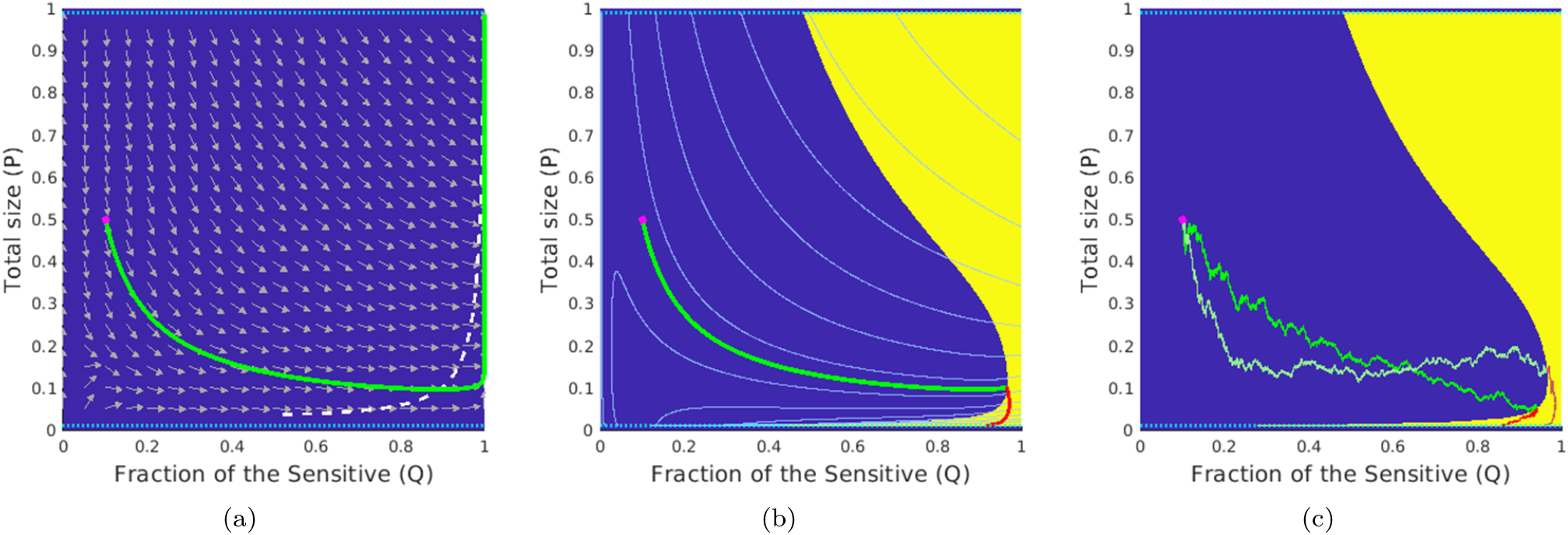
Deterministic-optimal policy in the Sensitive-Resistant model. Starting from an initial state (*q*_0_, *p*_0_) = (0.1, 0.5) (magenta dot), the subfigures show (a) the deterministic trajectory without therapy that ends in the Δ_fail_; (b) the optimal trajectory found from the deterministically driven system (2.1,2.19) with cost 49.30; (c) two representative sample paths generated under the deterministic-optimal policy but subject to a stochastic perturbation in (*g*_S_, *g*_R_) (the brighter one has a cost of 70.45 while the other has a cost of 49.43); In (a), the *white dashed-line* is part of the nullcline where 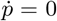 In both (b)&(c), *the green parts of trajectories* correspond to not prescribing drugs and *the red parts of trajectories* correspond to prescribing drugs at the MTD rate. The level sets of the value function *u* in the deterministic case are shown in *light blue*.

#### Box 5

Example 1 (base model adopted from [25, 39])

**The deterministic base model (components for the approach in Box 1):**

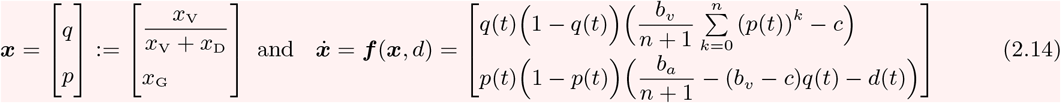

The above reflects the formulas for subpopulation fitnesses (*ψ*_G_, *ψ*_D_, *ψ*_V_); see details in SM § 1S.1.

**The stochastic model (components for the approaches in Boxes 2-4):**

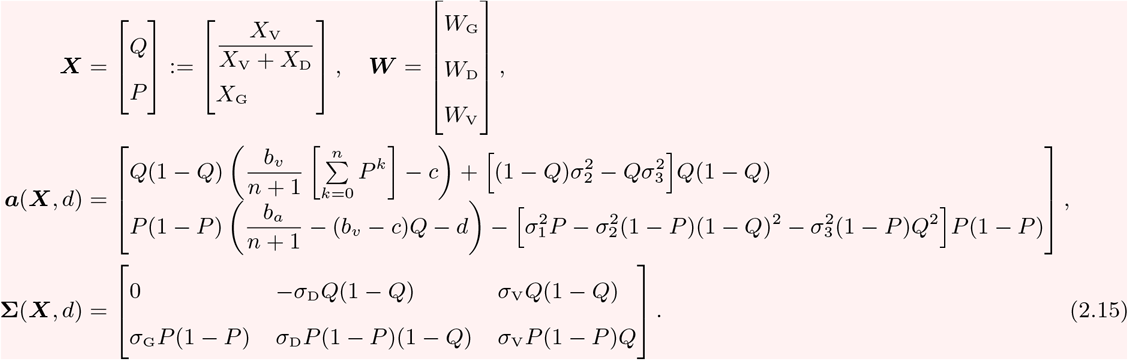

**Definitions and Parameters:**

- *d* : ℝ_+_ → [0, *d*_max_], time-dependent intensity of GLY-targeting therapy;
- Δ_succ_ = {(*q, p*) ∈ [0, 1]^2^| *p < γ*_r_}, success region where *γ*_r_ is the *stabilization barrier*;
- Δ_fail_ = {(*q, p*) ∈ [0, 1]^2^ | *p > γ*_f_}, failure region where *γ*_f_ is the *failure barrier*;
- *K*(***X***, *d*) = *d* + *δ*, running cost function where *δ* is the treatment time penalty;
- 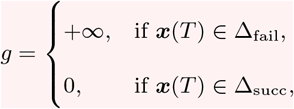 terminal cost;
- ***W*** = (*W*_G_, *W*_D_, *W*_V_), standard 3D Brownian motion for (GLY, DEF, VOP) cells;
- (*σ*_G_, *σ*_D_, *σ*_V_), volatilities for (GLY, DEF, VOP) cells;
- *b*_*a*_, the benefit per unit of acidification;
- *b*_*v*_, the benefit from the oxygen per unit of vascularization;
- *c*, the cost of production of VEGF;
- (*n* + 1), the number of cells in the interaction group.

**Conditions for the heterogeneous regime (coexistence of all cell types):**

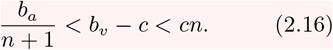

**The optimal policy in feedback form:**

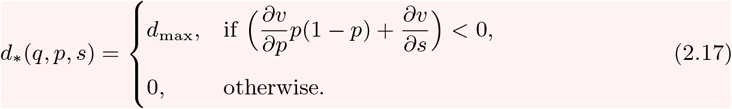

A natural way to introduce stochastic perturbations into this base model is to assume that the rates of sub-population growth/decay are actually random and normally distributed at any instant, with the fitness functions (*ψ*_G_, *ψ*_D_, *ψ*_V_) encoding the expected values of those rates and the scale of random perturbations specified by (*σ*_G_, *σ*_D_, *σ*_V_). This approach, originating from Fudenberg and Harris paper [22], is suitable for modeling heterogeneous tumors, in which subpopulations not only interact [47] but can also vary in their growth rates over time [58]. Adopting the usual probabilistic notation of using capital letters for random variables, we can again start with the subpopulation sizes (*Z*_G_, *Z*_D_, *Z*_V_) evolving based on the *Stochastic Differential Equations* (SDEs) *dZ*_*i*_ = (*ψ*_*i*_*dt*+*σ*_*i*_*dW*_*i*_)*Z*_*i*_, where *i* ∈ {*G, D, V*} and each *W*_*i*_ is a standard one-dimensional Brownian motion, modeling independent perturbations to the fitness of the respective subpopulation. This can be used to derive the SDEs for the corresponding fractions^V^ (*X*_G_, *X*_D_, *X*_V_); see the summary in Box 5 for the reduced (*Q, P*) coordinates and the derivation in SM §2S.2. The terminal set Δ is still the same: the process terminates as soon as *P* (*t*) crosses a stabilization barrier (GLY’s are low, leaving mostly aerobic cells in the tumor) or the failure barrier (GLY’s are high, the patient dies). But the terminal time *T* and the incurred cumulative cost 𝒥 will also be random even if we fix the initial tumor configuration (*q*_0_, *p*_0_) and choose a specific treatment policy *d*(·). Figure 1(b) shows one example of using the deterministic-optimal policy *d*(*t*) = *d*_⋆_ (*Q*(*t*), *P* (*t*)) in this stochastic setting. Gathering statistics from many random simulations that start from the same (*q*_0_, *p*_0_), we can approximate the *Cumulative Distribution Function* (CDF), measuring the probability of keeping 𝒥 below any given threshold *s*:

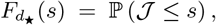

whose graph is shown in blue in Figure 1(c). If one instead opts to solve the PDE (2.9) to maximize the probability of reaching Δ_succ_ while avoiding Δ_fail_, this yields a simple MTD-policy *d* = *d*_max_, whose CDF (shown in brown in Figure 1(c)) is strictly worse than that of *d*_⋆_. This is not surprising since the more selective *d*_⋆_ is quite safe for this particular (*q*_0_, *p*_0_), with Δ_fail_ avoided in all of our 10^5^ simulations. However, its resulting “cost” can be still high in many scenarios. E.g., in 47.4% of the *d*_⋆_-based simulations, 𝒥 exceeded 5; in 72.6% of all cases it exceeded 4.5.

This motivates our optimization approach: deriving a *threshold-aware optimal policy* 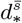 to maximize the probability of stabilization without exceeding a specific cost threshold 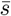. As explained in §2.1 and summarized in Box 4, this is accomplished for a range of threshold values and all initial cancer configurations simultaneously. Figure 1(c) already shows that such policies can provide significant threshold-specific advantages over the deterministic-optimal therapy. Additional simulation results and the actual policies are illustrated in §3.1.

### 2.3 Example 2: a Sensitive-Resistant competition model

We also illustrate our approach by extending a model proposed by Carrère [6], which focuses on the actual size of lung cancer cell populations studied *in vitro*. They consider a heterogeneous tumor that consists of two types of lung cancer cells: the sensitive (*S*) “A549” (sensitive to the drug “Epothilene”) and the resistant (*R*) “A549 Epo40”. A series of *in vitro* experiments were conducted by Manon Carréat the CRO2^VI^ where only the above two types of cells were present. Mutation events were neglected due to their rarity at the considered dosages of Epothilene and due to relatively short treatment durations. The competition model presented below was derived based on phenotypical observations, with fluorescent marking used to trace and differentiate the cells.

Considered separately, both of these types obey a logistic growth model with respective intrinsic growth rates *g*_S_ and *g*_R_. The carrying capacity of the Petri dish (*C*) is assumed to be shared, with the resistant cells assumed to be *m* times bigger than the sensitive; so, the fraction of space used by the time *t* is 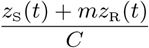. When cultivated together, it was observed that the sensitive cells quickly outgrow the resistant ones despite the fact that their intrinsic growth rates are similar [6]. To model this competitive advantage, they have used an additional competition term *−βz*_S_*z*_R_ to describe the rate of change of *z*_R_(*t*), with the coefficient *β* calibrated based on experimental data. It was further assumed that *R* cells are completely resistant to a specific drug, which reduces the population of *S* cells at the rate of *αz*_S_(*t*)*d*(*t*), with *d*(*t*) reflecting the current rate of drug delivery and the constant coefficient *α* reflecting that drug’s effectiveness. With a normalization *z*_S_(*t*) → *z*_S_(*t*)*/C, z*_R_(*t*) → *z*_R_(*t*)*/C*, the resulting dynamics are summarized by

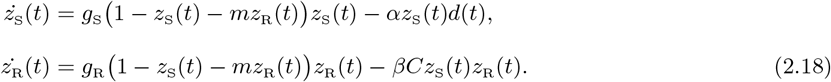

In both the original deterministic case and its stochastic extension, it is more convenient to restate the dynamics in terms of the total size of the tumor *p*(*t*) = *z*_S_(*t*) + *mz*_R_(*t*) and the fraction of tumor size taken by sensitive cells *q*(*t*) = *z*_S_(*t*)*/p*(*t*). This change of coordinates yields an ODE model (2.19) summarized in Box 6; see SM §1S.2 for the derivation. In this case, the goal of our adaptive therapy is eradication: i.e., driving the total tumor size *p*(*t*) below some remission barrier *γ*_r_ (e.g., a physical detection level) while ensuring that throughout the treatment this *p*(*t*) stays below a significantly higher failure barrier *γ*_f_ *<* 1.

Figure 2(a) illustrates the natural dynamics of this model with no drug use. In this case, the competitive pressure reduces the population *R*, which at first decreases the tumor size for many initial conditions. But a rapid growth in *S* eventually increases the overall tumor, leading to an inevitable failure (*p*(*t*) *> γ*_f_). The optimal drug therapy is again sought to minimize a weighted sum of total drugs used and the time of treatment (with the running cost *K* = *d*(*t*) + *δ*) until the eradication. It is obtained in feedback form *d* = *d*_⋆_(*q, p*) after solving the PDE (2.4). Figure 2(b) shows that, for smaller tumor sizes, this *d*_⋆_ prescribes MTD-level treatment only after this initial tumor reduction is over, once *S* gets rid of most *R* cells which are not sensitive to Epothilene. However, for larger initial *p*, this deterministic-optimal policy starts using the drugs much earlier, planning to keep *S* cells in check as soon as they are numerous enough to control *R*.

Stochastic perturbations can be similarly introduced here by assuming that the intrinsic growth rates are actually random and normally distributed at any instant^VII^. In Example 1, we assumed that the fitness function of each subpopulation was affected by its own random perturbations. (The Brownian motion in Box 5 was three-dimensional.) Depending on the nature of perturbations, a similar assumption might be reasonable in the current example as well. But this is not a necessary feature for the threshold-aware optimization approach to be applicable. To demonstrate this, we will instead assume in what follows that a single (1D) Brownian motion perturbs the intrinsic growth rates of both *S* and *R*, with (*g*_S_, *g*_R_) still representing the expected rates and (*σ*_S_, *σ*_R_) denoting their respective volatilities. This yields SDEs for the stochastic evolution of (*Q, P*), which are derived in §2S.3 of SM and summarized in Box 6. As shown in Figure 2(c), if the deterministic-optimal policy *d*_⋆_ is used in this stochastic setting, the initiation time of the MTD-based therapy (and the resulting overall cost 𝒥) can vary significantly. This motivates us again to use the threshold-aware approach based on the PDE (2.11), with the policies illustrated and advantages quantified in §3.2.

#### Box 6

Example 2 (base model adopted from [6])

**The deterministic base model (components for the approach in Box 1):**

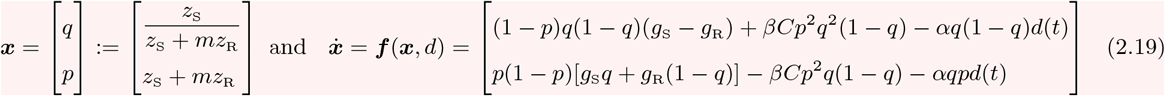

**The stochastic model (components for the approaches in Boxes 2-4):**

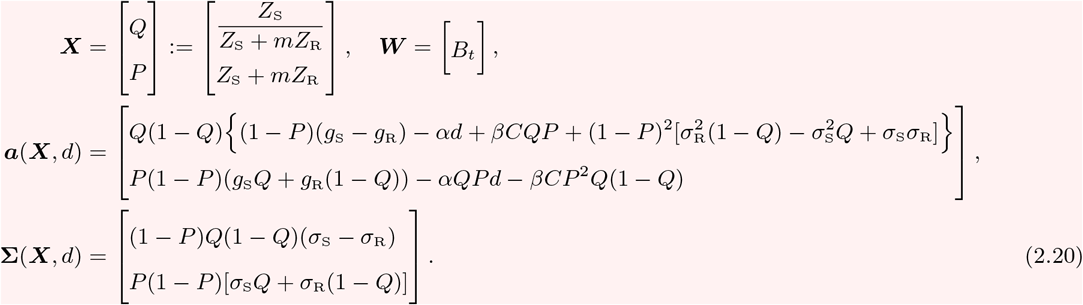

**Definitions and Parameters:**

- *d* : ℝ_+_ → [0, *d*_max_], time-dependent intensity of *S*-targeting therapy;
- Δ_succ_ = {(*q, p*) ∈ [0, 1]^2^ | *p < γ*_r_}, success region where *γ*_r_ is the *remission barrier*;
- Δ_fail_ = {(*q, p*) ∈ [0, 1]^2^ | *p > γ*_f_}, failure region where *γ*_f_ is the *failure barrier*;
- *K*(***X***, *d*) = *d* + *δ*, running cost function where *δ* is the treatment time penalty;
- 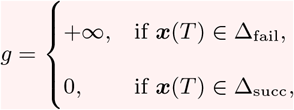 terminal cost;
- (*g*_S_, *g*_R_), growth rate for the sensitive and resistant cells, respectively;
- *B*_*t*_, standard 1D Brownian motion;
- (*σ*_S_, *σ*_R_), volatilities for the sensitive and resistant cells, respectively;
- *m*, size ratio between *S* and *R* cells;
- *C*, Petri dish carrying capacity;
- *α*, drug efficiency;
- *β*, action of sensitive on resistant.

**Parameter values** are specified in Supplementary Materials §5S.2.

**The optimal policy in feedback form:**

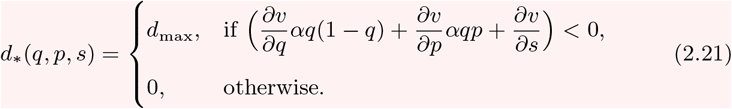

## 3 Results

### 3.1 Policies, trajectories, and CDFs for the EGT-based model

We explore the structure and performance of threshold-aware policies computed for the system described in §2.2. The parameter values *d*_max_ = 3, *b*_*a*_ = 2.5, *b*_*v*_ = 2, *c* = 1, *n* = 4 are the same ones provided in Kaznatcheev et al. [39] and Gluzman et al. [25]. However, we use *γ*_r_ = 1 *− γ*_f_ = 10^*−*2^ and *δ* = 0.05 as opposed to *γ*_r_ = 1 *− γ*_f_ = 10^*−*1.5^ and *δ* = 0.01 in [25]. Additionally, we consider small uniform constant volatilities *σ*_G_ = *σ*_D_ = *σ*_V_ = 0.15, characterizing the scale of random perturbations in fitness function for all 3 cancer subpopulations. The details of our Monte-Carlo simulations used to build all CDFs can be found in SM §4S.3. Additional examples, including those with higher volatilities, in which the threshold-performance advantages are even more significant, can be found in SM §5S.

In Figure 3, we present some representative *s*-slices of threshold-aware optimal policies and their corresponding optimal probability of success for respective threshold values. To be consistent with [25], in all of our figures, the drugs-on region (at the MTD level) is shown in yellow and the drugs-off region is shown in blue. We observe that this drugs-on region is strongly *s*-dependent and completely different from the one in the deterministic-optimal case shown in Figure 1(a). Since the cancer evolution considered here has stochastic dynamics given in (2.5, 2.15), different realizations of random perturbations will result in entirely different sample paths even if the starting configuration and the feedback policy remain the same. Three such representative sample paths are shown in Figure 4, starting from the same initial tumor configuration (*q*_0_, *p*_0_) = (0.26, 0.665) already used in Figure 1 and focusing on a threshold 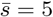. We use the example from Figure 4(a), in which the stabilization is achieved while incurring the total cost of 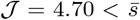, to illustrate the general use of threshold-aware policies. Starting from the initial budget 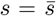, the optimal decision on whether to use drugs right away is based on the first diagram in Figure 3(a). For our initial tumor state, this indicates that *d*_⋆_(*q*_0_, *p*_0_, 5) = 0 (not prescribing drugs initially) would maximize the probability of stabilizing the tumor without exceeding the threshold 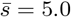. As time passes, we accumulate the cost, thus decreasing the budget, even if the drugs are not used. If we stay in the blue region for the time *θ* = 1*/δ*, the second diagram (the “*s* = 4.0” case) in Figure 3(a) becomes relevant, with subsequent budget decreases shifting us to lower and lower *s* slices. Of course, in reality we constantly reevaluate the decision on *d*_⋆_ (as *s* changes continuously while Figure 3(a) presents just a few representative slices^VIII^) taking into account the changing tumor configuration (*Q*(*t*), *P* (*t*)).

**Figure 3:**
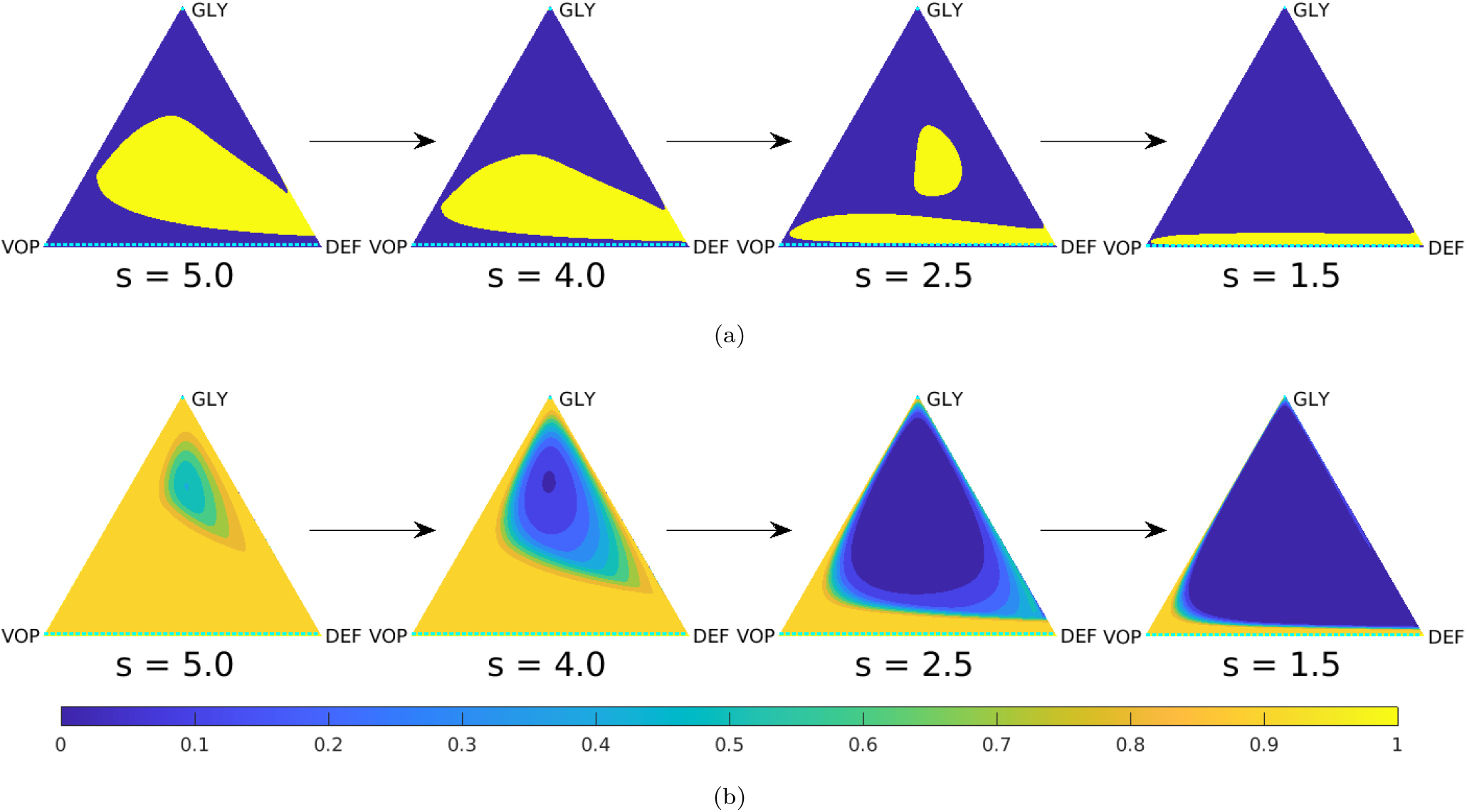
Representative slices of the threshold-aware optimal policy (top row) and the corresponding probability of success (bottom row) for the EGT-based model. Each triangle represents all possible tumor compositions (proportions of GLY/VOP/DEF cells in the population). Top row shows the policy, which prescribes the optimal instantaneous decisions on drug usage given the indicated remaining budget (*s*) and the current tumor state. Bottom row shows the probability of “stabilization within the budget” if the optimal policy is followed from this time point and onward. Each column corresponds to a specific budget level *s*, which is shown below each triangle. The arrows indicate the natural decrease of the remaining budget while implementing the policy.

**Figure 4:**
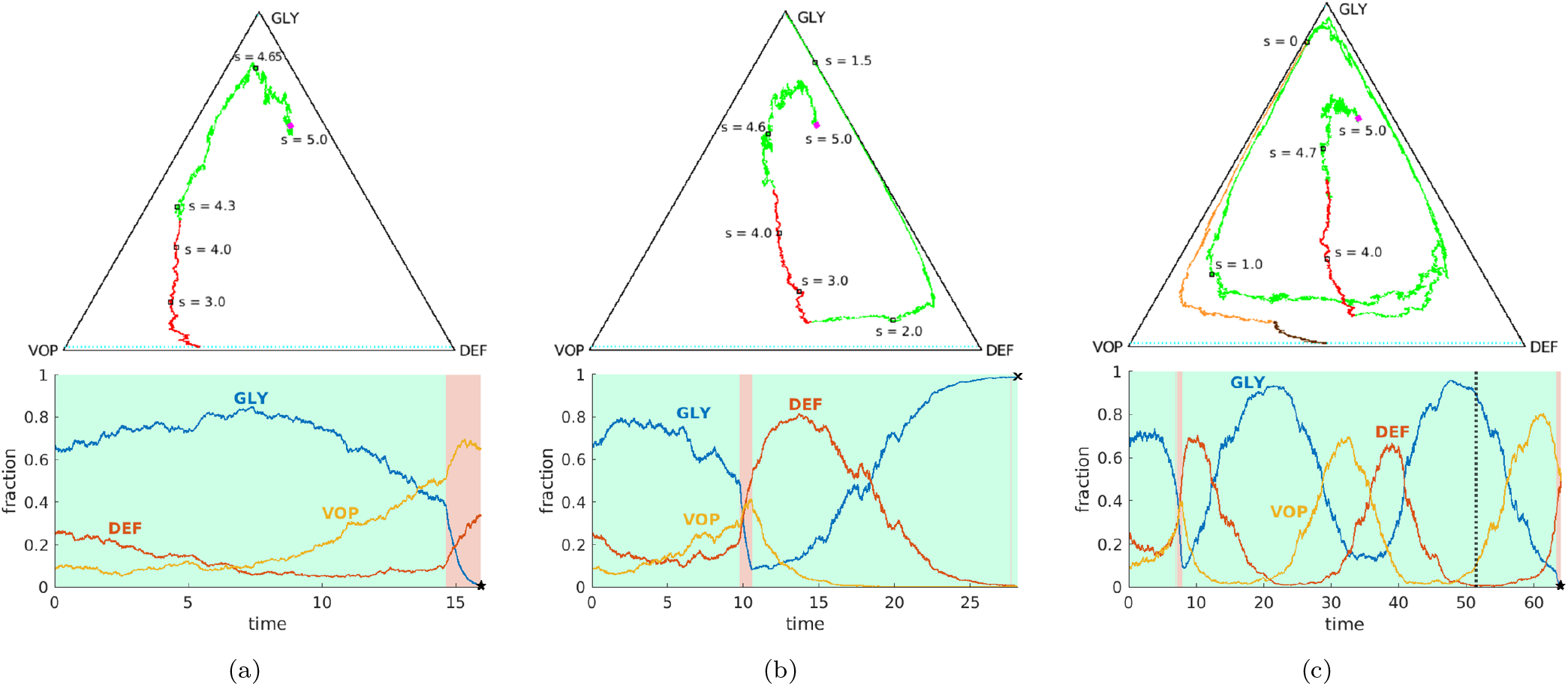
Representative sample paths starting from the same initial state (*q*_0_, *p*_0_) = (0.26, 0.665) (magenta dot) and the same initial budget 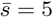. **Top row:** sample paths on a GLY-DEF-VOP triangle. (a) eventual stabilization with a cost of 4.70 (within the budget); (b) eventual death; (c) failure by running out of budget (eventual stabilization with a total cost of 7.80 by switching to the deterministic-optimal policy after *s* = 0). Some representative tumor states along these paths (with indications of how much budget is left) are marked by *black squares*. In (c), the part where 𝒥 *>* 5 is specified in *orange* (no drugs) and *brown* (at MTD level). **Bottom row:** evolution of sub-populations with respect to time based on the sample paths from the top row. Here we use *light green* and *light pink backgrounds* to indicate the time interval(s) of prescribing no drugs and of prescribing drugs at the MTD-rate, respectively. We use *black pentagrams* and *black crosses* to indicate eventual stabilization and death, respectively. In (c), we use a *dashed black* line to indicate the budget depletion time.

In contrast to the success story in Figure 4(a), we note that there are two very different ways of “failing”. First, the process can stop if the proportion of GLY cells becomes too high, as in Figure 4(b). When VOP is relatively low, the deterministic portion of the dynamics can bring us close to the failure barrier, with random perturbations resulting in a noticeable probability of crossing into Δ_fail_. Second, even if we stay away from Δ_fail_, the budget might be exhausted before reaching Δ_succ_, as in Figure 4(c). Threshold-aware policies provide no guidance once *s* = 0, but it is reasonable to continue (using some different treatment policy) since the patient is still alive. In our numerical simulations, we switch in this case to a deterministic-optimal policy *d*_⋆_ illustrated in Figure 1. This decision is somewhat arbitrary; e.g., one could choose instead to switch to an MTD-based policy, which in this example maximizes the probability of reaching Δ_succ_ while avoiding Δ_fail_ without any regard to additional cost incurred thereafter. For this initial tumor configuration and parameter values, continuing with *d*_⋆_ typically yields smaller costs while only slightly increasing the chances of eventual failure (e.g., *≈* 0.45% of crossings into Δ_fail_ using *d*_⋆_ versus *≈* 0.1% using the full MTD once the original budget of 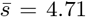 is exhausted). But whatever new policy is chosen for such “unlucky” cases, this choice will only affect the right tail of *J* ‘s distribution; i.e., 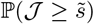 will be affected only for 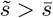.

Returning to the optimal probability of success *v*(*q, p, s*) shown in Figure 3(b), we observe that *v* has particularly large gradient near the level curves of the deterministic-optimal value function *u* shown in Figure 1(a). (The particular level curve of *u* near which *v* changes the most is again *s*-dependent as the budget decreases.) If the remaining budget is relatively low (e.g., *s* = 1.5), one can see from Figure 3(b) that there is no chance to stabilize the tumor within this budget unless the GLY is already low (and a short burst of drug therapy would likely be enough) or VOP is high (and the no-drugs dynamics will bring us to a low GLY concentration later on). Consequently, the optimal policy for *s* = 1.5 is to not use drugs for the majority of tumor states.

The contrast in threshold-specific performance is easy to explain when the deterministic-optimal and threshold-aware policies prescribe different actions from the very beginning. To illustrate this, we consider (*q*_0_, *p*_0_) = (0.27, 0.4), for which *d*_⋆_ = *d*_max_ while 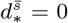 for a range of 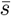 values; see Figures 5(a) and (b) for representative paths and 5(C) for the respective CDFs. Under the deterministic-optimal policy (whose CDF is shown in blue), only 50% of simulations yield the cost not exceeding 4.71. A threshold-aware policy (implemented for 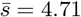, with CDF shown in pink) maximizes this 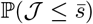 and succeeds in 63.7% of all cases. The potential for improvement is even more significant with lower threshold values. For instance, we see that ℙ( 𝒥 (*d*_⋆_) ≤ 4.35) *<* 10%, while our threshold-aware policy (implemented for 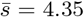, with CDF shown in orange) ensures that 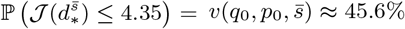. This improvement can also be translated to simple medical terms: starting from this initial tumor configuration, the deterministic-optimal policy will likely keep using the drugs at the maximum rate *d*_max_ all the way to stabilization; see Figure 5(a). In contrast, our threshold-aware policies tend not to prescribe drugs until GLY is relatively low and VOP is relatively high; see Figure 5(b). As a result, the patient would suffer less toxicity from drugs in most scenarios.

**Figure 5:**
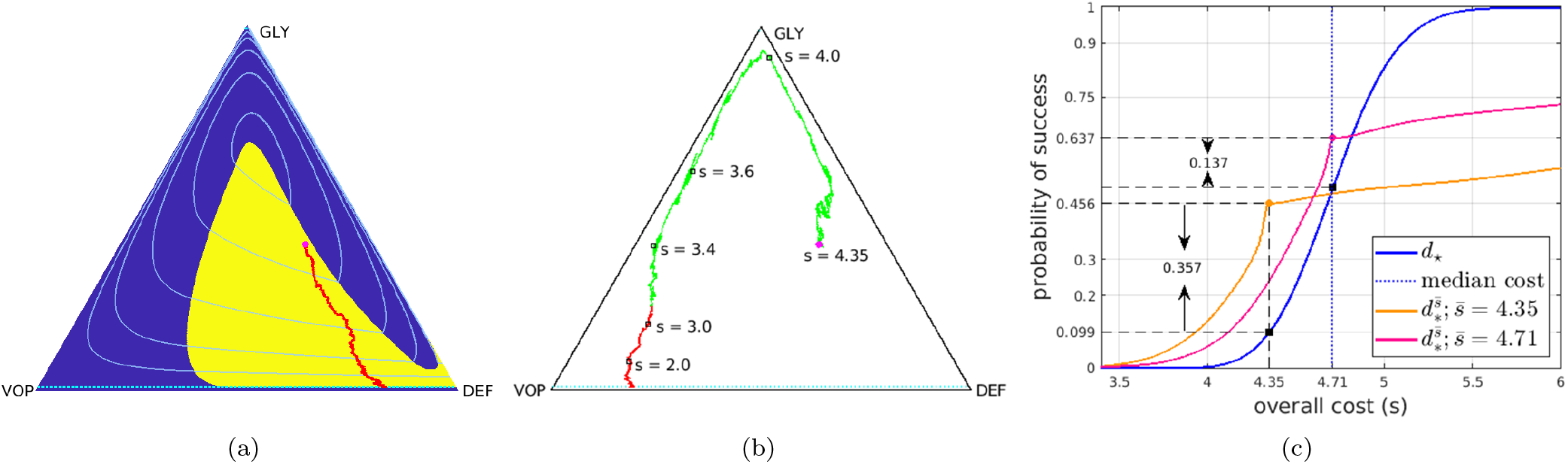
Comparison between threshold-aware policies and the deterministic-optimal policy. Starting from an initial state (*q*_0_, *p*_0_) = (0.27, 0.4) (magenta dot): (a) a sample path with cost 4.75 under the deterministic-optimal policy; (b) a sample path starting at 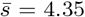 with a realized total cost of 4.02 under the (orange) threshold-aware policy; (c) CDFs of the cumulative cost 𝒥 approximated using 10^5^ random simulations. In (c), the *solid blue* curve is the CDF generated with the deterministic-optimal policy. Its median (*dashed blue* line) is 4.71 while its mean conditioning on success is 4.72. The *solid orange* curve is the CDF generated with the threshold-aware policy with 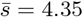 and the *solid pink* curve is the CDF generated with the threshold-aware policy with 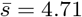.

It is worth noting that each threshold-aware policy maximizes the probability of success for a single/specific threshold value only. E.g., for all the pink/orange CDFs we have provided, the probability of success is only maximized at those pink/orange dots. Moreover, we clearly see from Figure 5(c) that the probability of 𝒥 not exceeding *any* 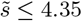 is lower on the pink CDF than on the orange CDF (computed for 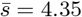). Intuitively, this is not too surprising. In the early stages of treatment, a (pink) policy computed to maximize the chances of not exceeding 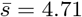 is more aggressive in using the drugs and thus spends the “budget” quicker than the (orange) policy, which starts from a lower initial budget 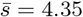. This is also consistent with the budget-dependent sizes of drugs-on regions in Figure 3(a).

### 3.2 Policies, trajectories, and CDFs for the SR-model

We now turn to the SR model system described in §2.3. Our numerical experiments use *d*_max_ = 3, *γ*_r_ = 1 *− γ*_f_ = 10^*−*2^, *δ* = 0.05, and volatilities *σ*_R_ = *σ*_S_ = 0.15. For other parameter values, see §5S.2 of SM.

We show the representative *s*-slices of threshold aware policies and the corresponding success probabilities in Figure 6. Similarly to the EGT-model, we observe that the drug-on regions (shown in yellow) are strongly budget-dependent and quite different from the ones specified by *d*_⋆_ in Figure 2(b). We note that the drugs-on region generally shrinks in size (toward the *Q* = 1 line, where only *S* cells are present) as the budget *s* decreases. For even tighter budgets, this yellow region becomes disconnected, prescribing the drugs for large *P* values (to substantially decrease the tumor size) and in a thin layer near Δ_succ_ (where a short burst of drugs is likely sufficient).

**Figure 6:**
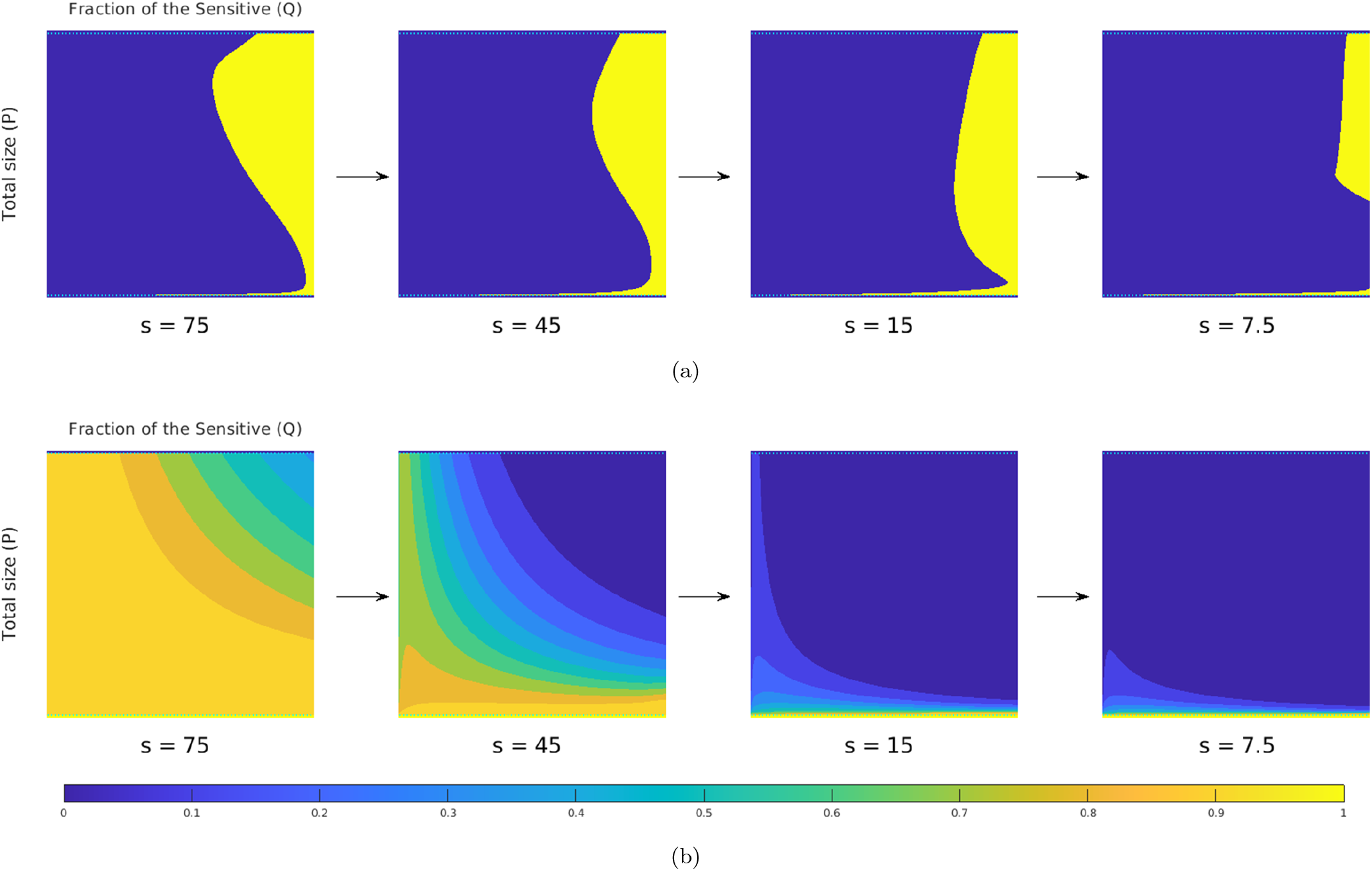
Representative slices of the threshold-aware optimal policy (top row) and the corresponding probability of success (bottom row) for the Carrère example. Each square represents all possible tumor states (sizes and compositions). The horizontal axis is the *fraction of the Sensitive (Q)* and the vertical axis is the *total population (P)*. Top row shows the policy, which prescribes the optimal instantaneous decisions on drug usage given the indicated remaining budget (*s*) and the current tumor state. Bottom row shows the probability of “eradication within the budget” if the optimal policy is followed from this time point and onward. Each column corresponds to a specific budget level *s*, which is shown below each square. The arrows indicate the natural decrease of the remaining budget while implementing the policy.

In Figure 7, we consider a particular initial tumor configuration (*q*_0_, *p*_0_) = (0.45, 0.9) and compare the performance of three different policies. Despite the fact that all three use no drugs at the very beginning, the deterministic-optimal policy typically starts prescribing drugs much earlier. See the comparison of sample trajectories under *d*_⋆_ and 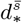 in Figures 7(a) and 7(b). As a result, our threshold-aware policy (implemented for 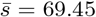, with CDF shown in pink) improves 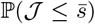 to 67.4% from 50% produced by *d*_⋆_. This advantage is even more significant with lower thresholds. E.g., ℙ (𝒥 (*d*_⋆_) ≤ 60) is only 19.6%, while our threshold-aware policy (CDF shown in orange) more than doubles this probability of under-threshold remission to 39.8%.

**Figure 7:**
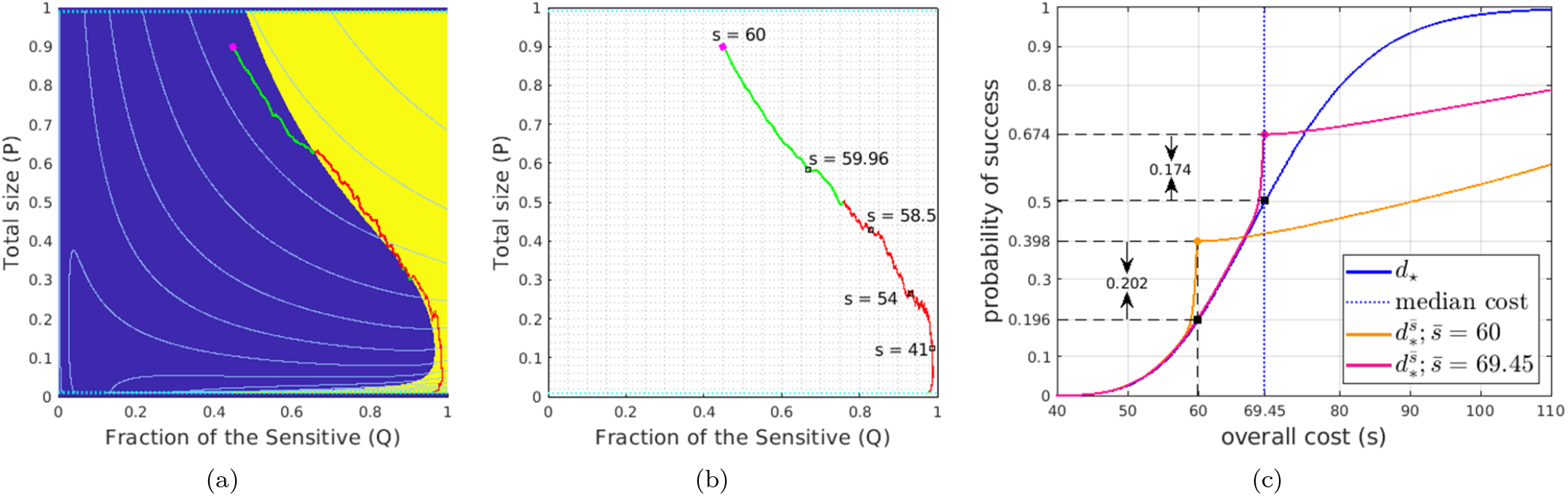
Comparison between threshold-aware policies and the deterministic-optimal policy. Starting from an initial state (*q*_0_, *p*_0_) = (0.45, 0.9) (magenta dot): (a) a sample path with cost 57.3 under the deterministic-optimal policy; (b) a sample path starting at 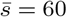 with a total cost of 53.7 under the (orange) threshold-aware policy; (c) CDFs of the cumulative cost 𝒥 with 10^5^ samples. In (c), the *solid blue* curve is the CDF generated with the deterministic-optimal policy. Its median (*dashed blue* line) is 69.45 while its mean conditioning on success is 70.5. The *solid orange* curve is the CDF generated with the threshold-aware policy with 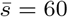 and the *solid pink* curve is the CDF generated with the threshold-aware policy with 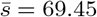.

## 4 Discussion

That cancers evolve during therapy is now an accepted fact, and is slowly being incorporated into therapeutic decision making. In some cases, this can be implemented simply by changing from one targeted therapy to another, but in most, where tumors are a heterogeneous mixture of interacting phenotypes, this is not feasible. In these cases, ecological thinking is rising to the fore in the form of adaptive therapy. Until recently, clinical trials, and theoretical investigations, of adaptive therapy have relied on *a priori* assumptions of the underlying interactions, and their effects on tumor composition over time. Several studies, both *in vitro*[38] and *in vivo*[17, 47], *however, have begun to provide methods for more rigorous quantification of these interactions. As these tools mature, the next challenges will be to understand these interactions in patients and to exploit them in improving personalized treatment*.

*The presented approach is a step in this direction, aiming to limit the probability of high-cost outcomes in the presence of stochastic perturbations. It is applicable to a broad class of stochastic cancer models and therapy goals (e*.*g*., *tumor eradication or stabilization). While it is standard to tune the treatment plan to maximize the probability of reaching its goal, we go farther and maximize the probability of goal attainment without exceeding a prescribed threshold on cumulative cost (interpreted as a combination of the total drugs used, cumulative disease burden, and the time to remission/stabilization). We show that these optimal treatment policies become threshold-aware*, with the drugs-on/drugs-off regions changing as the treatment progresses and the initial “cost budget” (for meeting the chosen threshold) gradually decreases. The comparison of CDFs generated for the deterministic-optimal policy and threshold-aware policies demonstrates clear advantages of the latter, often resulting in a significant reduction of drugs used to treat the patient.

More generally, dynamic programming provides an excellent framework for finding optimal treatment policies by solving Hamilton-Jacobi-Bellman (HJB) equations. The fact that these policies are recovered in feedback form makes this approach particularly suitable for optimization of adaptive therapies. But even though the use of general optimal control in cancer treatment is by now common [55], the same is not true for the more robust HJB-based methods, which so far have been used in only a handful of cancer-related applications [1, 7, 19, 25, 36, 42, 53]. This is partly due to the HJBs’ well-known *curse of dimensionality*: the rapid increase in computational costs when the system state becomes higher-dimensional. This is a relevant limitation since our threshold-aware approach introduces the “budget” as an additional component of the state. Similarly to the presented examples, our current implementation would be easy to adopt to any cancer model based on a two-dimensional (*q, p*) state space, with the budget *s* adding the third dimension. For cancer models with a larger number of subpopulations, the general approach would remain the same, but the approximate HJB-solver would likely need to rely on sparse grids [48], tensor decompositions [15], or deep neural networks [64].

The presented examples did not model any mutations, but we note that this is not really a limitation of the method itself. E.g., drug-usage-dependent mutations would be easy to incorporate into our EGT-based example by switching to a Replicator-Mutator ODE/SDE eco-evolutionary model [28, 32]. We did not pursue such examples here primarily to make for an easier comparison with prior work [25] and to limit the number of model parameters.

Our SDE models of global (or environmental) stochastic perturbations to subpopulation fitnesses and intrinsic growth rates are based on perspectives well-established in biological applications [5, 22]. While our focus on environmental stochasticity is motivated by “averaging-out” the variability within each subpopulation, it is worth noting that this assumption is not always justifiable. Whenever the subpopulation size becomes sufficiently small, the demographic stochasticity becomes crucially important. (This is also the regime in which the validity of ODE/SDE models is far less obvious.) Even though we do not deal with this important limitation here, we note that our threshold-aware approach can be used with a variety of perturbation types^IX^, including jump-diffusion processes, which could be used to build future models that account for demographic stochasticity in these special small-subpopulation regimes. Dynamic programming is also used in discrete population models focused on demographic stochasticity [49]. It will be also interesting to investigate the usability of our approach in that discrete setting.

Sensitivity with respect to threshold variation can be tested by comparing CDFs of 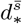 for different 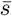 values. While it is also possible to perform a similar comparison under perturbation of model parameters, we believe that another approach is more promising: any bounded uncertainty in parameter values can be treated as a “game against nature,” leading to a Hamilton-Jacobi-Isaacs PDE, whose solution will yield policies optimizing the threshold-performance in the “worst parameter variation” scenarios [8].

Another important extension will be to move to “partial observability” since the state of the tumor is only occasionally assessed directly through biopsies and some proxy measurements have to be used at all other times [19]. Finally, it will be also interesting to study the multiobjective control problem of optimizing threshold-aware policies for two different threshold values simultaneously.

In summary, we have presented a theoretical and computational advance for the toolbox of evolutionary therapy, a new subfield of medicine focused on using knowledge of evolutionary responses to inform therapeutic scheduling. While there are a number of cancer trials using this type of evolutionary-informed thinking, most are based on heuristic designs and are not formulated to consider the underlying stochasticities. Developing a theoretical foundation for future clinical studies requires EGT models directly grounded in objectively measurable biology [38]. Therapy optimization based on such models requires efficient computational methods, particularly in the presence of stochastic perturbations. We hope that the general approach presented here will be useful for a broad range of increasingly accurate stochastic cancer models.

## Supporting information

Mathematical details & additional examples.

## Funding and Acknowledgements

The authors are grateful to Roberto Ferretti and Lars Grüne for their advice on some aspects of numerical methods used in this project. The authors would also like to thank the anonymous reviewers for their very insightful and helpful comments. MW and AV are grateful to the National Science Foundation (DMS-1645643, DMS-1738010, and DMS-2111522) for supporting this project. JGS would like to thank the National Institutes of Health (5R37CA244613, 5U54CA274513, U01CA280829) and the American Cancer Society (RSG-20-096-01) for their generous support.

## SM Supplementary Materials

Mathematical details and additional computational experiments.

## Footnotes

I E.g., in eradication therapy models, Δ_succ_ might correspond to tumors below the detection level, while Δ_fail_ might specify a much larger size that effectively kills a patient; see §2.3. On the other hand, for models that only track the relative abundance of cancer subpopulations, Δ_succ_ might be defined in terms of the desired low abundance of specific subpopulations affected by ***d***(*t*), with the idea that the tumor size stabilizes or an entirely different therapy strategy is adopted after ***x***(*t*) enters Δ_succ_; see §2.2.

II In classical stochastic optimal control, parabolic HJB equations are usually encountered when dealing with finite horizon problems, where the terminal time *T* is specified in advance. See [5, 19, 52] for typical examples in cancer-related literature.In contrast, the parabolicity in PDE (2.11) arises because of the monotone decrease in the remaining budget *s*(*t*).

III The details of our numerical method are included in Supplementary Materials (SM) §4S.1. In the interest of computational reproducibility, we provide the source code for approximating value functions and computing threshold-aware policies for all the examples from §3 at https://github.com/eikonal-equation/Stochastic-Cancer

IV In [39], this goal is justified by noting that, with GLY gone, the DEF cells will then quickly overcome VOP, leading to “an aerobic tumor with no - or significantly diminished - ability to recruit blood vessels,” which stabilizes (or at least significantly slows down the growth of) the tumor.

V We note that Related Stochastic Replicator Equations arise naturally in ecology, where they have been studied in depth to address a possible coexistence of species in randomly perturbed environments [31, 56].

VI Center for Research in Oncobiology and Oncopharmacology, Aix-Marseille Université

VII This approach was also used in modeling persistence strategies among bacteria in [5].

VIII Movies with additional information for Figures 3 and 4 are available at https://eikonal-equation.github.io/Stochastic-Cancer/examples.html

IX A similar threshold-aware method has been developed in [8] for controlling “piecewise-deterministic” processes, where perturbations happen at discrete points in time and amount to abrupt switches in system dynamics. More recently, it has been also used to control the hybrid dynamics of a sailboat navigating in stochastically changing wind conditions and trying to reach the destination prior to a specified deadline 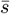 [59].

## Notes

https://eikonal-equation.github.io/Stochastic-Cancer/

### Competing Interest Statement

The authors have declared no competing interest.

### Summary of Updates

A significant revision shifting the focus of presentation to the method, which is now presented first for an abstract/general system and only then illustrated by two specific stochastic cancer models. Expanded bibliography and additional context. Many new figures.

