## Supplementary material for "Threshold-awareness in adaptive cancer therapy": Mathematical details & additional examples.

#### Abstract

This document provides additional details for the manuscript "Threshold-awareness in adaptive cancer therapy." Note the numbering of figures continues from the main text. We continue to use reference numbers of equations and figures previously introduced in the main text. Bibliographic references used in Supplementary Materials (SM) are listed and numbered separately.

We provide the full source code of our numerical schemes presented in §4S and movies for both small (section 3.1 in the main text) and large volatilities (§5S.3) at
`https://eikonal-equation.github.io/Stochastic-Cancer/`.

### 1S Derivations of controlled unperturbed/deterministic systems

#### 1S.1 The EGT competition model

We start by producing the main derivations of the controlled unperturbed system of equations (labeled (2.14) in the main text) adopted from Gluzman et al. [16]. The original model from Kaznatcheev et al. [20] describes a competition of 3 types of cancer cells: Glycolytic cells (GLY) are anaerobic and produce lactic acid. The other two types are aerobic and benefit from the oxygen from vascularization. The VEGF (over)-producing cells (VOP) improve the vasculature development, while the remaining aerobic cells are defectors (DEF). As mentioned in our paper, the competition of cells in the tumor is thus modeled as a "public goods" / "club goods" game: VEGF is a "club good" since it benefits only VOP and DEF cells, while the acid generated by GLY is a "public good" as it benefits all cancer cells.

The original model in [20] is based on a game of $(n+1)$ locally interacting cancer cells. The fitness of each of them depends on the proportion of all three types among these game participants. Define $A_n(k) = \dfrac{b_a k}{n+1}$ to be the benefit to each cell due to acidity if $k$ of $(n+1)$ participants are producing acid. Suppose $(m+1)$ of these $(n+1)$ cells are aerobic. Then $V_m(k) = \dfrac{b_v k}{m+1}$ is defined to be the benefit (to each aerobic cell) due to increased vascularization if $k$ of these $(m+1)$ cells are (over) producing VEGF at a personal cost $c$.

Focusing on one specific participant, suppose $n_{\mathrm{G}}, n_{\mathrm{D}}, n_{\mathrm{V}}$ are the numbers of GLY, DEF, and VOP cells among all others, with $n_{\mathrm{G}}+n_{\mathrm{D}}+n_{\mathrm{V}}=n$. Thus, if that "focus participant" is a GLY cell, the benefit due to acidity is $A_n(n_{\mathrm{G}}+1)$ (it is actually its payoff since GLY is the only anaerobic type). Similarly, if the focus participant is an aerobic type, the benefit due to acidity becomes $A_n(n_{\mathrm{G}})$. When the focus participant is a VOP cell, the benefit it receives due to vascularization is $V_{n-n_{\mathrm{G}}}(n_{\mathrm{V}}+1)$ while paying a constant cost $c$ to produce VEGF. Note that the group size of aerobic cells among $n$ total nearby cells is $n-n_{\mathrm{G}}$. On the other hand, if the focus participant is a DEF cell, it receives a benefit of $V_{n-n_{\mathrm{G}}}(n_{\mathrm{V}})$ due to vascularization while paying no cost. We note that the benefit received by a focus participant depends heavily on that participant's type and on the types of others. To determine the fitness functions $\psi$, Kaznatcheev et al. further assumed that all participants are drawn uniformly at random from a large well-mixed population. They then computed the *expected benefit* for each type of focus participants, by averaging over all possible compositions of their local interaction group [20].

**To simplify the notation for the EGT model, we will use subscripts 1, 2, and 3 to indicate GLY, DEF and VOP cells respectively.** Summarizing the above in this new notation, we list the fitness function for each of the three types of cancer cells:

$$\begin{cases} \psi_1 = \langle A_n(n_1+1)\rangle_{n_1\sim B_n(x_1)}, \\ \psi_2 = \langle A_n(n_1)\rangle_{n_1\sim B_n(x_1)} + \langle V_{n-n_1}(n_3)\rangle_{(n_1,n_3)\sim M_n(x_1,x_3)}, \\ \psi_3 = \langle A_n(n_1)\rangle_{n_1\sim B_n(x_1)} + \langle V_{n-n_1}(n_3+1)\rangle_{(n_1,n_3)\sim M_n(x_1,x_3)} - c, \end{cases} \tag{1S.1.1}$$

where $B_n(x)$ is the binomial distribution with $n$ samples and $x$ is the probability of success, $M_n(x,y)$ is the multinomial distribution with $n$ samples, three possible outcomes with respective probabilities $(x,\ 1-x-y,\ y)$; and $\langle f(\zeta)\rangle_{\zeta\sim\Omega}$ is the expected value of $f(\zeta)$ over a random variable $\zeta$ with distribution $\Omega$. The expectations in (1S.1.1) can be also computed explicitly; we refer to Section 1S of SM in [16] and Appendix A in [20] for details.

Let $z_1, z_2, z_3$ denote the absolute sizes of subpopulations of GLY, DEF, VOP cells. We assume each sub-population grows with rate equal to its respective fitness, and the GLY-targeting therapy (control) kills GLY cells only with time-dependent rate $d:\mathbb{R}_+\to[0,d_{\max}]$. Then the controlled dynamics of sub-populations is

$$\begin{cases} \dot z_1(t) = \psi_1(t)z_1(t) - d(t)z_1(t), \\ \dot z_2(t) = \psi_2(t)z_2(t), \\ \dot z_3(t) = \psi_3(t)z_3(t). \end{cases}$$

By definition, the proportion of GLY cells in the entire tumor is $x_1 = z_1/(z_1+z_2+z_3)$. Differentiating both sides with respect to $t$, we obtain the dynamics for $x_1$:

$$\begin{aligned}
\dot{x}_1 &= \frac{\dot{z}_1}{z_1+z_2+z_3} - \frac{(\dot{z}_1+\dot{z}_2+\dot{z}_3)x_1}{z_1+z_2+z_3} \\
&= \frac{z_1}{z_1+z_2+z_3}(\psi_1 - d(t)) - \left[\frac{z_1}{z_1+z_2+z_3}(\psi_1 - d(t)) + \frac{z_2}{z_1+z_2+z_3}\psi_2 + \frac{z_3}{z_1+z_2+z_3}\psi_3\right] x_1 \\
&= (\psi_1 - dd(t))x_1 - [x_1(\psi_1 - d(t)) + x_2\psi_2 + x_3\psi_3]\, x_1 \\
&= (\psi_1 - dd(t))x_1 + dd(t)\; x_1^2 - \langle\psi\rangle x_1 \\
&= (\psi_1 - \langle\psi\rangle)x_1 - x_1(1-x_1)d(t),
\end{aligned}$$

where $\langle\psi\rangle = x_1\psi_1 + x_2\psi_2 + x_3\psi_3$ is the average fitness.

The ODEs for $\dot{x}_2$ and $\dot{x}_3$ follow a similar argument, and hence the controlled unperturbed system (a replicator equation) is:

$$\begin{cases}
\dot{x}_1 = (\psi_1 - \langle\psi\rangle)x_1 - x_1(1-x_1)d(t), \\
\dot{x}_2 = (\psi_2 - \langle\psi\rangle)x_2 + x_2x_1d(t), \\
\dot{x}_3 = (\psi_3 - \langle\psi\rangle)x_3 + x_3x_1d(t).
\end{cases}$$

Since $x_1 + x_2 + x_3 = 1$, one can transform the above 3-dimensional system into 2-dimensional. Let the proportion of glycolytic cells in the tumor to be $p(t) = x_1(t)$ and the proportion of VOP among aerobic cells to be $q(t) = x_3(t)/(x_2(t) + x_3(t))$.

Hence

$$\begin{aligned}
\dot{p} &= p(\psi_1 - \langle\psi\rangle) - p(1-p)d(t) \\
&= p(\psi_1 - x_1\psi_1 - x_2\psi_2 - x_3\psi_3) - p(1-p)d(t) \\
&= p(\psi_1 - p\psi_1 - x_2\psi_2 - x_3\psi_3) - p(1-p)d(t) \\
&= p(1-p)(\psi_1 - \langle\psi\rangle_{2,3} - d(t)),
\end{aligned}$$

where $\langle\psi\rangle_{2,3} = \psi_2 + q(\psi_3 - \psi_2)$.

Similarly,

$$\dot{q} = \frac{\dot{x}_3x_2 - \dot{x}_2x_3}{(x_2+x_3)^2} = \frac{x_3x_2}{(x_2+x_3)^2}(\psi_3 - \psi_2) = q(1-q)(\psi_3 - \psi_2).$$

By definitions of $\psi_1, \psi_2, \psi_3$ from above, one can find

$$\begin{cases} \psi_3 - \psi_2 = \dfrac{b_v}{n+1}\left[\sum\limits_{k=0}^{n} p^k\right] - c, \\ \psi_1 - \langle\psi\rangle_{2,3} = \dfrac{b_a}{n+1} - q(b_v - c). \end{cases}$$

See Section 1S of SM of [16] for detailed derivations of the above expressions. Substituting them into equations for $\dot{p}$ and $\dot{q}$, one obtains the controlled reduced unperturbed system (labeled (2.14) in Box 5 in the main text):

$$\begin{cases} \dot{q} = q(1-q)\left(\dfrac{b_v}{n+1}\left[\sum\limits_{k=0}^{n} p^k\right] - c\right), \\ \dot{p} = p(1-p)\left(\dfrac{b_a}{n+1} - (b_v - c)q - d(t)\right). \end{cases} \tag{1S.1.2}$$

#### 1S.2 The SR competition model

We now provide derivations of the controlled unperturbed system of equations (labeled (2.19) in the main text) based on a model proposed by Carrère [8]. As explained in the main text, their original model describes competition between two types of lung cancer cells *in vitro*: the sensitive ($S$) "A549" (sensitive to the drug "Epothilene") and the resistant ($R$) "A549 Epo40". The model was derived based on phenotypical observations and neglected mutation events.

We denote the absolute size of their respective population as $z_{\text{S}}$ and $z_{\text{R}}$. The concentration of the drug (control) has a time-dependent rate $d : \mathbb{R}_+ \to [0, d_{\text{max}}]$. With a normalization $z_{\text{S}}(t) \to z_{\text{S}}(t)/C$, $z_{\text{R}}(t) \to z_{\text{R}}(t)/C$, we have shown in the main text that the controlled dynamics follow

$$\begin{aligned} \dot{z_{\text{S}}}(t) &= g_{\text{S}}\big(1 - z_{\text{S}}(t) - m z_{\text{R}}(t)\big) z_{\text{S}}(t) - \alpha z_{\text{S}}(t) d(t), \\ \dot{z_{\text{R}}}(t) &= g_{\text{R}}\big(1 - z_{\text{S}}(t) - m z_{\text{R}}(t)\big) z_{\text{R}}(t) - \beta C z_{\text{S}}(t) z_{\text{R}}(t). \end{aligned} \tag{1S.2.1}$$

Focusing on the total size of the tumor $p(t) = z_{\text{S}}(t) + m z_{\text{R}}(t)$ and the fraction of tumor size taken by sensitive cells $q(t) = z_{\text{S}}(t)/p(t)$, we now derive the ODEs in this new coordinates $(q(t), p(t))$.

$$\begin{aligned} \dot{p} &= \dot{z_{\text{S}}} + m\dot{z_{\text{R}}} \\ &= g_{\text{S}}\big(1 - z_{\text{S}} - m z_{\text{R}}\big) z_{\text{S}} - \alpha z_{\text{S}} d(t) + g_{\text{R}}\big(1 - z_{\text{S}} - m z_{\text{R}}\big) z_{\text{R}} - \beta C z_{\text{S}} z_{\text{R}} \\ &= g_{\text{S}} q p(1-p) - \alpha q p\, d(t) + g_{\text{R}}(1-q)p(1-p) - \beta C p^2 q(1-q) \\ &= p(1-p)[g_{\text{S}} q + g_{\text{R}}(1-q)] - \beta C p^2 q(1-q) - \alpha q p\, d(t) \end{aligned}$$

$$
\begin{aligned}
\dot{q} &= \frac{\dot{z}_{\mathrm{S}}}{p} - q \cdot \frac{\dot{p}}{p} \\
&= \frac{g_{\mathrm{S}}\big(1-p\big)qp - \alpha qp\, d(t)}{p} - q \cdot \frac{p(1-p)[g_{\mathrm{S}}q + g_{\mathrm{R}}(1-q)] - \beta C p^2 q(1-q) - \alpha qp\, d(t)}{p} \\
&= g_{\mathrm{S}}q(1-p) - \alpha q\, d(t) - q(1-p)[g_{\mathrm{S}}q + g_{\mathrm{R}}(1-q)] + \alpha q^2 d(t) + \beta C p q^2(1-q) \\
&= g_{\mathrm{S}}q(1-p)(1-q) - g_{\mathrm{R}}q(1-q)(1-p) + \beta C p q^2(1-q) - \alpha q(1-q)d(t) \\
&= (1-p)q(1-q)(g_{\mathrm{S}} - g_{\mathrm{R}}) + \beta C p q^2(1-q) - \alpha q(1-q)d(t)
\end{aligned}
$$

We thus obtains the controlled unperturbed system (labeled (2.19) in Box 6 in the main text) in $(q, p)$ coordinates:

$$
\begin{cases}
\dot{q} = (1-p)q(1-q)(g_{\mathrm{S}} - g_{\mathrm{R}}) + \beta C p q^2(1-q) - \alpha q(1-q)d(t), \\
\dot{p} = p(1-p)[g_{\mathrm{S}}q + g_{\mathrm{R}}(1-q)] - \beta C p^2 q(1-q) - \alpha qp\, d(t).
\end{cases}
\tag{1S.2.2}
$$

### 2S Problem setup and derivations for stochastic models

#### 2S.1 Problem setup under a general stochastic optimal control framework

Let $(\Omega, \mathcal{F}, \{\mathcal{F}_t\}_{t\geq 0}, \mathbb{P})$ denote a complete filtered probability space. The canonical reference probability space for a standard $m$-dimensional Brownian motion is defined by the following 5-tuple [10, 14]

$$
\nu := \big(\Omega, \mathcal{F}, \mathbb{P}, \{\mathcal{F}_t\}_{t\geq 0}, \mathbf{W}\big),
$$

where

- $\Omega := C([0,\infty), \mathbb{R}^n)$ is the space of all continuous functions from $[0,\infty)$ to $\mathbb{R}^n$ equipped with the metric of uniform convergence on compact sets;
- $\mathcal{F} := \mathcal{B}(\Omega)$ is the Borel $\sigma$-algebra on $\Omega$;
- $\mathbb{P}$ is the Wiener measure;
- $\{\mathcal{F}_t\}_{t\geq 0}$ is the completion of the natural filtration of an $m$-dimensional Brownian motion defined on $(\Omega, \mathcal{F}, \mathbb{P})$;
- $\boldsymbol{W} = (W_1, W_2, \ldots W_m) : [0,\infty) \times \Omega \to \mathbb{R}^m$ in $\mathbb{R}^m$ is an $\mathcal{F}_t$-Brownian motion.

Let $\boldsymbol{d}(\cdot) : [0,\infty) \times \Omega \to \mathcal{D}$ ($\mathcal{D}$ compact) be a $\mathcal{F}_t$-progressively measurable control process. By "$\mathcal{F}_t$-progressively measurable", we mean for each $t \in [0,\infty)$, the map $(t,\omega) \to \boldsymbol{d}(t,\omega)$ from $[0,t] \times \Omega$ into $[0, d_{\max}]$ is $\mathcal{B}([0,t]) \otimes \mathcal{F}_t$-

measurable [10, 14]. For the sake of notational simplicity, we will suppress the $\omega$-dependence in the rest of this document.

We focus on stochastic dynamics of the following form:

$$\mathrm{d}\boldsymbol{X} = \boldsymbol{a}(\boldsymbol{X}, \boldsymbol{d})\ \mathrm{d}t + \boldsymbol{\Sigma}(\boldsymbol{X}, \boldsymbol{d})\ \mathrm{d}\boldsymbol{W}, \tag{2S.1.1}$$

where $\boldsymbol{X} \in \mathbb{R}^n$ encodes the state variables, $\boldsymbol{a}(\boldsymbol{X}, \boldsymbol{d}) \in \mathbb{R}^n$ denotes the drift coefficient function, and $\boldsymbol{\Sigma}(\boldsymbol{X}, \boldsymbol{d}) \in \mathbb{R}^{n\times m}$ encodes the diffusion coefficient function. If $\boldsymbol{a}(Y, \boldsymbol{d})$ and $\boldsymbol{\Sigma}(Y, \boldsymbol{d})$ are *Lipschitz* in $Y$ and *uniformly continuous* in $\boldsymbol{d}$, then Eq. (2S.1.1) has a strong solution for a given I.C. [6, 9]

Consider an exit-time problem where $\Delta \subset [0, 1]^n$ is the terminal set (for simplicity, we have normalized $\boldsymbol{X} \in [0, 1]^n$). Furthermore, $\Delta = \Delta_{\text{succ}} \cup \Delta_{\text{fail}}$ is the union of the success region $\Delta_{\text{succ}}$ and the failure region $\Delta_{\text{fail}}$. We define the terminal time as

$$T = T(\boldsymbol{\xi}, \boldsymbol{d}(\cdot)) := \inf \Big\{ t \in \mathbb{R}_+ \ \mid\ \boldsymbol{X}(t) \in \Delta, \quad \boldsymbol{X}(0) = \boldsymbol{\xi} \Big\},$$

where $\boldsymbol{\xi}$ is any starting configuration.

We further define the total cost of using a policy $\boldsymbol{d}(\cdot)$ as

$$\mathcal{J}(\boldsymbol{\xi}, \boldsymbol{d}(\cdot)) \;=\; \int_0^T K(\boldsymbol{X}(\tau), \boldsymbol{d}(\tau))\, \mathrm{d}\tau \;+\; g\Big(\boldsymbol{X}(T)\Big),$$

for some *strictly positive* running cost function $K$ and a some non-negative terminal cost function $g$.

#### 2S.2 Derivation of controlled perturbed system for Example 1

In this section, we provide detailed derivations of the controlled stochastic cancer dynamics for our first example (labeled (2.15) in the main text).

##### 2S.2.1 Derivation of controlled perturbed system for the fractions $(X_1, X_2, X_3)$

We start by providing detailed derivations of the controlled stochastic cancer dynamics for the fractions $(X_1, X_2, X_3)$. As mentioned in the main text, we choose the stochastic replicator equation originated from Fudenberg and Harris [15] to describe the driven dynamics.

In the ordinary replicator equation shown in §1S.1, we assume the actual sub-population size of each type of cancer cells $z_i$ grows with rate $\psi_i$. For the stochastic replicator equation, standard Brownian motion is added to the fitness function $\psi_i$ as the source of perturbation. As a result, the driven differential equation for $Z_i$ (we adopt the convention to use capital letters to denote random variables in this document as well) is modeled by geometric Brownian motion. The growth rate for $Z_1$ will be further affected by our control - the rate of drug administration $d(\cdot)$.

The controlled stochastic dynamics for sub-populations $Z_1, Z_2, Z_3$ is then:

$$\begin{cases} \mathrm{d}Z_1 = [(\psi_1 - d(t))\,\mathrm{d}t + \sigma_1\,\mathrm{d}W_1]\,Z_1, \\ \mathrm{d}Z_2 = [\psi_2\,\mathrm{d}t + \sigma_2\,\mathrm{d}W_2]\,Z_2, \\ \mathrm{d}Z_3 = [\psi_3\,\mathrm{d}t + \sigma_3\,\mathrm{d}W_3]\,Z_3, \end{cases}$$

where $(W_1, W_2, W_3)$ is a standard 3-dimensional Brownian motion for $(Z_1, Z_3, Z_3)$, and $(\sigma_1, \sigma_2, \sigma_3) \geq 0$ are the constant volatilities. In such a way, the expected fitness of each type is still $\psi_i$ since standard Brownian motion has zero mean.

We first derive the Stochastic Differential Equation (SDE) for $X_1 = Z_1/(Z_1 + Z_2 + Z_3)$. By Itô's lemma, for multivariate function $f(X, Y, Z)$,

$$\begin{aligned} \mathrm{d}f = f_x\,\mathrm{d}X + f_y\,\mathrm{d}Y + f_z\,\mathrm{d}Z + \frac{1}{2}f_{xx}\,(\mathrm{d}X)^2 + \frac{1}{2}f_{yy}\,(\mathrm{d}Y)^2 + \frac{1}{2}f_{zz}\,(\mathrm{d}Z)^2 \\ + f_{xy}\,(\mathrm{d}X)(\mathrm{d}Y) + f_{xz}\,(\mathrm{d}X)(\mathrm{d}Z) + f_{yz}\,(\mathrm{d}Y)(\mathrm{d}Z). \end{aligned}$$

For $X_1$, we have $f(x, y, z) = \dfrac{x}{x+y+z}$, then

$$\begin{aligned} f_x &= \frac{y+z}{(x+y+z)^2}, \; f_y = -\frac{x}{(x+y+z)^2}, \; f_z = -\frac{x}{(x+y+z)^2} \\ f_{xx} &= -\frac{2(y+z)}{(x+y+z)^3}, \; f_{yy} = \frac{2x}{(x+y+z)^3}, \; f_{zz} = \frac{2x}{(x+y+z)^3} \\ f_{xy} &= \frac{x-y-z}{(x+y+z)^3}, \; f_{xz} = \frac{x-y-z}{(x+y+z)^3}, \; f_{yz} = \frac{2x}{(x+y+z)^3}. \end{aligned}$$

By convention, for independent Brownian motions,

$$(\mathrm{d}t)^2 = 0, \; \mathrm{d}t\,\mathrm{d}W_i = 0\;\forall i, \; \mathrm{d}W_i\,\mathrm{d}W_j = 0\;\forall i \neq j, \; (\mathrm{d}W_i)^2 = \mathrm{d}t\;\forall i.$$

Thus,

$$
\begin{aligned}
\mathrm{d}X_1 &= \mathrm{d}\left(\frac{Z_1}{Z_1+Z_2+Z_3}\right) \\
&= f_x\ \mathrm{d}Z_1 + f_y\ \mathrm{d}Z_2 + f_z\ \mathrm{d}Z_3 + \frac{1}{2} f_{xx}\ (\mathrm{d}Z_1)^2 + \frac{1}{2} f_{yy}\ (\mathrm{d}Z_2)^2 + \frac{1}{2} f_{zz}\ (\mathrm{d}Z_3)^2 \\
&\quad + f_{xy}\ (\mathrm{d}Z_1)(\mathrm{d}Z_2) + f_{xz}\ (\mathrm{d}Z_1)(\mathrm{d}Z_3) + f_{yz}\ (\mathrm{d}Z_2)(\mathrm{d}Z_3) \\
&= \frac{Z_2+Z_3}{(Z_1+Z_2+Z_3)^2}\left[(\psi_1 - d(t))\ \mathrm{d}t + \sigma_1\ \mathrm{d}W_1\right] Z_1 \\
&\quad - \frac{Z_1}{(Z_1+Z_2+Z_3)^2}\left[\psi_2\ \mathrm{d}t + \sigma_2\ \mathrm{d}W_2\right] Z_2 - \frac{Z_1}{(Z_1+Z_2+Z_3)^2}\left[\psi_3\ \mathrm{d}t + \sigma_3\ \mathrm{d}W_3\right] Z_3 \\
&\quad - \frac{1}{2}\frac{2(Z_2+Z_3)}{(Z_1+Z_2+Z_3)^3} Z_1^2\sigma_1^2\ \mathrm{d}t + \frac{1}{2}\frac{2Z_1}{(Z_1+Z_2+Z_3)^3} Z_2^2\sigma_2^2\ \mathrm{d}t + \frac{1}{2}\frac{2Z_1}{(Z_1+Z_2+Z_3)^3} Z_3^2\sigma_3^2\ \mathrm{d}t \\
&= \left[(X_2+X_3)\psi_1 - (X_2+X_3)d(t) - X_2\psi_2 - X_3\psi_3\right] X_1\ \mathrm{d}t + \left[(X_2+X_3)\sigma_1\ \mathrm{d}W_1 - X_2\sigma_2\ \mathrm{d}W_2 - X_3\sigma_3\ \mathrm{d}W_3\right] X_1 \\
&\quad - \left[X_1(X_2+X_3)\sigma_1^2 - X_2^2\sigma_2^2 - X_3^2\sigma_3^2\right] X_1\ \mathrm{d}t \\
&= \left[X_1(\psi_1 - \langle\psi\rangle) - X_1(1-X_1)d(t)\right]\ \mathrm{d}t + \left(\sigma_1\ \mathrm{d}W_1 - \sum_{j=1}^{3} X_j\sigma_j\ \mathrm{d}W_j\right) X_1 - \left(\sigma_1^2 X_1 - \sum_{j=1}^{3} X_j^2\sigma_j^2\right) X_1\ \mathrm{d}t.
\end{aligned}
$$

For $X_2 = Z_2/(Z_1+Z_2+Z_3)$, we have $f(x,y,z) = \dfrac{y}{x+y+z}$. Then

$$
\begin{aligned}
f_x &= -\frac{y}{(x+y+z)^2},\ f_y = \frac{x+z}{(x+y+z)^2},\ f_z = -\frac{y}{(x+y+z)^2} \\
f_{xx} &= \frac{2y}{(x+y+z)^3},\ f_{yy} = -\frac{2(x+z)}{(x+y+z)^3},\ f_{zz} = \frac{2y}{(x+y+z)^3} \\
f_{xy} &= \frac{y-x-z}{(x+y+z)^3},\ f_{xz} = \frac{2y}{(x+y+z)^3},\ f_{yz} = \frac{y-x-z}{(x+y+z)^3}.
\end{aligned}
$$

Thus,

$$
\begin{aligned}
\mathrm{d}X_2 &= \mathrm{d}\left(\frac{Z_2}{Z_1+Z_2+Z_3}\right) \\
&= -\frac{Z_2}{(Z_1+Z_2+Z_3)^2}\left[(\psi_1 - d(t))\ \mathrm{d}t + \sigma_1\ \mathrm{d}W_1\right] Z_1 \\
&\quad + \frac{Z_1+Z_3}{(Z_1+Z_2+Z_3)^2}\left[\psi_2\ \mathrm{d}t + \sigma_2\ \mathrm{d}W_2\right] Z_2 - \frac{Z_2}{(Z_1+Z_2+Z_3)^2}\left[\psi_3\ \mathrm{d}t + \sigma_3\ \mathrm{d}W_3\right] Z_3 \\
&\quad + \frac{1}{2}\frac{2Z_2}{(Z_1+Z_2+Z_3)^3} Z_1^2\sigma_1^2\ \mathrm{d}t - \frac{1}{2}\frac{2(Z_1+Z_3)}{(Z_1+Z_2+Z_3)^3} Z_2^2\sigma_2^2\ \mathrm{d}t + \frac{1}{2}\frac{2Z_2}{(Z_1+Z_2+Z_3)^3} Z_3^2\sigma_3^2\ \mathrm{d}t \\
&= \left[-X_1\psi_1 + X_1 d(t) + (X_1+X_3)\psi_2 - X_3\psi_3\right] X_2\ \mathrm{d}t + \left[-X_1\sigma_1\ \mathrm{d}W_1 + (X_1+X_3)\sigma_2\ \mathrm{d}W_2 - X_3\sigma_3\ \mathrm{d}W_3\right] X_2 \\
&\quad - \left[-X_1^2\sigma_1^2 + (X_1+X_3)X_2\sigma_2^2 - X_3^2\sigma_3^2\right] X_2\ \mathrm{d}t \\
&= \left[X_2(\psi_2 - \langle\psi\rangle) + X_2 X_1 d(t)\right]\ \mathrm{d}t + \left(\sigma_2\ \mathrm{d}W_2 - \sum_{j=1}^{3} X_j\sigma_j\ \mathrm{d}W_j\right) X_2 - \left(\sigma_2^2 X_2 - \sum_{j=1}^{3} X_j^2\sigma_j^2\right) X_2\ \mathrm{d}t.
\end{aligned}
$$

Similarly,

$$\mathrm{d}X_3 = [X_3(\psi_3 - \langle\psi\rangle) + X_3X_1d(t)]\ \mathrm{d}t + \left(\sigma_3\ \mathrm{d}W_3 - \sum_{j=1}^{3} X_j\sigma_j\ \mathrm{d}W_j\right) X_3 - \left(\sigma_3^2 X_3 - \sum_{j=1}^{3} X_j^2\sigma_j^2\right) X_3\ \mathrm{d}t.$$

Hence the controlled stochastic replicator dynamics is:

$$\begin{cases} \mathrm{d}X_1 = & \Big[X_1(\psi_1 - \langle\psi\rangle) - X_1(1-X_1)d(t)\Big]\ \mathrm{d}t + \left(\sigma_1\ \mathrm{d}W_1 - \sum_{j=1}^{3} X_j\sigma_j\ \mathrm{d}W_j\right) X_1 \\ & - \left(\sigma_1^2 X_1 - \sum_{j=1}^{3} X_j^2\sigma_j^2\right) X_1\ \mathrm{d}t, \\ \mathrm{d}X_2 = & \Big[X_2(\psi_2 - \langle\psi\rangle) + X_2X_1d(t)\Big]\ \mathrm{d}t + \left(\sigma_2\ \mathrm{d}W_2 - \sum_{j=1}^{3} X_j\sigma_j\ \mathrm{d}W_j\right) X_2 \\ & - \left(\sigma_2^2 X_2 - \sum_{j=1}^{3} X_j^2\sigma_j^2\right) X_2\ \mathrm{d}t, \\ \mathrm{d}X_3 = & \Big[X_3(\psi_3 - \langle\psi\rangle) + X_3X_1d(t)\Big]\ \mathrm{d}t + \left(\sigma_3\ \mathrm{d}W_3 - \sum_{j=1}^{3} X_j\sigma_j\ \mathrm{d}W_j\right) X_3 \\ & - \left(\sigma_3^2 X_3 - \sum_{j=1}^{3} X_j^2\sigma_j^2\right) X_3\ \mathrm{d}t. \end{cases} \tag{2S.2.1}$$

##### 2S.2.2 Derivation of SDEs of Example 1 in reduced coordinates

We now derive the SDEs in the reduced coordinates $(Q,\ P)$. Since $X_1+X_2+X_3 = 1$ in the stochastic case as well, we can again take advantage of this and transform the 3-dimensional system into a 2-dimensional one. Let $P(t) = X_1(t)$ and $Q(t) = \dfrac{X_3(t)}{X_2(t) + X_3(t)}$ as in §1S.1.

Then with $X_2 = (1-P)(1-Q)$ and $X_3 = (1-P)Q$, we have

$$\begin{aligned}\mathrm{d}P = &\Big[P(\psi_1 - \langle\psi\rangle) - P(1-P)d(t)\Big]\ \mathrm{d}t + \Big(\sigma_1\ \mathrm{d}W_1 - \Big\{P\sigma_1\ \mathrm{d}W_1 + (1-P)(1-Q)\sigma_2\ \mathrm{d}W_2 + (1-P)Q\sigma_3\ \mathrm{d}W_3\Big\}\Big)P \\ &- \Big[\sigma_1^2 P - (\sigma_1^2 P^2 + \sigma_2^2(1-P)^2(1-Q)^2 + \sigma_3^2(1-P)^2Q^2)\Big]P\ \mathrm{d}t.\end{aligned}$$

Let

$$\begin{cases} \phi(Q,P;\mathrm{d}W_1,\mathrm{d}W_2,\mathrm{d}W_3) & = P\sigma_1\ \mathrm{d}W_1 + (1-P)(1-Q)\sigma_2\ \mathrm{d}W_2 + (1-P)Q\sigma_3\ \mathrm{d}W_3, \\ \lambda(Q,P) & = \sigma_1^2P^2 + \sigma_2^2(1-P)^2(1-Q)^2 + \sigma_3^2(1-P)^2Q^2. \end{cases}$$

Then $\phi^2 = \lambda(Q,P)\ \mathrm{d}t$. Therefore, the SDE for $P(t)$ can be rewritten as

$$\mathrm{d}P = [P(\psi_1 - \langle\psi\rangle) - P(1-P)d(t)]\ \mathrm{d}t + \big[\sigma_1\ \mathrm{d}W_1 - \phi\big]P - \big[\sigma_1^2 P - \lambda\big]\ P\ \mathrm{d}t. \tag{2S.2.2}$$

By Itô's lemma, for multivariate function $f(X, Y)$,

$$\mathrm{d}f = f_x\ \mathrm{d}X + f_y\ \mathrm{d}Y + \frac{1}{2} f_{xx}\ (\mathrm{d}X)^2 + \frac{1}{2} f_{yy}\ (\mathrm{d}Y)^2 + f_{xy}\ (\mathrm{d}X)(\mathrm{d}Y)$$

For $Q = X_3/(X_2 + X_3)$, we have $f(x, y) = \dfrac{x}{x+y}$, then

$$f_x = \frac{y}{(x+y)^2},\ f_y = -\frac{x}{(x+y)^2},\ f_{xx} = -\frac{2y}{(x+y)^3},\ f_{yy} = \frac{2x}{(x+y)^3},\ f_{xy} = \frac{x-y}{(x+y)^3}.$$

Hence,

$$\begin{aligned}
\mathrm{d}Q &= \mathrm{d}\left(\frac{X_3}{X_2 + X_3}\right) \\
&= \frac{X_2}{(X_2+X_3)^2}\ \mathrm{d}X_3 - \frac{X_3}{(X_2+X_3)^2}\ \mathrm{d}X_2 - \frac{1}{2}\frac{2X_2}{(X_2+X_3)^3}\ (\mathrm{d}X_3)^2 + \frac{1}{2}\frac{2X_2}{(X_2+X_3)^3}\ (\mathrm{d}X_2)^2 \\
&\quad + \frac{X_3 - X_2}{(X_2+X_3)^3}\ (\mathrm{d}X_2)(\mathrm{d}X_3) \\
&= \frac{(1-P)(1-Q)}{(1-P)^2}\ \mathrm{d}X_3 - \frac{(1-P)Q}{(1-P)^2}\ \mathrm{d}X_2 - \frac{(1-P)(1-Q)}{(1-P)^3}\ (\mathrm{d}X_3)^2 + \frac{(1-P)Q}{(1-P)^3}\ (\mathrm{d}X_2)^2 \\
&\quad + \frac{(1-P)Q - (1-P)(1-Q)}{(1-P)^3}\ (\mathrm{d}X_2)(\mathrm{d}X_3) \\
&= \frac{1-Q}{1-P}\ \mathrm{d}X_3 - \frac{Q}{1-P}\ \mathrm{d}X_2 - \frac{1-Q}{(1-P)^2}\ (\mathrm{d}X_3)^2 + \frac{Q}{(1-P)^2}\ (\mathrm{d}X_2)^2 + \frac{2Q-1}{(1-P)^2}\ (\mathrm{d}X_2)(\mathrm{d}X_3).
\end{aligned}$$

Now substituting (2S.2.1) and (2S.2.2) into the above equation, we have

$$\begin{aligned}
\mathrm{d}Q = \frac{1-Q}{1-P}\Big[&(1-P)Q(\psi_3 - \langle\psi\rangle)\ \mathrm{d}t + d(t)PQ(1-P)\ \mathrm{d}t \\
&+ (1-P)Q(\sigma_3\ \mathrm{d}W_3 - \phi) - (1-P)Q(\sigma_3^2(1-P)Q - \lambda)\ \mathrm{d}t\Big] \\
- \frac{Q}{1-P}\Big[&(1-P)(1-Q)(\psi_2 - \langle\psi\rangle)\ \mathrm{d}t + d(t)P(1-P)(1-Q)\ \mathrm{d}t \\
&+ (1-P)(1-Q)(\sigma_2\ \mathrm{d}W_2 - \phi) - (1-P)(1-Q)(\sigma_2^2(1-P)(1-Q) - \lambda)\mathrm{d}t\Big] \\
- \frac{1-Q}{(1-P)^2}\ &\left[(1-P)^2Q^2(\sigma_3^2\ \mathrm{d}t + \phi^2 - 2(1-P)Q\sigma_3^2\ \mathrm{d}t)\right] \\
+ \frac{Q}{(1-P)^2}\ &\left[(1-P)^2(1-Q)^2(\sigma_2^2\ \mathrm{d}t + \phi^2 - 2(1-P)(1-Q)\sigma_2^2\ \mathrm{d}t)\right] \\
+ \frac{2Q-1}{(1-P)^2}\ &\left[(1-P)^2Q(1-Q)(\phi^2 - (1-P)(1-Q)\sigma_2^2\ \mathrm{d}t - (1-P)Q\sigma_3^3\ \mathrm{d}t)\right] \\
= Q(1-Q)&(\psi_3 - \psi_2)\ \mathrm{d}t + \cancel{d(t)PQ(1-Q)\ \mathrm{d}t} - \cancel{d(t)PQ(1-Q)\ \mathrm{d}t} \\
+ Q(1-Q)&(\sigma_3\ \mathrm{d}W_3 - \sigma_2\ \mathrm{d}W_2) - Q^2(1-Q)\sigma_3^2\ \mathrm{d}t + Q(1-Q)^2\sigma_2^2\ \mathrm{d}t \\
= Q(1-Q)&(\psi_3 - \psi_2)\ \mathrm{d}t + Q(1-Q)(\sigma_3\ \mathrm{d}W_3 - \sigma_2\ \mathrm{d}W_2) + Q(1-Q)\ \left[(1-Q)\sigma_2^2 - Q\sigma_3^2\right]\ \mathrm{d}t.
\end{aligned}$$

Recall from §1S.1 that

$$\begin{cases} P(\psi_1 - \langle\psi\rangle) = P(1-P)\left(\dfrac{b_a}{n+1} - (b_v - c)Q\right), \\ \psi_3 - \psi_2 = \dfrac{b_v}{n+1}\left[\sum\limits_{k=0}^{n} P^k\right] - c. \end{cases}$$

Thus, the controlled reduced stochastic system provided in Box 5 in the main text is

$$\begin{cases} \mathrm{d}Q = Q(1-Q)\left(\dfrac{b_v}{n+1}\left[\sum\limits_{k=0}^{n} P^k\right] - c\right)\,\mathrm{d}t + Q(1-Q)\Big(\sigma_3\,\mathrm{d}W_3 - \sigma_2\,\mathrm{d}W_2\Big) \\ \qquad + \Big[(1-Q)\sigma_2^2 - Q\sigma_3^2\Big]Q(1-Q)\,\mathrm{d}t, \\ \mathrm{d}P = P(1-P)\left(\dfrac{b_a}{n+1} - (b_v - c)Q - d(t)\right)\,\mathrm{d}t \\ \qquad + \sigma_1 P(1-P)\,\mathrm{d}W_1 + \sigma_2 P(1-P)(1-Q)\,\mathrm{d}W_2 + \sigma_3 P(1-P)Q\,\mathrm{d}W_3 \\ \qquad - \Big[\sigma_1^2 P - \sigma_2^2(1-P)(1-Q)^2 - \sigma_3^2(1-P)Q^2\Big]P(1-P)\,\mathrm{d}t. \end{cases} \tag{2S.2.3}$$

Since $Q,\ P \in [0,1]$ and $d \in [0, d_{\max}]$, the drift coefficient function of the above system is Lipschitz with respect to $Q$, $P$, and $d$, and the diffusion coefficient function of the above system is Lipschitz with respect to $Q$ and $P$. Consequently, with any fixed initial data $(q_0, p_0)$, the above system has a unique strong solution on our reference probability space $\nu$ [6, 9].

#### 2S.3 Derivation of controlled perturbed system for Example 2

Next, we provide detailed derivations for the controlled stochastic cancer dynamics for our second example (labeled (2.20) in the main text).

As mentioned in the main text, for this example we assume a single (1D) Brownian motion $B_t$ affecting the intrinsic growth rates of both $S$ and $R$ simultaneously, with expectation $(g_{\mathrm{S}}, g_{\mathrm{R}})$ and volatilities $(\sigma_{\mathrm{S}}, \sigma_{\mathrm{R}})$. This results in the following SDEs for $(Z_{\mathrm{S}}, Z_{\mathrm{R}})$:

$$\begin{aligned} \mathrm{d}Z_{\mathrm{S}} &= (g_{\mathrm{S}}\mathrm{d}t + \sigma_{\mathrm{S}}\,\mathrm{d}B_t)(1 - Z_{\mathrm{S}} - mZ_{\mathrm{R}})Z_{\mathrm{S}} - \alpha d(t) Z_{\mathrm{S}}\mathrm{d}t \\ &= [g_{\mathrm{S}}(1 - Z_{\mathrm{S}} - mZ_{\mathrm{R}}) - \alpha d(t)]Z_{\mathrm{S}}\mathrm{d}t + \sigma_{\mathrm{S}} Z_{\mathrm{S}}(1 - Z_{\mathrm{S}} - mZ_{\mathrm{R}})\mathrm{d}B_t. \\ \mathrm{d}Z_{\mathrm{R}} &= (g_{\mathrm{R}}\mathrm{d}t + \sigma_{\mathrm{R}}\,\mathrm{d}B_t)(1 - Z_{\mathrm{S}} - mZ_{\mathrm{R}})Z_{\mathrm{R}} - \beta C Z_{\mathrm{S}} Z_{\mathrm{R}}\mathrm{d}t \\ &= [g_{\mathrm{R}}(1 - Z_{\mathrm{S}} - mZ_{\mathrm{R}}) - \beta C Z_{\mathrm{S}}]Z_{\mathrm{R}}\mathrm{d}t + \sigma_{\mathrm{R}} Z_{\mathrm{R}}(1 - Z_{\mathrm{S}} - mZ_{\mathrm{R}})\mathrm{d}B_t. \end{aligned}$$

Let $P(t) = Z_{\rm S}(t) + mZ_{\rm R}(t)$ and $Q(t) = \dfrac{Z_{\rm S}(t)}{P(t)}$ as in §1S.2. It follows that $Z_{\rm S} = QP$, $mZ_{\rm R} = (1-Q)P$, and hence

$$\begin{aligned}
\mathrm{d}P &= \mathrm{d}Z_{\rm S} + m\mathrm{d}Z_{\rm R} \\
&= (g_{\rm S}\mathrm{d}t + \sigma_{\rm S}\,\mathrm{d}B_t)(1 - Z_{\rm S} - mZ_{\rm R})Z_{\rm S} - \alpha d(t) Z_{\rm S}\mathrm{d}t \\
&= [g_{\rm S}(1 - Z_{\rm S} - mZ_{\rm R}) - \alpha d(t)]Z_{\rm S}\mathrm{d}t + \sigma_{\rm S} Z_{\rm S}(1 - Z_{\rm S} - mZ_{\rm R})\mathrm{d}B_t \\
&\quad + [g_{\rm R}(1 - Z_{\rm S} - mZ_{\rm R}) - \beta C Z_{\rm S}]Z_{\rm R}\mathrm{d}t + \sigma_{\rm R} Z_{\rm R}(1 - Z_{\rm S} - mZ_{\rm R})\mathrm{d}B_t \\
&= \big[g_{\rm S} QP(1-P) - \alpha QPd(t) + g_{\rm R}(1-Q)P(1-P) - \beta CP^2 Q(1-Q)\big]\mathrm{d}t \\
&\quad + \big[\sigma_{\rm S} QP(1-P) + \sigma_{\rm R}(1-Q)P(1-P)\big]\mathrm{d}B_t \\
&= \Big[P(1-P)(g_{\rm S}Q + g_{\rm R}(1-Q)) - \alpha QPd(t) - \beta CP^2Q(1-Q)\Big]\mathrm{d}t \\
&\quad + P(1-P)\Big[\sigma_{\rm S}Q + \sigma_{\rm R}(1-Q)\Big]\mathrm{d}B_t
\end{aligned}$$

By Itô's lemma, for multivariate function $f(X, Y)$,

$$\mathrm{d}f = f_x\ \mathrm{d}X + f_y\ \mathrm{d}Y + \frac{1}{2} f_{xx}\ (\mathrm{d}X)^2 + \frac{1}{2} f_{yy}\ (\mathrm{d}Y)^2 + f_{xy}\ (\mathrm{d}X)(\mathrm{d}Y)$$

For $Q = Z_{\rm S}/P$, we have $f(x,y) = \dfrac{x}{y}$, then

$$f_x = \frac{1}{y},\ f_y = -\frac{x}{y^2},\ f_{xx} = 0,\ f_{yy} = \frac{2x}{y^3},\ f_{xy} = -\frac{1}{y^2}.$$

Hence,

$$\begin{aligned}
\mathrm{d}Q &= \mathrm{d}\left(\frac{Z_{\rm S}}{P}\right) \\
&= \frac{1}{P}\mathrm{d}Z_{\rm S} - \frac{Z_{\rm S}}{P^2}\mathrm{d}P + \frac{Z_{\rm S}}{P^3}(\mathrm{d}P)^2 - \frac{1}{P^2}(\mathrm{d}Z_{\rm S})(\mathrm{d}P) \\
&= \frac{1}{P}\Big\{\big[g_{\rm S}QP(1-P) - \alpha QPd(t)\big]\mathrm{d}t + \sigma_{\rm S}QP(1-P)\mathrm{d}B_t\Big\} \\
&\quad - \frac{Q}{P}\Big\{\Big[P(1-P)(g_{\rm S}Q + g_{\rm R}(1-Q)) - \alpha QPd - \beta CP^2Q(1-Q)\Big]\mathrm{d}t + P(1-P)\Big[\sigma_{\rm S}Q + \sigma_{\rm R}(1-Q)\Big]\mathrm{d}B_t\Big\} \\
&\quad + \frac{Q}{P^2}\Big\{P^2(1-P)^2\big[\sigma_{\rm S}Q + \sigma_{\rm R}(1-Q)\big]^2\Big\}\mathrm{d}t \\
&\quad - \frac{1}{P^2}\Big\{\sigma_{\rm S}QP(1-P)\cdot P(1-P)\big[\sigma_{\rm S}Q + \sigma_{\rm R}(1-Q)\big]\Big\}\mathrm{d}t \\
&= \Big[(1-Q)Q(1-P)(g_{\rm S} - g_{\rm R}) + \beta CPQ^2(1-Q) - \alpha Q(1-Q)d(t)\Big]\mathrm{d}t \\
&\quad + Q(1-Q)(1-P)(\sigma_{\rm S} - \sigma_{\rm R})\mathrm{d}B_t \\
&\quad + Q(1-Q)(1-P)^2\Big[\sigma_{\rm R}^2(1-Q) - \sigma_{\rm S}^2 Q + \sigma_{\rm S}\sigma_{\rm R}\Big]\mathrm{d}t
\end{aligned}$$

Thus, the controlled reduced stochastic system provided in Box 6 in the main text is

$$\begin{cases} \mathrm{d}Q = Q(1-Q)\Big\{(1-P)(g_{\mathrm{S}}-g_{\mathrm{R}}) - \alpha d(t) + \beta CQP + (1-P)^2[\sigma_{\mathrm{R}}^2(1-Q) - \sigma_{\mathrm{S}}^2 Q + \sigma_{\mathrm{S}}\sigma_{\mathrm{R}}]\Big\}\mathrm{d}t \\ \qquad + Q(1-Q)(1-P)(\sigma_{\mathrm{S}}-\sigma_{\mathrm{R}})\mathrm{d}B_t, \\ \mathrm{d}P = \Big[P(1-P)(g_{\mathrm{S}}Q + g_{\mathrm{R}}(1-Q)) - \alpha QPd(t) - \beta CP^2Q(1-Q)\Big]\mathrm{d}t + P(1-P)\Big[\sigma_{\mathrm{S}}Q + \sigma_{\mathrm{R}}(1-Q)\Big]\mathrm{d}B_t. \end{cases} \tag{2S.3.1}$$

By the same argument as in §2S.2.2, with any fixed initial data $(q_0, p_0)$, the above system has a unique strong solution on our reference probability space $\nu$ [6, 9].

### 3S Derivation of Hamilton-Jacobi-Bellman equations

We derive the threshold-awareness HJB equation (labeled (2.11) in Box 4 of the main text) via tools of dynamic programming in §3S.2. We also summarize the objective and the first-order HJB equation in the deterministic case since we extensively compare our threshold-aware optimal policy to the deterministic-optimal policy in the main text. (The derivation of PDEs in Box 3 is omitted; it follows the standard methods in [14].)

#### 3S.1 The first-order HJB equation in the deterministic case

In this section, we derive the first-order HJB PDE (2.4) provided in Box 1 in the main text. As mentioned there, we consider a general ODE describing the deterministic cancer dynamics:

$$\dot{\boldsymbol{x}} = \boldsymbol{f}(\boldsymbol{x}, \boldsymbol{d}), \qquad \boldsymbol{x}(0) = \boldsymbol{\xi}. \tag{3S.1.1}$$

We follow the same definitions (but now in the deterministic setting) of the terminal time, terminal cost function $g$, running cost function $K$ and the total cost function $\mathcal{J}$ as presented in §2S.1. The objective (value function) minimizing the (deterministic) total cost $\mathcal{J}$ over all available (deterministic) policies is now defined as

$$u(\boldsymbol{\xi}) = \inf_{\boldsymbol{d}(\cdot)} \mathcal{J}\big(\boldsymbol{\xi}, \boldsymbol{d}(\cdot)\big), \tag{3S.1.2}$$

and a policy $\boldsymbol{d}_\star(\cdot)$ is optimal if $u(\boldsymbol{\xi}) = \mathcal{J}\big(\boldsymbol{\xi}, \boldsymbol{d}_\star(\cdot)\big)$. To simplify the derivation, we assume that such $\boldsymbol{d}_\star$ exists. (Otherwise, a similar argument can be built using $\epsilon$-suboptimal policies.) For a sufficiently small $\theta > 0$, by Bellman's

Optimality Principle we have

$$
\begin{aligned}
u(\boldsymbol{\xi}) &= \int_0^\theta K(\boldsymbol{x}(\tau), \boldsymbol{d}_\star(\tau))\,\mathrm{d}\tau + u(\boldsymbol{x}(\theta)) \\
&= \theta K(\boldsymbol{\xi}, \boldsymbol{d}_\star(0)) + \Big[u(\boldsymbol{\xi}) + \theta \nabla u(\boldsymbol{\xi}) \cdot \boldsymbol{f}(\boldsymbol{\xi}, \boldsymbol{d}_\star(0)) + \Big] + o(\theta); \\
0 &= \theta K(\boldsymbol{\xi}, \boldsymbol{d}_\star(0)) + \theta \nabla u(\boldsymbol{\xi}) \cdot \boldsymbol{f}(\boldsymbol{\xi}, \boldsymbol{d}_\star(0)) + o(\theta).
\end{aligned}
$$

Now dividing both sides by $\theta$ and sending $\theta$ to 0, we obtain

$$
0 = K(\boldsymbol{\xi}, \boldsymbol{d}_\star(0)) + \nabla u(\boldsymbol{\xi}) \cdot \boldsymbol{f}(\boldsymbol{\xi}, \boldsymbol{d}_\star(0)).
$$

Notice that the above equation involves $\boldsymbol{d}_\star(0)$ only. It is then natural to switch to a state-dependent optimal control in feedback form. The HJB equation for (3S.1.2) is then obtained by maximizing over $\boldsymbol{d} = \boldsymbol{d}_\star(0) \in \mathcal{D}$. By demanding the above equation holds for all $\boldsymbol{\xi} \in [0,1]^n \setminus \Delta$, the PDE can be written as:

$$
0 = \min_{\boldsymbol{d} \in \mathcal{D}} \Big\{ K(\boldsymbol{\xi}, \boldsymbol{d}) + \nabla u(\boldsymbol{\xi}) \cdot \boldsymbol{f}(\boldsymbol{\xi}, \boldsymbol{d}) \Big\}. \tag{3S.1.3}
$$

Recall from (1S.1.2) that Eq. (3S.1.1) for our Example 1 in component-wise form is

$$
\begin{cases}
\dot{q} = q(1-q)\left(\dfrac{b_v}{n+1}\left[\sum\limits_{k=0}^{n} p^k\right] - c\right), \\
\dot{p} = p(1-p)\left(\dfrac{b_a}{n+1} - (b_v - c)q - d\right).
\end{cases}
$$

It follows that Eq. (3S.1.3) in component-wise form for Example 1 is

$$
\min_{d \in [0, d_{\max}]} \Big[ \Big(1 - u_p p(1-p)\Big) d \Big] \;+\; u_q q(1-q)\Big(\frac{b_v}{n+1}\sum_{k=0}^{n} p^k - c\Big) + u_p p(1-p)\Big(\frac{b_a}{n+1} - q(b_v - c)\Big) + \delta \;=\; 0. \tag{3S.1.4}
$$

The linear dependence on $d$ of the minimized expression yields the *bang-bang* property:

$$
d_\star(q,p) = \begin{cases}
d_{\max}, & \text{if } \Big(1 - u_p p(1-p)\Big) < 0; \\
0, & \text{otherwise.}
\end{cases} \tag{3S.1.5}
$$

Similarly, Eq. (3S.1.1) for our Example 2 from (1S.2.2) in component-wise form is

$$
\begin{cases}
\dot{q} = (1-p)q(1-q)(g_{\mathrm{S}} - g_{\mathrm{R}}) + \beta C p q^2 (1-q) - \alpha q(1-q) d, \\
\dot{p} = p(1-p)[g_{\mathrm{S}} q + g_{\mathrm{R}}(1-q)] - \beta C p^2 q(1-q) - \alpha q p\, d.
\end{cases}
$$

Therefore, Eq. (3S.1.3) in component-wise form for Example 2 is

$$\min_{d\in[0,d_{\max}]}\Big[\Big(1-u_q\alpha q(1-q)-u_p\alpha qp\Big)d\Big] \;+\; u_q q(1-q)\Big[(1-p)(g_{\rm S}-g_{\rm R})+\beta Cqp\Big]+ \\ u_p\Big[p(1-p)\big(g_{\rm S}q+g_{\rm R}(1-q)\big)-\beta Cp^2q(1-q)\Big]+\delta \;=\; 0. \tag{3S.1.6}$$

Again, the linear dependence on $d$ yields the *bang-bang* property:

$$d_\star(q,p)=\begin{cases} d_{\max}, & \text{if } \Big(1-u_q\alpha q(1-q)-u_p\alpha qp\Big)<0;\\ 0, & \text{otherwise.}\end{cases} \tag{3S.1.7}$$

#### 3S.2 Derivation of the threshold-awareness HJB equation

Given the fixed reference probability space $\nu$ defined in §2S.1, we define the set of all *admissible* controls as

$$\mathcal{A}_\nu := \Big\{\boldsymbol{d}(\cdot):[0,\infty)\times\Omega\to\mathcal{D} \;\mid\; \boldsymbol{d}(\cdot) \text{ is } \mathcal{F}_t-\text{progressively measurable}\Big\}.$$

Ideally, we would want to consider controls in *feedback form*; i.e., $\boldsymbol{d}$ would be determined based on the current tumor configuration and perhaps the amount of drugs administered so far. But for technical reasons, we will first use *progressively measurable* open-loop controls, which we define below, to derive dynamic programming equations, and only then show that an optimal control can be found in feedback form.

To work under the framework of dynamic programming, we define a *value function* $v(\boldsymbol{\xi},\bar{s})$ encoding the maximal probability of reaching $\Delta_{\text{succ}}$ while keeping $\mathcal{J}$ from exceeding the threshold/budget value $\bar{s}$:

$$v(\boldsymbol{\xi},\bar{s})=\sup_{\boldsymbol{d}(\cdot)\in\mathcal{A}_\nu}\mathbb{P}\Big(\mathcal{J}(\boldsymbol{\xi},\boldsymbol{d}(\cdot))\le\bar{s}\Big). \tag{3S.2.1}$$

Any policy $\boldsymbol{d}_*(\cdot)$ is called optimal if $v(\boldsymbol{\xi},\bar{s})=\mathbb{P}\big(\mathcal{J}(\boldsymbol{\xi},\boldsymbol{d}_*(\cdot))\le\bar{s}\big)$.

Notice that $\mathbb{P}\big(\mathcal{J}(\boldsymbol{\xi},\boldsymbol{d}(\cdot))\le\bar{s}\big)$ can be treated as $\mathbb{E}\big[\mathbf{1}_{\{\mathcal{J}(\boldsymbol{\xi},\boldsymbol{d}(\cdot))\le\bar{s}\}}\big]$, where $\mathbf{1}_{\{Y\le y\}}$ is the indicator function such that it has value 1 if $Y\le y$ and 0 otherwise.

As explained in the main text, we introduce a new component of the state, the budget variable $s$. The (random) ODE describing its rate of change is

$$\dot{s}=-K(\boldsymbol{X}(t),d(t)), \qquad s(0)=\bar{s}.$$

Notice that $s(t)$ is strictly decreasing as time progresses. Thus, to ensure a nonnegative budget, we define a new terminal set

$$\widehat{\Delta} = \Big\{(x,s) \in [0,1]^n \times [0,\bar{\mathcal{S}}] \ \mid \ x \in \Delta \quad \text{or} \quad s = 0\Big\}$$

and correspondingly a new terminal time

$$\widehat{T} = \widehat{T}(\boldsymbol{\xi}, \bar{s}, \boldsymbol{d}(\cdot)) := \inf\Big\{t \in \mathbb{R}_+ \ \mid \ \Big(\boldsymbol{X}(t),\ s(t)\Big) \in \widehat{\Delta}, \quad \boldsymbol{X}(0) = \boldsymbol{\xi},\ s(0) = \bar{s}\Big\}.$$

We define

$$\Psi\Big(\boldsymbol{X}(\widehat{T})\Big) = \begin{cases} 1, \text{ if } \boldsymbol{X}(\widehat{T}) \in \Delta_{\text{succ}}, \\ 0, \text{ otherwise.} \end{cases}$$

as the last step of transforming our problem into a Mayer form [13].

Notice that regardless of random realizations, the largest possible $\widehat{T}$ is $\bar{s}/\delta$. In principle, we can recast it as a finite-horizon problem with $[0, \bar{s}/\delta]$ representing the horizon.

Now the original problem is equivalent to

$$v(\boldsymbol{\xi}, \bar{s}) = \max_{\boldsymbol{d}(\cdot)} \ \mathbb{E}^0\Big[\Psi(\boldsymbol{X}(\widehat{T}))\Big], \tag{3S.2.2}$$

where

$$\mathbb{E}^0\big[\ \cdot\ \big] = \mathbb{E}\big[\ \cdot\ \mid \text{Initial Conditions}\big].$$

This is now an exit-time problem in Mayer form over the domain $(\boldsymbol{\xi}, \bar{s}) \in [0,1]^n \times [0, \bar{\mathcal{S}}]$. We can therefore apply the stochastic dynamic programming principle [14] to derive the HJB PDE.

Assume the optimal $\boldsymbol{d}_*(\cdot)$ exists, and let $\widehat{T}_* = \widehat{T}(\boldsymbol{\xi}, \bar{s}, \boldsymbol{d}_*)$. The stochastic dynamic programming principle (Chapter V.2 in [14]) states for any (nonrandom) $\theta \in (0, \bar{s}/\delta)$,

$$v((\boldsymbol{\xi}, \bar{s}) = \mathbb{E}^0\left[v(\boldsymbol{X}(\widehat{T}_* \wedge \theta), s(\widehat{T}_* \wedge \theta))\right], \tag{3S.2.3}$$

where $a \wedge b = \min(a,b)$.

We now provide a formal derivation of the PDE that $v$ has to satisfy if it is sufficiently smooth. Notice that Eq. (3S.2.3) can be rewritten as

$$v((\boldsymbol{\xi}, \bar{s}) = \mathbb{E}^0\left[\mathbf{1}_{\{\widehat{T}_* < \theta\}} v(\boldsymbol{X}(\widehat{T}_*), s(\widehat{T}_*)) + \mathbf{1}_{\{\widehat{T}_* \geq \theta\}} v(\boldsymbol{X}(\theta), s(\theta))\right].$$

Seeking stochastic Taylor expansion of $v(\boldsymbol{X}(\theta), s(\theta))$ around $\theta = 0$, we have

$$v(\boldsymbol{X}(\theta), s(\theta)) - v((\boldsymbol{\xi}, \bar{s}) = \int_0^\theta \nabla v(\boldsymbol{X}(\tau), s(\tau)) \cdot \mathrm{d}\boldsymbol{X}(\tau) + \int_0^\theta \frac{\partial}{\partial s} v(\boldsymbol{X}(\tau), s(\tau))\, \mathrm{d}s(\tau) + \frac{1}{2} \sum_{i,j=1}^n \int_0^\theta \frac{\partial^2}{\partial \xi_i \partial \xi_j} v(\boldsymbol{X}(\tau), s(\tau)) \Big( \mathrm{d}X_i(\tau) \mathrm{d}X_j(\tau) \Big) + o(\theta).$$

Let $\boldsymbol{B} = \boldsymbol{\Sigma}\boldsymbol{\Sigma}^\top$. It follows that

$$\begin{aligned} &v(\boldsymbol{X}(\theta), s(\theta)) - v(\boldsymbol{\xi}, \bar{s}) \\ =& \int_0^\theta \nabla v(\boldsymbol{X}(\tau), s(\tau)) \cdot [\boldsymbol{a}(\boldsymbol{X}(\tau), \boldsymbol{d}_*(\tau))\, \mathrm{d}\tau + \boldsymbol{\Sigma}(\boldsymbol{X}(\tau), \boldsymbol{d}_*(\tau))\, \mathrm{d}\boldsymbol{W}(\tau)] \\ & - \int_0^\theta K(\boldsymbol{X}(\tau), \boldsymbol{d}_*(\tau)) \frac{\partial}{\partial s} v(\boldsymbol{X}(\tau), s(\tau))\, \mathrm{d}\tau + \frac{1}{2} \sum_{i,j=1}^n \int_0^\theta \frac{\partial^2}{\partial \xi_i \partial \xi_j} v(\boldsymbol{X}(\tau), s(\tau)) \boldsymbol{B}(\boldsymbol{X}(\tau), \boldsymbol{d}_*(\tau))_{i,j}\, \mathrm{d}\tau + o(\theta). \end{aligned}$$

Since Itô integrals with bounded functions have zero mean, we have

$$\begin{aligned} &\mathbb{E}^0 \left[ \mathbf{1}_{\{\widehat{T}_* \geq \theta\}} v(\boldsymbol{X}(\theta), s(\theta)) \right] \\ =& v(\boldsymbol{\xi}, \bar{s}) \mathbb{P}(\widehat{T}_* \geq \theta) + \mathbb{E}^0 \Bigg[ \mathbf{1}_{\{\widehat{T}_* \geq \theta\}} \Bigg\{ \int_0^\theta \nabla v(\boldsymbol{X}(\tau), s(\tau)) \cdot \boldsymbol{a}(\boldsymbol{X}(\tau), \boldsymbol{d}_*(\tau))\, \mathrm{d}\tau - \int_0^\theta K(\boldsymbol{X}(\tau), \boldsymbol{d}_*(\tau)) \frac{\partial}{\partial s} v(\boldsymbol{X}(\tau), s(\tau))\, \mathrm{d}\tau \\ & + \frac{1}{2} \sum_{i,j=1}^n \int_0^\theta \frac{\partial^2}{\partial \xi_i \partial \xi_j} v(\boldsymbol{X}(\tau), s(\tau)) \boldsymbol{B}(\boldsymbol{X}(\tau), \boldsymbol{d}_*(\tau))_{i,j}\, \mathrm{d}\tau + o(\theta) \Bigg\} \Bigg]. \end{aligned}$$

Therefore,

$$\begin{aligned} v(\boldsymbol{\xi}, \bar{s}) =& \mathbb{E}^0 \left[ \mathbf{1}_{\{\widehat{T}_* < \theta\}} v(\boldsymbol{X}(\widehat{T}), s(\widehat{T})) \right] + v(\boldsymbol{\xi}, \bar{s}) \mathbb{P}(\widehat{T}_* \geq \theta) \\ & + \mathbb{E}^0 \Bigg[ \mathbf{1}_{\{\widehat{T}_* \geq \theta\}} \Bigg\{ \int_0^\theta \nabla v(\boldsymbol{X}(\tau), s(\tau)) \cdot \boldsymbol{a}(\boldsymbol{X}(\tau), \boldsymbol{d}_*(\tau))\, \mathrm{d}\tau - \int_0^\theta K(\boldsymbol{X}(\tau), \boldsymbol{d}_*(\tau)) \frac{\partial}{\partial s} v(\boldsymbol{X}(\tau), s(\tau))\, \mathrm{d}\tau \\ & + \frac{1}{2} \sum_{i,j=1}^n \int_0^\theta \frac{\partial^2}{\partial \xi_i \partial \xi_j} v(\boldsymbol{X}(\tau), s(\tau)) \boldsymbol{B}(\boldsymbol{X}(\tau), \boldsymbol{d}_*(\tau))_{i,j}\, \mathrm{d}\tau \Bigg\} \Bigg] + o(\theta). \end{aligned}$$

Now dividing both sides by $\theta$ and sending $\theta \to 0$, we have

$$\begin{aligned} v(\boldsymbol{\xi}, \bar{s}) \lim_{\theta \to 0} \frac{1 - \mathbb{P}(\widehat{T}_* \geq \theta)}{\theta} =& \lim_{\theta \to 0} \mathbb{E}^0 \left[ \frac{\mathbf{1}_{\{\widehat{T}_* < \theta\}}}{\theta} v(\boldsymbol{X}(\widehat{T}_*), s(\widehat{T}_*)) \right] \\ & + \lim_{\theta \to 0} \mathbb{E}^0 \Bigg[ \frac{\mathbf{1}_{\{\widehat{T}_* \geq \theta\}}}{\theta} \Bigg\{ \int_0^\theta \nabla v(\boldsymbol{X}(\tau), s(\tau)) \cdot \boldsymbol{a}(\boldsymbol{X}(\tau), \boldsymbol{d}_*(\tau))\, \mathrm{d}\tau \\ & - \int_0^\theta K(\boldsymbol{X}(\tau), \boldsymbol{d}_*(\tau)) \frac{\partial}{\partial s} v(\boldsymbol{X}(\tau), s(\tau))\, \mathrm{d}\tau \\ & + \frac{1}{2} \sum_{i,j=1}^n \int_0^\theta \frac{\partial^2}{\partial \xi_i \partial \xi_j} v(\boldsymbol{X}(\tau), s(\tau)) \boldsymbol{B}(\boldsymbol{X}(\tau), \boldsymbol{d}_*(\tau))_{i,j}\, \mathrm{d}\tau \Bigg\} \Bigg] \end{aligned}$$

From [14, Chapter V], we know

$$\lim_{\theta\to 0}\frac{\mathbb{P}(\widehat{T}_*<\theta)}{\theta}=0,$$

and hence

$$v(\boldsymbol{\xi},\bar{s})\lim_{\theta\to 0}\frac{1-\mathbb{P}(\widehat{T}_*\geq\theta)}{\theta}=v(\boldsymbol{\xi},\bar{s})\lim_{\theta\to 0}\frac{\mathbb{P}(\widehat{T}_*<\theta)}{\theta}=0;$$

$$\lim_{\theta\to 0}\mathbb{E}^0\left[\frac{\mathbf{1}_{\{\widehat{T}_*<\theta\}}}{\theta}v(\boldsymbol{X}(\widehat{T}_*),s(\widehat{T}_*))\right]\leq\lim_{\theta\to 0}\frac{\mathbb{P}(\widehat{T}_*<\theta)}{\theta}=0.$$

Since $K$ and all components of $\boldsymbol{a}$ and $\boldsymbol{B}$ are bounded, $v$ is assumed to be sufficiently smooth, and $\mathbf{1}_{\{\widehat{T}\geq\theta\}}\to 1$ as $\theta\to 0$, by the dominated convergence theorem we have

$$0=\mathbb{E}^0\left[\nabla v(\boldsymbol{\xi},\bar{s})\cdot\boldsymbol{a}(\boldsymbol{\xi},\boldsymbol{d}_*(0,\omega))-\frac{\partial}{\partial s}v(\boldsymbol{\xi},\bar{s})K(\boldsymbol{\xi},\boldsymbol{d}_*(0,\omega))+\frac{1}{2}\sum_{i,j=1}^{n}\frac{\partial^2}{\partial\xi_i\partial\xi_j}v(\boldsymbol{\xi},\bar{s})\boldsymbol{B}(\boldsymbol{\xi},\boldsymbol{d}_*(0,\omega))_{i,j}\right]$$

Notice that the above equation only involves $\boldsymbol{d}_*(0)$ for every $\omega\in\Omega$. Namely, the starting optimal control value depends only on the initial state $(\boldsymbol{\xi},\bar{s})$ rather than a specific $\omega$. We then switch to a budget-dependent optimal control $\boldsymbol{d}_*(0)(\boldsymbol{\xi},\bar{s})$ in feedback form. The HJB equation for (3S.2.1) is then obtained by maximizing over $d\in[0,d_{\max}]$. By demanding the above equation holds for all $(\boldsymbol{\xi},\bar{s})\in([0,1]^n\setminus\Delta)\times[0,+\infty)$, the PDE can be written as:

$$0=\max_{\boldsymbol{d}\in\mathcal{D}}\left\{-\frac{\partial}{\partial s}v(\boldsymbol{\xi},s)K(\boldsymbol{\xi},\boldsymbol{d})+\nabla v(\boldsymbol{\xi},s)\cdot\boldsymbol{a}(\boldsymbol{\xi},\boldsymbol{d})+\frac{1}{2}\sum_{i,j=1}^{n}\frac{\partial^2}{\partial\xi_i\partial\xi_j}v(\boldsymbol{\xi},s)\boldsymbol{B}(\boldsymbol{\xi},\boldsymbol{d})_{i,j}\right\}.\tag{3S.2.4}$$

In both Example 1 and Example 2, we consider

$$K(\boldsymbol{X},d)=d+\delta,\ \text{ where } d\in[0,d_{\max}],$$
$$T=T(q_0,p_0,d(\cdot)):=\inf\Big\{t\in\mathbb{R}_+\ \mid\ \big(Q(t),P(t)\big)\in\Delta,\quad Q(0)=q_0,\ P(0)=p_0\Big\},$$

where $\Delta=\Big\{(q,p)\in[0,1]^2\ \mid\ p<\gamma_{\mathrm{r}}\quad\text{or}\quad p>\gamma_{\mathrm{f}}\Big\}$. The terminal function is defined as

$$g\Big(\boldsymbol{X}(T)\Big)=\begin{cases}+\infty, & \text{if } P(T)>\gamma_{\mathrm{f}},\\ 0, & \text{if } P(T)<\gamma_{\mathrm{r}}.\end{cases}$$

With the same extension to $\widehat{\Delta}$ and $\widehat{T}$, we have

$$\Psi\Big(Q(\widehat{T}), P(\widehat{T})\Big) = \begin{cases} 1, \text{ if } P(\widehat{T}) \leq \gamma_{\mathrm{r}}, \\ 0, \text{ otherwise.} \end{cases}$$

##### 3S.2.1 Specific formulations for Example 1

Recall from Box 5 in the main text and derivations for Eq. (2S.2.3) in §2S.2.2 that the components for Eq. (2S.1.1) are

$$\boldsymbol{X} = \begin{bmatrix} Q \\ P \end{bmatrix},$$

$$\boldsymbol{a}(\boldsymbol{X}, d) = \begin{bmatrix} Q(1-Q)\left(\dfrac{b_v}{n+1}\left[\sum\limits_{k=0}^{n} P^k\right] - c\right) + \Big[(1-Q)\sigma_2^2 - Q\sigma_3^2\Big] Q(1-Q) \\ P(1-P)\left(\dfrac{b_a}{n+1} - (b_v - c)Q - d\right) - \Big[\sigma_1^2 P - \sigma_2^2(1-P)(1-Q)^2 - \sigma_3^2(1-P)Q^2\Big] P(1-P) \end{bmatrix},$$

$$\boldsymbol{\Sigma}(\boldsymbol{X}, d) = \begin{bmatrix} 0 & -\sigma_2 Q(1-Q) & \sigma_3 Q(1-Q) \\ \sigma_1 P(1-P) & \sigma_2 P(1-P)(1-Q) & \sigma_3 P(1-P)Q \end{bmatrix}.$$

Thus, Eq. (3S.2.4) in component-wise form is

$$\begin{aligned} 0 = \max_{d\in[0,d_{\max}]} &\left\{ -\left[\frac{\partial v}{\partial p} p(1-p) + \frac{\partial v}{\partial s}\right] d \right\} - \delta \frac{\partial v}{\partial s} \\ &+ \frac{\partial v}{\partial q}\left[\left(\frac{b_v}{n+1}\sum_{k=0}^{n} p^k - c\right) - q\sigma_3^2 + (1-q)\sigma_2^2\right] q(1-q) \\ &+ \frac{\partial v}{\partial p}\left[\left(\frac{b_a}{n+1} - q(b_v - c)\right) - [\sigma_1^2 p - \sigma_2^2(1-p)(1-q)^2 - \sigma_3^2(1-p)q^2]\right] p(1-p) \\ &+ \frac{1}{2}\frac{\partial^2 v}{\partial q^2}\left[q^2(1-q)^2(\sigma_2^2 + \sigma_3^2)\right] + \frac{1}{2}\frac{\partial^2 v}{\partial p^2}\left[\sigma_1^2 + (1-q)^2\sigma_2^2 + q^2\sigma_3^2\right] p^2(1-p)^2 \\ &+ \frac{\partial^2 v}{\partial q \partial p}\left[q\sigma_3^2 - (1-q)\sigma_2^2\right] pq(1-p)(1-q). \end{aligned} \quad (3S.2.5)$$

With degenerate parabolicity present in the problem, the value function $v$ does not have to be smooth, and there may not be a *classical solution* to the above PDE. Thus, $v$ has to be interpreted as a (possibly discontinuous) *viscosity solution* of this equation [1, Chapter 5].

The linear dependence on $d$ yields the *bang-bang* property:

$$d_*(q,p,s) = \begin{cases} d_{\max}, & \text{if } \left(\dfrac{\partial v}{\partial p} p(1-p) + \dfrac{\partial v}{\partial s}\right) < 0, \\ 0, & \text{otherwise.} \end{cases} \tag{3S.2.6}$$

##### 3S.2.2 Specific formulations for Example 2

Recall from Box 6 in the main text and derivations for Eq. (2S.3.1) in §2S.3 that the components for Eq. (2S.1.1) are

$$\boldsymbol{X} = \begin{bmatrix} Q \\ P \end{bmatrix} := \begin{bmatrix} \dfrac{Z_{\text{S}}}{Z_{\text{S}} + mZ_{\text{R}}} \\ Z_{\text{S}} + mZ_{\text{R}} \end{bmatrix}, \quad \boldsymbol{W} = \begin{bmatrix} B_t \end{bmatrix} \text{ (where } B_t \text{ is a standard 1D Brownian motion)},$$

$$\boldsymbol{a}(\boldsymbol{X}, d) = \begin{bmatrix} Q(1-Q)\Big\{(1-P)(g_{\text{S}} - g_{\text{R}}) - \alpha d + \beta CQP + (1-P)^2[\sigma_{\text{R}}^2(1-Q) - \sigma_{\text{S}}^2 Q + \sigma_{\text{S}}\sigma_{\text{R}}]\Big\} \\ P(1-P)(g_{\text{S}}Q + g_{\text{R}}(1-Q)) - \alpha QPd - \beta CP^2Q(1-Q) \end{bmatrix},$$

$$\boldsymbol{\Sigma}(\boldsymbol{X}, d) = \begin{bmatrix} (1-P)Q(1-Q)(\sigma_{\text{S}} - \sigma_{\text{R}}) \\ P(1-P)[\sigma_{\text{S}}Q + \sigma_{\text{R}}(1-Q)] \end{bmatrix}.$$

Thus, Eq. (3S.2.4) in component-wise form is

$$\begin{aligned} 0 = \max_{d \in [0, d_{\max}]} &\left\{ -\left[\frac{\partial v}{\partial q}\alpha q(1-q) + \frac{\partial v}{\partial p}\alpha qp + \frac{\partial v}{\partial s}\right] d \right\} - \delta \frac{\partial v}{\partial s} \\ &+ \frac{\partial v}{\partial q}\left[(1-p)(g_{\text{S}} - g_{\text{R}}) + \beta Cpq + (1-p)^2\{\sigma_{\text{R}}^2(1-q) - \sigma_{\text{S}}^2 q + \sigma_{\text{S}}\sigma_{\text{R}}\}\right] q(1-q) \\ &+ \frac{\partial v}{\partial p}\left[(1-p)[g_{\text{S}}q + g_{\text{R}}(1-q)] - \beta Cpq(1-q)\right] p \\ &+ \frac{1}{2}\frac{\partial^2 v}{\partial q^2}(1-p)^2 q^2(1-q)^2(\sigma_{\text{S}} - \sigma_{\text{R}})^2 + \frac{1}{2}\frac{\partial^2 v}{\partial p^2} p^2(1-p)^2[\sigma_{\text{S}}q + \sigma_{\text{R}}(1-q)]^2 \\ &+ \frac{\partial^2 v}{\partial q \partial p}(\sigma_{\text{S}} - \sigma_{\text{R}})(1-p)^2 pq(1-q)[\sigma_{\text{S}}q + \sigma_{\text{R}}(1-q)]. \end{aligned} \tag{3S.2.7}$$

Again, the linear dependence on $d$ yields the *bang-bang* property:

$$d_*(q,p,s) = \begin{cases} d_{\max}, & \text{if } \left(\dfrac{\partial v}{\partial q}\alpha q(1-q) + \dfrac{\partial v}{\partial p}\alpha qp + \dfrac{\partial v}{\partial s}\right) < 0, \\ 0, & \text{otherwise.} \end{cases} \tag{3S.2.8}$$

### 4S Numerical methods and implementation details

We provide the numerical schemes and implementation details of: (i) solving for $v(\boldsymbol{\xi}, s)$; (ii) solving for $u(q,p)$; and (iii) generating CDFs in §4S.1, §4S.2, and §4S.3 respectively. In both (i) and (ii), the optimal policy is found by numerically solving the corresponding HJB equation.

#### 4S.1 Threshold-aware optimal case

We approximate the solution to (3S.2.4) by a first-order accurate semi-Lagrangian discretization [11] over the $(\boldsymbol{\xi}, s)$ space in standard $(n+1)$-dimensional Cartesian coordinates. Since we are really only interested in the expected value, it suffices to consider weak approximations [12, 21] of (2S.1.1). For any small $\tau > 0$, a first order weak approximation of $\boldsymbol{X}^{\tau,\boldsymbol{d}} \approx \boldsymbol{X}(\tau; \boldsymbol{d})$ starting from $\boldsymbol{X}(0) = \boldsymbol{\xi}$ with any control value $d \in \mathcal{D}$ is

$$\boldsymbol{X}^{\tau,\boldsymbol{d}} = \boldsymbol{\xi} + \tau \boldsymbol{a}(\boldsymbol{\xi}, \boldsymbol{d}) + \boldsymbol{\Sigma}(\boldsymbol{\xi}, \boldsymbol{d}) \Delta \boldsymbol{W}^{\tau},$$

where $\Delta \boldsymbol{W}^{\tau} = (\Delta W_1^{\tau}, \dots, \Delta W_m^{\tau})$ with $\Delta W_j^{\tau}$ representing a two-point distributed variable with the distribution

$$\mathbb{P}\left(\Delta W_j^{\tau} = \pm\sqrt{\tau}\right) = \frac{1}{2}.$$

Recall $\boldsymbol{W}$ is a standard $m$-dimensional Brownian motion. It follows from above that $\boldsymbol{X}^{\tau,\boldsymbol{d}}$ will have $2^m$ possible locations to "land" with equal probability. Let $\tilde{\boldsymbol{X}}_{\ell}^{\boldsymbol{d}}$, $\ell \in \{1, 2, \dots, 2^m\}$, denote the $2^m$ possible locations of $\boldsymbol{X}^{\tau,\boldsymbol{d}}$. Assuming that $\mu_{j,\ell}$ is the $j$-th bit in a binary representation of $(\ell - 1)$ and $\Delta \boldsymbol{W}_{\ell}^{\tau} = (\Delta W_{1,\ell}^{\tau}, \dots, \Delta W_{m,\ell}^{\tau})$ with $\Delta W_{j,\ell}^{\tau} := (-1)^{\mu_{j,\ell}} \sqrt{\tau}$, we can now express these possible locations explicitly as

$$\boldsymbol{X}_{\ell}^{\tau,\boldsymbol{d}} = \boldsymbol{\xi} + \tau \boldsymbol{a}(\boldsymbol{\xi}, \boldsymbol{d}) + \boldsymbol{\Sigma}(\boldsymbol{\xi}, \boldsymbol{d}) \Delta \boldsymbol{W}_{\ell}^{\tau}.$$

Assume the running cost is constant over $\tau$ units of time. The dynamic programming equation obtained by weak approximations is then

$$v(\boldsymbol{\xi}, \bar{s}) = \max_{\boldsymbol{d} \in \mathcal{D}} \left\{ \frac{1}{2^m} \sum_{\ell=1}^{2^m} v\left( \tilde{\boldsymbol{X}}_{\ell}^{\boldsymbol{d}}, \bar{s} - \tau K(\boldsymbol{\xi}, \boldsymbol{d}) \right) \right\} + o(\tau). \tag{4S.1.1}$$

To solve Eq. (4S.1.1) numerically, we discretize $[0, \bar{\mathcal{S}}]$ into $(M+1)$ equidistant slices, with $s_k = k\Delta s$ for $k = 0, 1, 2, \dots, M$. Each $s$-slice consists of a uniform rectangular grid on $[0,1]^n$, with $\boldsymbol{x}_i = i\Delta x$ for a multi-index $i = (i_1, i_2, \dots, i_n)$ where $i_j = 0, 1, 2, \dots, N_x$ for all $j \in \{1, 2, \dots, n\}$.

Suppose starting from a gridpoint $\boldsymbol{x}_i$ on an $s_k$-slice, We have

$$v(\boldsymbol{x}_i, s_k) = \max_{\boldsymbol{d}\in\mathcal{D}} \left\{ \frac{1}{2^m} \sum_{\ell=1}^{2^m} v\Big(\tilde{\boldsymbol{x}}^{\boldsymbol{d}}_{i,\ell},\; s_k - \tau K(\boldsymbol{x}_i, \boldsymbol{d})\Big) \right\} + o(\tau),$$

where $\tilde{\boldsymbol{x}}^{\boldsymbol{d}}_{i,\ell} := \boldsymbol{x}_i + \tau\; \boldsymbol{a}(\boldsymbol{x}_i, \boldsymbol{d}) + \boldsymbol{\Sigma}(\boldsymbol{x}_i, \boldsymbol{d})\Delta \boldsymbol{W}^{\tau}_{\ell}$. We choose an explicit "$s$-marching" scheme, in which the causality is maintained in threshold variable $s$ since it is strictly decreasing along a path. By "causality", we mean the scheme uses the values from the previous $s$-slice (known) to approximate the values for the current $s$-slice (unknown). Namely, the above equation is exactly

$$v(\boldsymbol{x}_i, s_k) = \max_{\boldsymbol{d}\in\mathcal{D}} \left\{ \frac{1}{2^m} \sum_{\ell=1}^{2^m} v\Big(\tilde{\boldsymbol{x}}^{\boldsymbol{d}}_{i,\ell}, s_{k-1}\Big) \right\} + o(\tau). \tag{4S.1.2}$$

As a result, $\tau$ must be chosen so that $\tau K(\boldsymbol{x}_i, \boldsymbol{d}) = \Delta s$, which makes it necessary to use a control-dependent $\tau_d$, yielding $\tilde{\boldsymbol{x}}^{\boldsymbol{d}}_{i,\ell} := \boldsymbol{x}_i + \tau_{\boldsymbol{d}}\; \boldsymbol{a}(\boldsymbol{x}_i, \boldsymbol{d}) + \boldsymbol{\Sigma}(\boldsymbol{x}_i, \boldsymbol{d})\Delta \boldsymbol{W}^{\tau_{\boldsymbol{d}}}_{\ell}$. Let $V^k_i \approx v(\boldsymbol{x}_i, s_k)$ denote the discretized approximate solution at $(\boldsymbol{x}_i, s_k)$ and $\widetilde{V}^{k-1,\boldsymbol{d}}_{i,\ell} \approx v(\tilde{\boldsymbol{x}}^{\boldsymbol{d}}_{i,\ell}, s_{k-1})$. The fully discretized equation is then

$$V^k_i = \max_{\boldsymbol{d}\in\mathcal{D}} \left\{ \frac{1}{2^m} \sum_{\ell=1}^{2^m} \widetilde{V}^{k-1,\boldsymbol{d}}_{i,\ell} \right\}. \tag{4S.1.3}$$

Let $D^k_i \approx \boldsymbol{d}_*(\boldsymbol{x}_i, s_k)$ denote the discretized approximate optimal feedback policy at $(\boldsymbol{x}_i, s_k)$. We note that it is constructed as a by-product of solving (4S.1.3) while determining the maximizing $d$ at each gridpoint.

It is worth noting that, regardless of $\boldsymbol{d}$ value, $\tilde{\boldsymbol{x}}^{\boldsymbol{d}}_{i,\ell}$, $\ell \in \{1, 2, \ldots, 2^m\}$, is generally not a gridpoint on the $s_{k-1}$-slice. Thus, each $\widetilde{V}^{k-1,\boldsymbol{d}}_{i,\ell}$ is computed by interpolating the values from the neighboring gridpoints of $\tilde{\boldsymbol{x}}^{\boldsymbol{d}}_{i,\ell}$, which is the essence of semi-Lagrangian schemes. In our implementation, we have chosen the fourth-order accurate ENO cubic interpolation [12, 23] to reduce the numerical diffusion. Our full method is summarized in Algorithm S1 where $\Xi = \{i\Delta x \mid i = (i_1, i_2, \ldots, i_n) \text{ where } i_j = 0, 1, 2, \ldots, N_x \; \forall j \in \{1, 2, \ldots, n\}\}$.

In accordance with [12], $\Delta x = o(\Delta s^{1/r})$ is needed to guarantee the convergence of this numerical scheme to the viscosity solution. Here, $\Delta x = \Delta x_j$ for all $j \in \{1, 2, \ldots, n\}$ is the space discretization step and $r = m + 1$ with $m$ being the order of the interpolating polynomial. Since we use cubic ENO interpolants, this requires $\Delta x = o(\Delta s^{1/4})$.

To obtain accurate optimal threshold-aware policies and optimal probability of success shown in Figures 3&6 of the main text, we have used $N = 1600$ on each side of the $qp$ square and $M = 6000$ slices along the positive $s$-axis in Example 1. We use $N = 3200$ and $M = 48000$ in Example 2. With these choices of $N$, the success and failure barriers are just grid lines (recall that we choose $\gamma_{\mathrm{r}} = 1 - \gamma_{\mathrm{f}} = 10^{-2}$), and it is easy to apply the boundary conditions.

Additionally, since we have causality in $s$ variable, we have further taken advantage of loop parallelism on the inner loops (iterating over spatial variables $q$ and $p$) with the aid of `OpenMP` in `C++`.

For visualization purposes, for all figures related to Example 1, we transform the approximate solution $V$ on each

**Algorithm S1:** Threshold(risk)-aware value function/policy computation

```
Initialize V, D at s = 0 using the initial & boundary conditions;
for s_k = kΔs, k = 1,...M do
    for every x_i ∈ Ξ do
        if x_i ∈ Δ_succ then
            V_i^k ← 1;
        else if x_i ∈ Δ_fail then
            V_i^k ← 0;
        else
            for every d ∈ D do
                for ℓ = 1, 2, ..., 2^m do
                    x̃_{i,ℓ}^d ← x_i + τ_d a(x_i, d) + Σ(x_i, d) ΔW_ℓ^{τ_d} ;
                    Compute Ṽ_{i,ℓ}^{k-1,d} by interpolation;
                V̄_i^{k,d} ← (1/2^m) Σ_{ℓ=1}^{2^m} Ṽ_{i,ℓ}^{k-1,d};
            V_i^k ← max_d { V̄_i^{k,d} };
            D_i^k ← arg max_d { V̄_i^{k,d} };
```

$s$-slice from the $qp$ square into the GLY-DEF-VOP triangle. See Figure 8 for the specific geometric transformation.

**Remark:** For both of our examples in the main text, $\mathcal{D} = \{0, d_{\max}\}$ and $K(\boldsymbol{x}, d) = d + \delta$. Thus, the $d$-dependent $\tau$ has only two possible values

$$\begin{cases} \tau_{\{d=0\}} = \dfrac{\Delta s}{\delta}, \\ \tau_{\{d=d_{\max}\}} = \dfrac{\Delta s}{d_{\max} + \delta}, \end{cases} \tag{4S.1.4}$$

and we can significantly improve the computational efficiency by pre-computing all $\tilde{\boldsymbol{x}}_{i,l}^{d}$.

We note that the disparity in $d$-dependent $\tau$ values described in (4S.1.4) leads to much larger spatial steps with $d = 0$. This does not prevent the convergence under the grid refinement, but does contribute to larger local truncation errors on a fixed grid. In the future we hope to investigate alternative $s$-implicit semi-Lagrangian discretizations, which avoid such $\tau$-disparities, but make equations coupled in each $s$-slice. A similar approach has already proven to be advantageous in solving first-order HJB equations with rapid transition layers [24].

**Remark II:** Recall the random ODE satisfied by the budget variable $s$ :

$$\frac{\mathrm{d}s}{\mathrm{d}t} = -K(\boldsymbol{x}(t), \boldsymbol{d}(t)), \qquad s(0) = \bar{s}.$$

If we rescale the time as $\tau = t/L$ (for some fixed $L > 0$) and denote $\tilde{s}(\tau) = s(t) = s(\tau L)$, we obtain the following

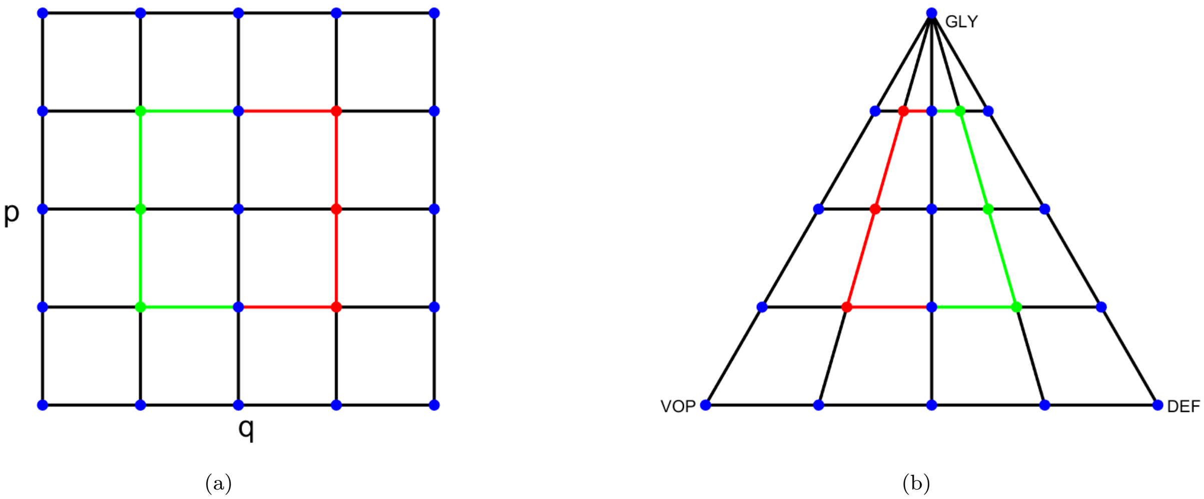


**Figure 8: Geometric transformation from the $qp$ squre to the GLY-DEF-VOP triangle in Cartesian coordinates:** (a) grid on $qp$ square; (b) grid on GLY-DEF-VOP triangle. See how the shape and the colors of boundary of the region enclosed by *green-red* lines change.

relation

$$\begin{aligned}\frac{\mathrm{d}\tilde{s}}{\mathrm{d}\tau} &= -LK(\tau)\\ \tilde{s}(\tau) &= \bar{s} - \int_0^\tau LK(\theta)\mathrm{d}\theta\\ &= L\left(\frac{\bar{s}}{L} - \int_0^\tau K(\theta)\mathrm{d}\theta\right)\end{aligned}$$

Consequently, the solution $v(\xi, s)$ on $s \in [0, \bar{\mathcal{S}}]$ is equivalent to $v(\xi, \tilde{s})$ on $\tilde{s} \in [0, \bar{\mathcal{S}}/L]$ with $v(\xi, s) = v(\xi, \tilde{s}L)$ for all $s$ and $\tilde{s}$. This enables us to reduce the size of $\bar{\mathcal{S}}$ if it is too large when implementing Algorithm S1 and makes it easier to fine-tune the value of $\Delta s$. In our implementations, we have chosen $L = 1$ for Example 1 and $L = 15$ for Example 2.

#### 4S.2 Deterministic-optimal case

In this section, we describe a numerical scheme for approximating a two-dimensional value function $u(q, p)$ in deterministic optimal control problems. This scheme has been applied to both Example 1 and Example 2 to solve (3S.1.4) and (3S.1.6). We use a first-order accurate semi-Lagrangian discretization [11] over the $(q, p)$ plane in standard two-dimensional Cartesian coordinates. We again discretize $(q, p) \in [0, 1]^2$ using a uniform $(N + 1) \times (N + 1)$ rectangular grid, on which the value function is approximated by $U_{i,j} \approx u(q_i, p_j)$.

Note that the semi-Lagrangian discretization presented here is similar to (but not quite the same as) the one used in [16]. In particular, Gluzman et al. used a *linear* interpolation on a *triangular* mesh with *adaptive* time step $\tau$, while our current version uses a *bi-linear* interpolation on a *rectangular* grid with a *fixed* time step $\tau$. While different, these two choices have the same formal order of accuracy and converge to the same result as the discretization parameters approach zero.

To be more compact, we further denote the deterministic dynamics (for both Example 1 and Example 2) as

$$\begin{bmatrix} \dot{q} \\ \dot{p} \end{bmatrix} = \begin{bmatrix} f_q(q,p,d) \\ f_p(q,p,d) \end{bmatrix} \tag{4S.2.1}$$

Assuming the rate of change is constant for a small amount of time $\tau > 0$, the foot of the characteristics starting from a gridpoint $(q_i, p_j)$ lands at a new state

$$\tilde{q}_i^d = q_i + \tau f_q(q_i, p_j, d),$$
$$\tilde{p}_j^d = p_j + \tau f_p(q_i, p_j, d).$$

From Bellman's Optimality Principle [2], we have

$$u(q_i, p_j) = \min_{dd \in \{0,\, d_{\max}\}} \left\{ \tau(d+\delta) + u\left(\tilde{q}_i^d, \tilde{p}_j^d\right) \right\} + o(\tau), \tag{4S.2.2}$$

yielding the discretized version

$$U_{i,j} = \min_{d \in \{0,\, d_{\max}\}} \left\{ \tau(d+\delta) + \widetilde{U}_{i,j}^d \right\}, \tag{4S.2.3}$$

where $\widetilde{U}_{i,j}^d \approx u(\tilde{q}_i^d, \tilde{p}_j^d)$ is computed through a *bi-linearly* interpolation of the $U$ values from the four neighboring gridpoints surrounding $(\tilde{q}_i^d, \tilde{p}_j^d)$. The optimal feedback policy is recovered as an argmin in (4S.2.3).

Since the discretization (4S.2.3) is not explicitly causal, an iterative method is needed to solve the resulting system. This is often done through standard Value Iterations (VI) [3–5], but in our case this approach results in a very slow convergence on a fixed grid. To address this, we have implemented a version of the "*hybrid Value-Policy*" Iteration (VPI) algorithm [18].

That is, we start with value iterations where we solve the nonlinear Eq. (4S.2.3) by Gauss-Seidel iterations with "Fast Sweeping" orderings [7, 22, 25]. We initialize `err` to be the $L_\infty$-norm of $U$-change in the very first value iteration, and use $\epsilon$ to denote the $L_\infty$-norm of $U$-change in the *last* value iteration performed so far. Whenever $\epsilon$ falls below $\rho *$ `err`, where $\rho \in (0,1)$ is a preset hyperparameter, we store this $\epsilon$ as the new `err` and proceed to the "*policy-evaluation*" (PE) step.

In the PE step, we compute the value function by solving a system of linear equations with a fixed policy $\hat{d}$ (recovered from the most recent value iteration). We thus solve a linear system of equations

$$U_{i,j} = \tau\big(\widehat{D}_{i,j} + \delta\big) + \widetilde{U}^{\widehat{D}_{i,j}}, \tag{4S.2.4}$$

where $\widehat{D}_{i,j} = \hat{d}(q_i, p_j)$ and $\widetilde{U}^{\widehat{D}_{i,j}} \approx u\left(\tilde{q}_i, \tilde{p}_j^{\widehat{D}_{i,j}}\right)$ is again computed through a bi-linear interpolation.

After obtaining the solution to Eq. (4S.2.4), we return to the value iteration part and repeat the process until $\epsilon <$ `tol`, where `tol` is a preset tolerance of convergence. In both Example 1 and Example 2, to obtain an accurate deterministic-optimal policy, we have used $N = 3200$ on each side of the unit $qp$-square, `tol` $= 10^{-6}$, and $\rho = 0.7$.

#### 4S.3 Generating CDFs

To generate the CDFs we provide in both the main text and §5S.3, we have used a Monte Carlo (MC) method with $10^5$ samples. That is, for each sample, we start with a fixed initial tumor configuration $(q_0, p_0)$, and then apply a strong approximation [21] of $(Q(t), P(t))$ to simulate a sample path till either stabilization/remission or death. We store the random cost $\mathcal{J}$ incurred along each sample path as a data point, and then generate the (empirical) CDF based on $10^5$ data points. In particular, we compute the Kaplan-Meier estimate [19] (empirical) CDF by `Matlab`'s built-in function `ecdf()`.

Let $\Delta t$ denote the uniform time step. The cost incurred for each time step along the sample path is

$$\Delta J = \begin{cases} \Big(d_{\max} + \delta\Big)\Delta t, & \text{if } d_* = d_{\max} \text{ at this step,} \\ \delta\,\Delta t, & \text{otherwise.} \end{cases}$$

Thus, the cost accumulated at the $n$-th time step can be defined recursively as

$$J^n = J^{n-1} + \Delta J, \qquad J_0 = 0.$$

Suppose we start with an initial budget value $\bar{s}$ at $t = 0$. Then the remaining budget at the $n$-th time step is

$$S^n = S^{n-1} - \Delta J, \qquad S_0 = \bar{s}.$$

As mentioned in §4S.1, we discretize $[0, \bar{s}]$ into equidistant slices, and $(q, p) \in [0, 1]^2$ into a uniform rectangular grid. Among many choices of strong approximations of SDEs, we choose the Euler-Maruyama scheme [21].

Let us denote the approximate solution to (2S.2.3) (or (2S.3.1)) at the $n$-th time step ($t_n = n\Delta t$) as $(Q^n, P^n) \approx (Q(t_n), P(t_n))$, and the optimal feedback policy at $(Q^n, P^n, S^n)$ as $D^n$. Then the Euler-Maruyama scheme defines

$(Q^n, P^n)$ recursively by

$$\begin{bmatrix} Q^n \\ P^n \end{bmatrix} = \begin{bmatrix} Q^{n-1} \\ P^{n-1} \end{bmatrix} + \boldsymbol{a}(Q^{n-1}, P^{n-1}, D^{n-1})\Delta t + \boldsymbol{\Sigma}(Q^{n-1}, P^{n-1})\Delta W^n. \tag{4S.3.1}$$

where $(Q_0, P_0) = (q_0, p_0)$. Unlike in the weak approximation mentioned in §4S.1, this time the $\Delta W_j^n$'s are *independent and identically distributed* (i.i.d.) normal random variables with mean zero and variance $\Delta t$ for all $j \in \{1, 2, \dots, m\}$.

Due to memory limitations, we only store the policy for a subsample of $s$-slices represented in the PDE discretization grid. The policy is stored for $s = 0$, $\Delta\hat{s}$, $2\Delta\hat{s}$, $3\Delta\hat{s}, \ldots$ (We use $\Delta\hat{s} = 0.005$ for Example 1 and $\Delta\hat{s} = 0.00125$ for Example 2). To obtain an accurate sample path approximation, we need a sufficiently small $\Delta t$, which often means that $(Q^n, P^n, S^n)$ is not on any stored $s$-slice. As a consequence, we have to determine whether to use drugs or not based on the *data cube* surrounding $(Q^n, P^n, S^n)$ (the cube is formed by the 4 data points on the $s$-slice above $(Q^n, P^n, S^n)$ and another 4 data points on the $s$-slice below it).

For all the figures related to Example 1 we provide in the main text and SM, we choose a "conservative" drugs-on/off determination strategy for Example 1. That is, we decide to use drugs at $(Q^n, P^n, S^n)$ if all of the 8 data points of the cube have the value $d_* = d_{\max}$. On the other hand, we used a "Majority" strategy for Example 2.[I] That is, we decide to use drugs at $(Q^n, P^n, S^n)$ if 5 of the 8 data points of the cube have the value $d_* = d_{\max}$. When the budget runs out, i.e., $S_n = 0$, we switch to the deterministic-optimal policy as mentioned in §3.1 of the main text. Notice that the cost will not stop accumulating until we cross either $\gamma_{\mathrm{r}}$ or $\gamma_{\mathrm{f}}$. In the deterministic-optimal case, we only need to consider the *data square* surrounding $(Q^n, P^n)$ since it is now $s$-independent.

In our implementations, we have used the following adaptive choice of $\Delta t$:

$$\text{Example 1: } \Delta t = \begin{cases} 2.5 \times 10^{-3}, & \text{if } d_* = 0, \\ 1.64 \times 10^{-4}, & \text{if } d_* = d_{\max}. \end{cases} \qquad \text{Example 2: } \Delta t = \begin{cases} 2.5 \times 10^{-3}, & \text{if } d_* = 0, \\ 4.10 \times 10^{-4}, & \text{if } d_* = d_{\max}. \end{cases}$$

A parallel `for` loop was used in `Matlab` to reduce the computational time of Monte Carlo simulations.

Here, we provide a table of details of all CDF figures in both the main text and SM. The "95% Confidence bounds of `ecdf`()" is computed via Greenwood's formula [17].

For Example 1, the discrepancy between the approximated value function and the MC simulation at a given $(q_0, p_0, \bar{s})$ is around $10^{-3}$ except for two examples where the discrepancy is around 0.01. For Example 2, the discrepancy between the approximated value function and the MC simulation at a given $(q_0, p_0, \bar{s})$ is around $10^{-4}$ except for

[I] Alternatively, one can also try define an "aggressive" strategy in either example; i.e., to use drugs as long as one of the 8 data points has $d_* = d_{\max}$. We have considered this and other determination strategies and they all yield results within the 95% confidence interval of each other.

one example where the discrepancy is around $10^{-3}$. These discrepancies are due to (a) the discretization errors in solving the Hamilton-Jacobi PDE, (b) subsampling of the optimal policy, and (c) the variability of outcomes and a time discretization in MC simulations. All of these can be reduced with a higher computational cost (e.g., using finer discretizations and more MC simulations).

| **Example 1** | **Initial configuration** $(q_0, p_0, \bar{s})$ | **Volatilities** $(\sigma_1 = \sigma_2 = \sigma_3)$ | **Value function** $v(q_0, p_0, \bar{s})$ | **Empirical CDF at** $\bar{s}$ | **95% Confidence bounds of** `ecdf()` **at** $\bar{s}$ |
|---|---|---|---|---|---|
| Figure 1(c) | $(0.26, 0.665, 5.0)$ | 0.15 | 0.6763 | 0.6737 | $[0.6708, 0.6766]$ |
| Figure 1(c) | $(0.26, 0.665, 4.5)$ | 0.15 | 0.4117 | 0.4041 | $[0.4011, 0.4072]$ |
| Figure 5(c) | $(0.27, 0.4, 4.71)$ | 0.15 | 0.6421 | 0.6371 | $[0.6341, 0.6401]$ |
| Figure 5(c) | $(0.27, 0.4, 4.35)$ | 0.15 | 0.4684 | 0.4562 | $[0.4532, 0.4593]$ |
| Figure 9(c) | $(0.8, 0.4, 3.77)$ | 0.15 | 0.6481 | 0.6503 | $[0.6474, 0.6533]$ |
| Figure 9(c) | $(0.8, 0.4, 3.45)$ | 0.15 | 0.3238 | 0.3145 | $[0.3116, 0.3173]$ |
| Figure 12(e) | $(0.27, 0.4, 4.46)$ | 0.5 | 0.8906 | 0.8936 | $[0.8917, 0.8956]$ |
| Figure 12(e) | $(0.27, 0.4, 3.65)$ | 0.5 | 0.7742 | 0.7761 | $[0.7735, 0.7787]$ |

| **Example 2** | **Initial configuration** $(q_0, p_0, \bar{s})$ | **Volatilities** $(\sigma_S = \sigma_R)$ | **Value function** $v(q_0, p_0, \bar{s})$ | **Empirical CDF at** $\bar{s}$ | **95% Confidence bounds of** `ecdf()` **at** $\bar{s}$ |
|---|---|---|---|---|---|
| Figure 7(c) | $(0.45, 0.9, 69.45)$ | 0.15 | 0.6746 | 0.6739 | $[0.6710, 0.6768]$ |
| Figure 7(c) | $(0.45, 0.9, 60)$ | 0.15 | 0.4013 | 0.3984 | $[0.3954, 0.4015]$ |
| Figure 10(c) | $(0.55, 0.9, 72.75)$ | 0.15 | 0.6697 | 0.6693 | $[0.6664, 0.6722]$ |
| Figure 10(c) | $(0.55, 0.9, 60)$ | 0.15 | 0.3040 | 0.3039 | $[0.3010, 0.3067]$ |

### 5S More numerical results

#### 5S.1 Policies, trajectories, and CDFs for the EGT-based model (additional example)

Using the same model and parameter values as in the main text, we now consider another initial tumor configuration at $(q_0, p_0) = (0.8, 0.4)$ located in the blue (drugs-off) region of $d_\star$.

Our threshold-aware policy (pink) still improves the $\mathbb{P}(\mathcal{J} \leq \bar{s}_{\text{med}})$ from 50% to 65%, where $\bar{s}_{\text{med}} = 3.77$ is the median cost of $\mathcal{J}$ associated with $d_\star$. When starting from a lower initial budget $\bar{s} = 3.45$, the deterministic-optimal policy provides only a 15.6% chance of stabilization, while our threshold-aware policy (orange) doubles this $\mathbb{P}(\mathcal{J} \leq 3.45)$ to 31.5%.

We can see from Figure 9(a) that the deterministic-optimal policy basically prescribes drugs till stabilization once

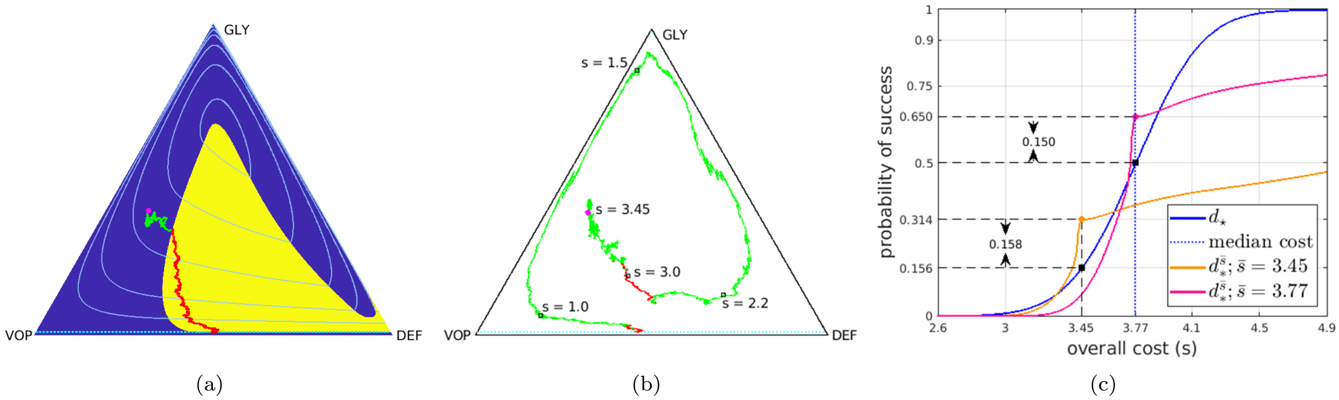


**Figure 9: Comparison between threshold-aware policies and the deterministic-optimal policy (EGT model; Case II).** Starting from an initial state $(q_0, p_0) = (0.8, 0.4)$ (magenta dot): (a) a sample path with cost 3.94 under the deterministic-optimal policy; (b) a sample path starting at $\bar{s} = 3.45$ with a total cost of 3.41 under the (orange) threshold-aware policy; (c) CDFs of the cumulative cost $\mathcal{J}$ approximated using $10^5$ random simulations. In (c), the *solid blue* curve is the CDF generated with the deterministic-optimal policy. Its median (*dashed blue* line) is 3.77 and its mean conditioning on success is also 3.77. The *solid orange* curve is the CDF generated with the threshold-aware policy with $\bar{s} = 3.45$; and the *solid pink* curve is the CDF generated with the threshold-aware policy with $\bar{s} = 3.77$.

the (random) tumor state enters the yellow region. In contrast, Figure 9(b) shows that under $d_*^{\bar{s}}$ there is more than one period of MTD drug use for many sample paths.

#### 5S.2 Policies, trajectories, and CDFs for the SR model (additional example)

Except for $d_{\max}$, most of the parameter listed below correspond to those from Table 1 in [8]. We have used these for all numerical experiments in both the main text and SM.

| Symbol | Meaning | Value | Unit |
|---|---|---|---|
| $C$ | Carrying capacity of the Petri dish | $4.8 \times 10^6$ | cells |
| $m$ | Size ratio between $S$ and $R$ cells | 30 | adimensional |
| $g_S$ | intrinsic growth rate of the sensitive | 0.031 | hour$^{-1}$ |
| $g_R$ | intrinsic growth rate of the resistant | 0.026 | hour$^{-1}$ |
| $d$ | drug concentration | Maximum: 3 | nM |
| $\alpha$ | drug efficiency | 0.06 | $(\text{nM} \cdot \text{hour})^{-1}$ |
| $\beta$ | action of sensitive on resistant | $6.25 \times 10^{-7}$ | $(\text{cells} \cdot \text{hour})^{-1}$ |

Similar to Figure 5, we here consider an initial tumor configuration $(q_0, p_0) = (0.55, 0.9)$ inside the yellow (drugs-on) region of $d_\star$.

In Figure 10, we see our threshold-aware policy (pink) improves the $\mathbb{P}(\mathcal{J} \leq 72.75)$ to 67% from 50%. In addition, when the budget is tight, the $\mathbb{P}(\mathcal{J} \leq 60)$ under the deterministic-optimal policy is only around 12%, while our

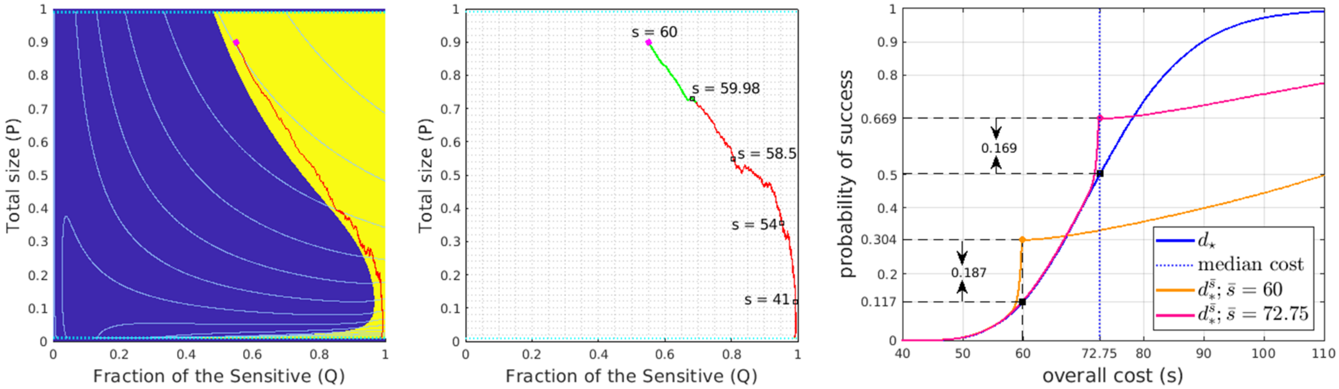


(a) (b) (c)

**Figure 10: Comparison between threshold-aware policies and the deterministic-optimal policy (SR model; Case II).** Starting from an initial state $(q_0, p_0) = (0.55, 0.9)$ (magenta dot): (a) a sample path with cost 56.4 under the deterministic-optimal policy; (b) a sample path starting at $\bar{s} = 60$ with a total cost of 54.3 under the (orange) threshold-aware policy; (c) CDFs of the cumulative cost $\mathcal{J}$ with $10^5$ samples. In (c), the *solid blue* curve is the CDF generated with the deterministic-optimal policy. Its median (*dashed blue* line) is 72.75 while its mean conditioning on success is 73.8. The *solid orange* curve is the CDF generated with the threshold-aware policy with $\bar{s} = 60$; and the *solid pink* curve is the CDF generated with the threshold-aware policy with $\bar{s} = 72.75$.

threshold-aware policy (orange) almost triples this probability to 30%. This is mainly because our threshold-aware policies leverage the inherent competitiveness of the sensitive for a brief period, prescribing drugs once the tumor size slightly diminishes. In contrast, the deterministic-optimal policy will prescribe $d_{\max}$ all the way till remission, yielding a significantly higher cumulative cost.

#### 5S.3 Example 1 with higher volatilities

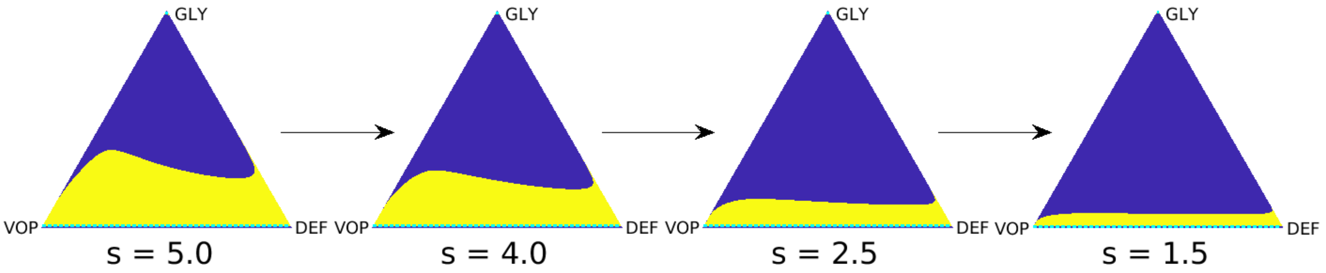


(a)

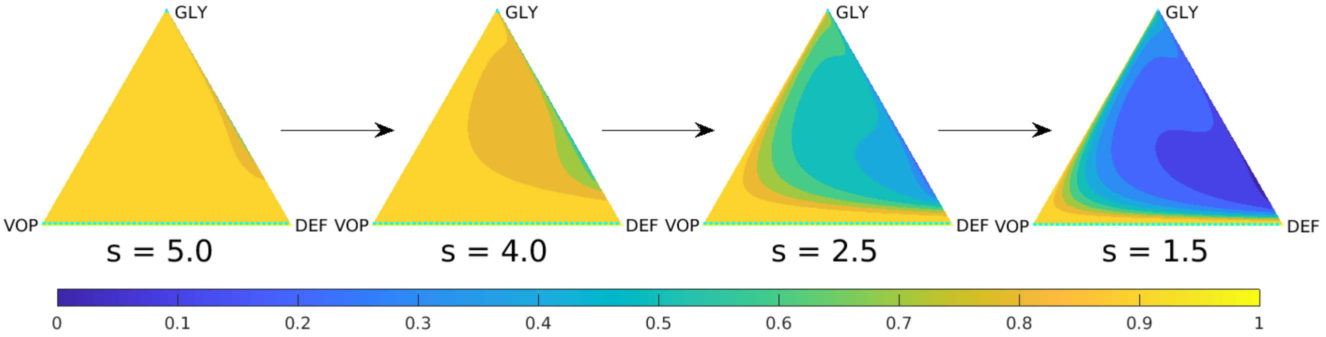


(b)

**Figure 11: Representative slices of the threshold-aware optimal policy (top row) and the corresponding probability of success (bottom row) for the EGT-based model with $\sigma_1 = \sigma_2 = \sigma_3 = 0.5$.** Each triangle represents all possible tumor compositions (proportions of GLY/VOP/DEF cells in the population). Top row shows the policy, which prescribes the optimal instantaneous decisions on drug usage given the indicated remaining budget ($s$) and the current tumor state. Bottom row shows the probability of "stabilization within the budget" if the optimal policy is followed from this time point and onward. Each column corresponds to a specific budget level $s$, which is shown below each triangle. The arrows indicate the natural decrease of the remaining budget while implementing the policy.

The results in this section are obtained by solving (3S.2.5) with $\sigma_1 = \sigma_2 = \sigma_3 = 0.5$. (Other parameter values $d_{\text{max}} = 3, b_a = 2.5, b_v = 2, c = 1, n = 4$ are the same ones as in the main text.)

Figure 11 presents some representative $s$-slices of the optimal policy and their corresponding optimal probability of success. One observes that both Figure 11(a) and Figure 11(b) are significantly different from those computed with $\sigma_1 = \sigma_2 = \sigma_3 = 0.15$; see Figure 3 in the main text.

Since in this case, the random perturbations significantly affect the cancer evolution dynamics, the deterministic portion of Eq. (2S.2.3) (the drift terms) is no longer the dominant force that leads to stabilization. As a consequence, one can see from Figure 11(b) that the closer to the right half of the GLY-DEF-VOP triangle, the higher the optimal probability of stabilization is. Furthermore, unlike with $\sigma_1 = \sigma_2 = \sigma_3 = 0.15$, the drugs-on (yellow) region in Figure 11(a) has just one connected component away from the GLY vertex of the triangle and the drug use is prescribed along the entire stabilization barrier. Consequently, vast majority of samples from simulations shows that once MTD-based therapy is turned on, it stays on until stabilization.

We again compare results from both the deterministic-optimal policy and threshold-aware optimal policies subject to this new stochastic dynamics. We use $(q_0, p_0) = (0.27, 0.4)$ (labeled as a magenta dot in Figure 12) as the initial tumor state, which is inside the drugs-on (yellow) region of the deterministic-optimal policy. Although the resulting sample paths would stay inside the yellow region with high probability (shown in Figure 12(a)), once the random perturbations bring it outside the yellow region, it has 5.3% chance of crossing the failure barrier as shown in Figure 12(b). The corresponding CDF (*solid blue* in Figure 12(e)) shows that the probability of stabilization under $\bar{s} = 3.65$ is only 0.18.

However, if our threshold-aware policy is used instead, the CDF (*solid orange* in Figure 12(e)) shows the maximized probability of stabilization under $\bar{s} = 3.65$ is 0.78, which significantly improves the chance of stabilization by 0.596. The median cost of $\mathcal{J}$ associated with the deterministic-optimal policy is 4.46 (*dashed blue* line in Figure 12(e)). Our threshold-aware policy (*solid pink* in Figure 12(e)) would increase the probability of stabilization under this budget from 0.5 to 0.89. Interestingly, it does not result in a markedly lower probability of success for $s$ values above $\bar{s}$: the pink and blue CDF curves are quite close for large $s$ values. Two representative sample paths under the threshold-aware policy with $\bar{s} = 4.46$ are given in Figures 12(c)&(d). As we see, with higher $\sigma_i$ values the stabilization might be attained without an overall counterclockwise direction of the random trajectory.

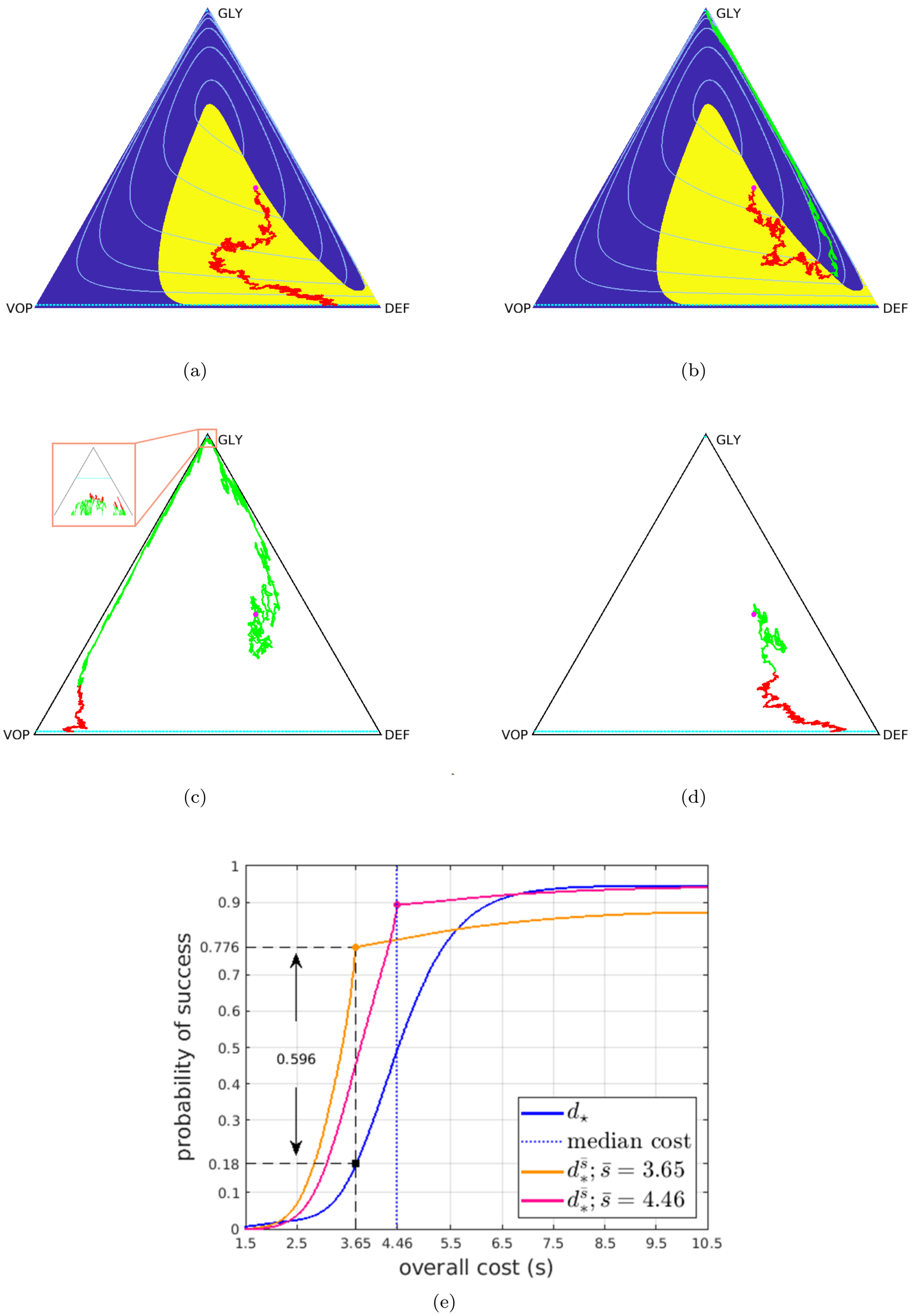


**Figure 12: Comparison between threshold-aware policies and the deterministic-optimal policy (EGT-based model with $\boldsymbol{\sigma_1 = \sigma_2 = \sigma_3 = 0.5}$).** Starting from an initial state $(q_0, p_0) = (0.27, 0.4)$ (magenta dot): (a) a sample path with cost 5.33 under the deterministic-optimal policy; (b) a sample path that leads to failure under the deterministic-optimal policy; (c) & (d) two sample paths starting at $\bar{s} = 4.46$ under the (pink) threshold-aware policy with respective total costs 3.47 and 3.21; (e) CDFs of the cumulative cost $\mathcal{J}$ approximated using $10^5$ random simulations. In (e), the *solid blue* curve is the CDF generated with the deterministic-optimal policy. Its median (*dashed blue* line) is 4.46 while its mean conditioning on success is 4.48. The *solid orange* curve is the CDF generated with the threshold-aware policy with $\bar{s} = 3.65$; and the *solid pink* curve is the CDF generated with the threshold-aware policy with $\bar{s} = 4.46$.
